# Full Bayesian Comparative Phylogeography from Genomic Data

**DOI:** 10.1101/324525

**Authors:** Jamie R. Oaks

## Abstract

A challenge to understanding biological diversification is accounting for community-scale processes that cause multiple, co-distributed lineages to co-speciate. Such processes predict non-independent, temporally clustered divergences across taxa. Approximate-likelihood Bayesian computation (ABC) approaches to inferring such patterns from comparative genetic data are very sensitive to prior assumptions and often biased toward estimating shared divergences. We introduce a full-likelihood Bayesian approach, `ecoevolity`, which takes full advantage of information in genomic data. By analytically integrating over gene trees, we are able to directly calculate the likelihood of the population history from genomic data, and efficiently sample the model-averaged posterior via Markov chain Monte Carlo algorithms. Using simulations, we find that the new method is much more accurate and precise at estimating the number and timing of divergence events across pairs of populations than existing approximate-likelihood methods. Our full Bayesian approach also requires several orders of magnitude less computational time than existing ABC approaches. We find that despite assuming unlinked characters (e.g., unlinked single-nucleotide polymorphisms), the new method performs better if this assumption is violated in order to retain the constant characters of whole linked loci. In fact, retaining constant characters allows the new method to robustly estimate the correct number of divergence events with high posterior probability in the face of character-acquisition biases, which commonly plague loci assembled from reduced-representation genomic libraries. We apply our method to genomic data from four pairs of insular populations of *Gekko* lizards from the Philippines that are *not* expected to have co-diverged. Despite all four pairs diverging very recently, our method strongly supports that they diverged independently, and these results are robust to very disparate prior assumptions.

## 1 Introduction

To understand the distribution of Earth’s biodiversity, we must consider the degree to which environmental changes explain diversity within and among species. A major component of this is understanding how community-scale processes cause co-diversification across evolutionary lineages. Such processes are expected to generate patterns of divergence times that are difficult to explain by lineage-specific processes of diversification. Specifically, finding that divergences are temporally clustered across multiple evolutionary lineages provides compelling evidence that a shared process was responsible for the lineages diverging. For example, the fragmentation of an environment, like an island, forest, or watershed, can cause multiple taxa distributed across that environment to co-diverge over a short period relative to evolutionary timescales (Fig. 1). One way to test the predictions of such processes of diversification is to infer the temporal pattern of divergences across multiple taxa, and determine whether any subsets of the taxa shared the same divergence times.

**Figure 1.**
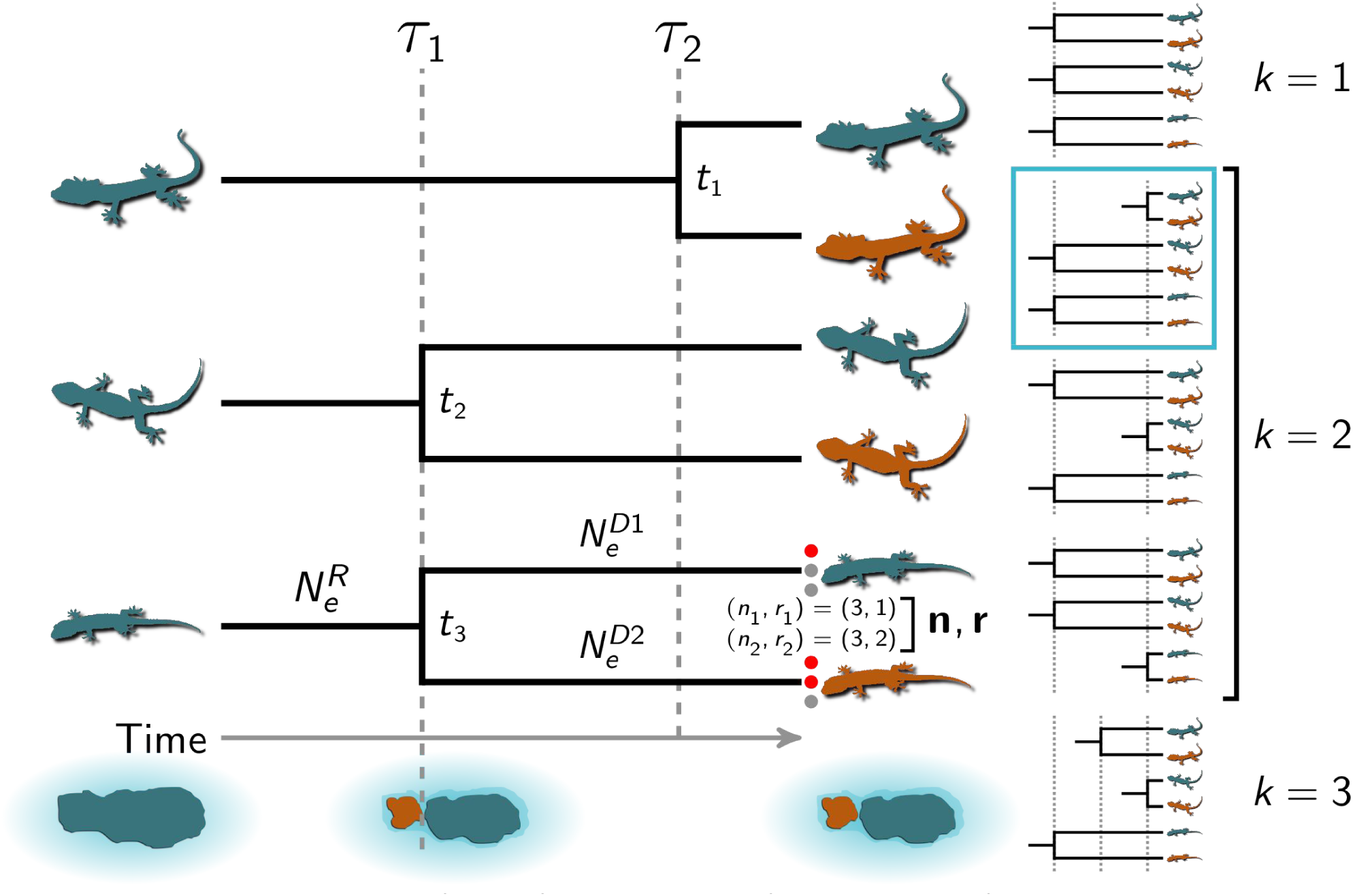
A cartoon depiction of the inference problem for three pairs of insular lizard populations. Three ancestral species of lizards co-occured on a paleo-island that was fragmented into two islands by a rise in sea levels at *τ*_1_. The island fragmentation caused the second and third (from the top) lineages to co-diverge; the first lineage diverged later (at *τ*_2_) via over-water dispersal. The five possible divergence models are shown to the right, with the correct model indicated. The divergence-time parameters (*τ*_1_ and *τ*_2_) and the pair-specific divergence times (*t*_1_, *t*_2_, and *t*_3_) are shown. The third population pair shows the notation used in the text for the biallelic character data (**n**, **r** = (*n*_1_, *r*_1_), (*n*_2_, *r*_2_)) and effective sizes of the ancestral 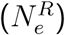 and descendant (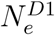 and 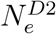) populations. The lizard silhouette for the middle pair is from pixabay.com, and the other two are from phylopic.org; all were licensed under the Creative Commons (CC0) 1.0 Universal Public Domain Dedication.

If researchers are interested in comparing the divergence times among a number of pairs of populations, we can approach this as a problem of model choice: How many divergence events, and what assignment of taxa to those events, best explain the genetic variation within and between the diverged populations of each pair (Fig. 1)? One challenge of this inference problem is the number of possible models. If we have *𝒩* pairs of populations, we would like to assign them to an unknown number of divergence events, *k*, which can range from one to *𝒩*. For a given number of divergence events, the Stirling number of the second kind tells us the number of ways of assigning the taxa to the divergence times (i.e., the number of models with *k* divergence-time parameters):

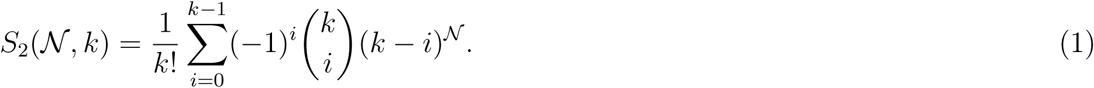

When the number of divergence times is unknown, we need to sum over all possible values of *k* to get the total number of possible divergence models (the Bell number; Bell, 1934)):

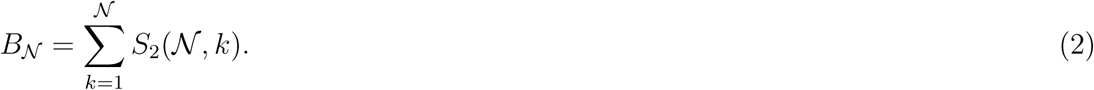

As the number of pairs we wish to compare grows, the prospect of comparing maximum or marginal likelihoods among all possible models quickly becomes daunting. As a result, a Bayesian model-averaging approach is appealing, because it allows the data to determine which models are most relevant.

Methods have been developed to perform this model averaging using approximate-likelihood Bayesian computation (ABC) (Hickerson et al., 2006; Huang et al., 2011; Oaks, 2014). However, these methods often struggle to detect multiple divergence times across pairs of populations (Oaks et al., 2013, 2014) or have little information to update *a priori* expectations (Oaks, 2014). More fundamentally, the loss of information inherent to ABC approaches can prevent them from discriminating among models (Robert et al., 2011; Marin et al., 2013; Green et al., 2015).

One proposed solution is to focus the inference problem on whether or not all pairs diverged at the same time (i.e., *k* = 1 versus *k >* 1) (Hickerson et al., 2014). However, limiting the inference in this way is often not satisfactory, because biogeographers rarely expect that all of the pairs of populations they wish to compare diverged at the same time. Limiting ourselves to the hypothesis of a single shared divergence would not recognize situations where only a subset of taxa co-diverged, or where multiple shared divergences have occurred. The latter is particularly relevant when multiple landscape changes are known to have occurred. More fundamentally, Papadopoulou and Knowles (2016) astutely point out that all of the pairs co-diverging is not the correct null hypothesis. If we wish to test for shared divergences, it is more appropriate to consider all the pairs diverging independently as the null expectation.

Here, our goal is to develop a new Bayesian model-choice approach to this problem that handles many more genetic loci, takes full advantage of the information in those loci, and therefore more reliably estimates the number of divergence events and the assignment of taxa to those events. Our method leverages recent analytical work (Bryant et al., 2012) to efficiently and directly compute the full-likelihood of divergence models from genomic data. By efficiently using all of the information in the data, the new method is faster, more accurate, and more precise than approximate-likelihood methods for estimating shared divergences. We introduce the new method and its assumptions, assess its performance with simulated data, and apply it to genomic data from geckos from the Philippine Islands.

## 2 Methods

### 2.1 The data

We assume we have genetic data from multiple pairs of populations, and our goal is to estimate the time at which the two populations of each pair diverged, and compare these divergence times across the pairs. For each pair of populations that we wish to compare, we assume that we have collected orthologous genetic markers with at most two states. We will refer to these as “biallelic characters,” but note that this includes constant characters (i.e., characters for which all the samples from the two populations share the same state). We follow Bryant et al. (2012) in referring to the two possible states as “red” and “green.” We assume each character is effectively unlinked, i.e., each marker evolved along a gene tree that is independent of the others, conditional on the population history. Examples include well-spaced, single-nucleotide polymorphisms (SNPs) or amplified fragment-length polymorphisms (AFLPs).

For each population and for each marker we sample *n* copies of the locus, *r* of which are copies of the red allele and the remaining *n – r* are copies of the green allele; *r* can range from zero to *n*. Thus, for each population of a pair, and for each locus, we have a count of the total sampled gene copies and how many of those are the red allele.

We will use **n** and **r** to denote allele counts for a locus from both populations of a pair; i.e., **n**, **r** = (*n*_1_, *r*_1_), (*n*_2_, *r*_2_) (Fig. 1 and Table 1). We will also use “character pattern” to refer to **n**, **r**. We will use *D*_*i*_ to denote these counts across all the loci from population pair *i*. In other words, *D*_*i*_ is all the genetic data collected from population pair *i*. Finally, we use **D** to represent the data across all the pairs of populations of which we wish to compare the divergence times. Note, because the pairs are unconnected (Fig. 1 & S1), different characters can be collected for each pair (i.e., characters do not need to be orthologous across the pairs).

**Table 1.**
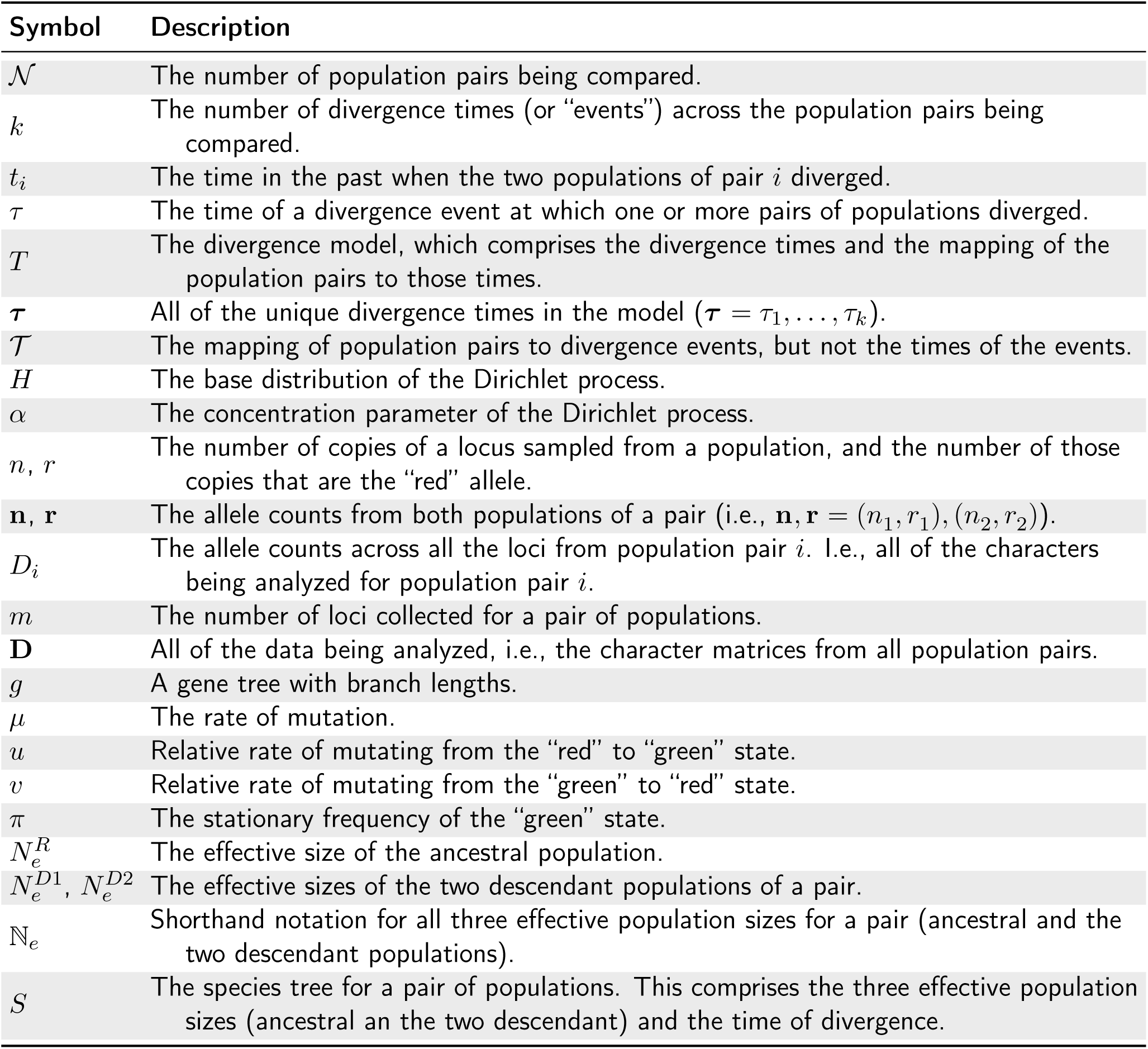
A key to some of the notation used in the text.

### 2.2. The model

#### 2.2.1 The evolution of markers

We assume a finite-sites, continuous-time Markov chain (CTMC) model for the evolution of the biallelic characters along a gene tree with branch lengths, *g*. As the marker evolves along the gene tree, forward in time, there is an instantaneous relative rate *u* of mutating from the red state to the green state, and a corresponding relative rate *v* of mutation from green to red. The stationary frequency of the red and green state is then *v/*(*u* + *v*) and *u/*(*u* + *v*), respectively. Thus, if given the stationary frequency of the green allele, *π*, we can obtain the relative rates of mutation between the two states. We will denote the overall rate of mutation as *µ*. If a mutation rate per site per unit time is given, then branch lengths are in absolute time. Alternatively, if *µ* = 1, the branch lengths of the gene tree are in units of expected substitutions per site. In such a case, for a given pair of populations, the *µ* is redundant, because it can be incorporated into the branch lengths of the gene tree. However, we introduce the notation here, because it will be useful later when we want to allow rate variation among pairs of populations.

#### 2.2.2 The evolution of gene trees

We assume that each marker sampled from a pair of populations evolved within a simple “species” tree with one ancestral root population that diverged into two descendant (terminal) branches at time *t* (Fig. 1). Again, if the *µ* is given, *t* is in units of absolute time; however, if *µ* is set to one, time is in units of expected substitutions per site. We will use ℕ_*e*_ to denote all three effective sizes of a population pair 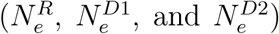. We will also use *S* as shorthand for the species tree, which comprises the population sizes and divergence time of a pair (ℕ_*e*_ and *t*).

#### 2.2.3 The likelihood

Given *µ, π*, and *S*, the probability of the observed data at a locus (**n** and **r**), is the probability of the character pattern given the gene tree multiplied by the probability of the gene tree given the species tree, summed over all possible gene tree topologies and integrated over all possible gene tree branch lengths,

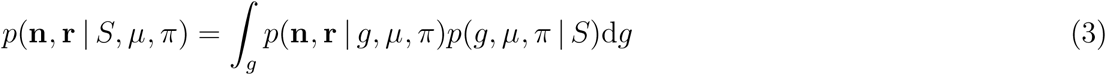

(Felsenstein, 1988; Nielsen and Wakeley, 2001; Rannala and Yang, 2003). We take advantage of the mathematical work of (Bryant et al., 2012) to analytically integrate over all possible gene trees and all possible character substitution histories along those gene trees. This allows us to compute the likelihood of the species tree directly from a biallelic character pattern under a coalescent model, i.e., *p*(**n**, **r|***S, µ, π*). We refer readers to Bryant et al. (2012) for the details of this likelihood and the algorithms to compute it.

Assuming independence among loci (conditional on the species tree), we can calculate the probability of *m* loci given the species tree by simply taking the product over them,

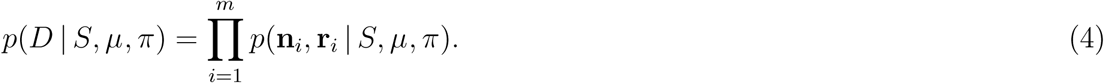

Finally, assuming our *𝒩* pairs are independent, the overall likelihood is simply the product of the likelihood of each pair,

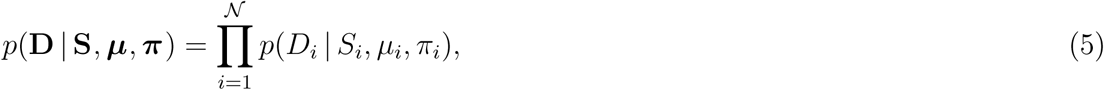

where **D** = *D*_1_, *D*_2_, *…, D*_*N*_, **S** = *S*_1_, *S*_2_, *…, S*_*N*_, ***µ*** = *µ*_1_, *µ*_2_, *…, µ*_*N*_, and ***π*** = *π*_1_, *π*_2_, *…, π*_*N*_.

#### 2.2.4 Correcting for excluded constant characters

If we exclude constant characters and only analyze variable characters, we need to correct the sample space for the excluded constant characters. We can correct the likelihood by simply dividing by the probability of a variable character, which is equal to one minus the probability of a constant character,

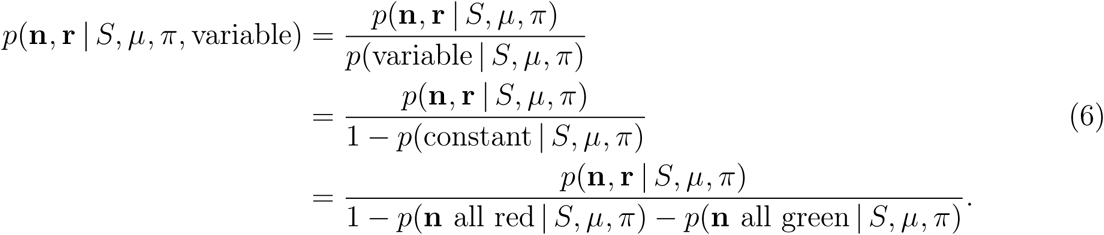

When we take the product over loci to get the probability of all the variable data collected from a pair of populations, we correct each character pattern to allow for different numbers of sampled gene copies among loci,

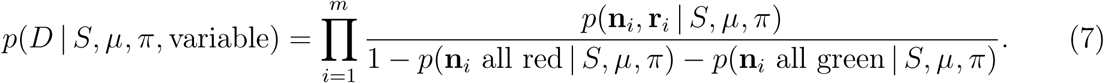

This is a bit different than the correction done in the software SNAPP (Bryant et al., 2012). If we use max(**n**) to denote the maximum number of gene copies sampled from each population, then the correction in SNAPP is

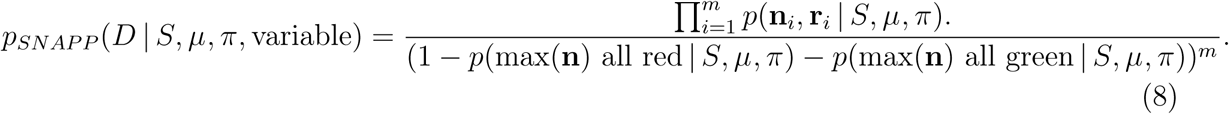

These are equivalent if the same number of samples are collected across all variable loci for each population (i.e., no missing gene copies), but will deviate if fewer copies are sampled for at least one locus. Thus, identical likelihoods between SNAPP and our method should not be expected when analyzing variable-only data.

### 2.3 Bayesian inference

We can obtain a posterior probability distribution by naively plugging the likelihood in Equation 5 into Bayes’ rule,

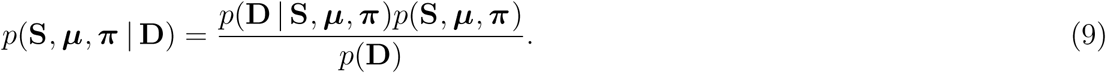

However, this assumes all pairs of populations diverged independently, not allowing us to learn about shared divergence times. What we want to do is relax this assumption and allow pairs to share divergence times.

Let’s use *T* to represent the divergence model, which comprises the divergence times— the number of which (*k*) can range from 1 to *𝒩*—and the mapping of non-overlapping subsets of the population pairs to these *k* divergence times. We will separate out *T* into two components,

1. the partitioning of the *𝒩* population pairs to divergence events, which we will denote as *𝒯*, and
2. the divergence times themselves, ***τ*** = *τ*_1_, *…, τ*_*k*_, the number of which (*k*) is determined by *𝒯*.

We relax the assumption of independent divergence times by treating the number of divergence events and the assignment of population pairs to those events as random variables under a Dirichlet process (Ferguson, 1973; Antoniak, 1974). Specifically, we use the Dirichlet process as a prior on divergence models, *T∼*DP(*H, α*), where *H* is the base distribution of the process and *α* is concentration parameter that controls how clustered the process is. The concentration parameter determines the prior probability of *𝒯* (the partitioning of the population pairs) and the base distribution determines the prior probability of the divergence time of each subset.

Under the Dirichlet process prior, the posterior becomes

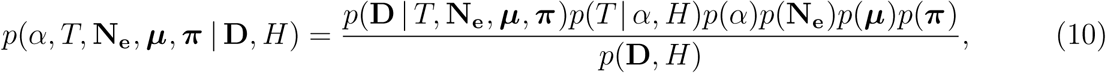

where **N**_**e**_ is the collection of the effective population sizes (ℕ_*e*_) across all of the pairs. By expanding the divergence model (*T*) into the partitioning of the population pairs to divergence events (𝒯) and the times of those events (***τ***), we get

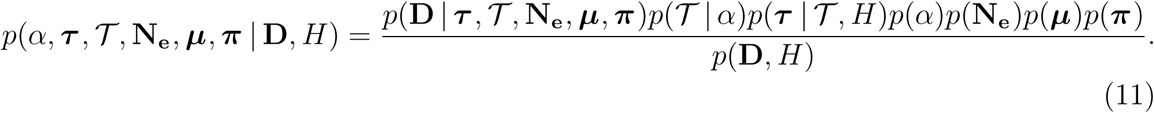

#### 2.3.1 Priors

##### Prior on the concentration parameter

Given a single parameter, *α*, the Dirichlet process determines the prior probability of all the possible ways the *𝒩* pairs of populations can be partitioned to *k* = 1, 2, *… 𝒩* divergence events. Given *α*, the prior probability that two pairs of populations, *i* and *j* (assuming *i ≠ j*), share the same divergence time is

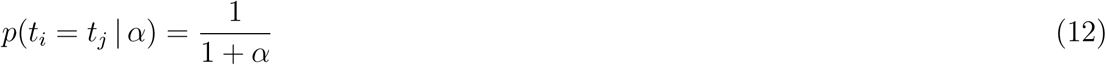

This illustrates that when *α* is small, the process tends to be more clumped, and as it increases, the process tends to favor more independent divergence times. One option is to simply fix the concentration parameter to a particular value, which is likely sufficient when the number of pairs is small. Alternatively, we allow a hierarchical approach to accommodate uncertainty in the concentration parameter by specifying a gamma distribution as a prior on *α* (Escobar and West, 1995; Heath et al., 2011).

##### Prior on the divergence times

Given the partitioning of the pairs to divergence events, we use a gamma distribution for the prior on the time of each event, *τ |𝒯* ∼ Gamma(·,·). This is the base distribution (*H*) of the Dirichlet process.

##### Prior on the effective population sizes

For the two descendant populations of each pair, we use a gamma distribution as the prior on the effective population sizes. For the root population, we use a gamma distribution on the effective population size *relative* to the mean size of the two descendant populations, which we denote as *R*_*N*_*R*. For example, a value of one would mean the root population size is equal to 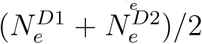. The goal of this approach is to allow more informative priors on the root population size; we often have stronger prior expectations for the relative size of the ancestral population than the absolute size. This is important, because the effective size of the ancestral population is a difficult nuisance parameter to estimate and can be strongly correlated with the divergence time. For example, if the divergence time is so old such that all the gene copies of a locus coalesce within the descendant populations, the locus provides very little information about the size of the ancestral population. As a result, a larger ancestral population and more recent divergence will have a very similar likelihood to a small ancestral population and an older divergence. Thus, placing more prior density on reasonable values of the ancestral population size can help improve the precision of divergence-time estimates.

##### Prior on mutation rates

In the model presented above, for each population pair, the divergence time (*τ*) and mutation rate (*µ*) are inextricably linked. For a single pair of populations, if little is known about the mutation rate, this problem is easily solved by setting it to one (*µ*_1_ = 1) such that time is in units of expected substitutions per site and the effective population sizes are scaled by *µ*. However, what about the second pair of populations for which we wish to compare the divergence time to the first? Because the species trees in our model are disconnected (Fig. 1 & S1), we cannot learn about the relative rates of mutation across the population pairs from the data. As a result, we need strong prior information about the relative rates of mutation across population pairs for this model to work.

If the second pair of populations is closely related to the first, and shares a similar life history, we could assume they share the same mutation rate and set the mutation rate of the second pair to one as well (*µ*_1_ = *µ*_2_ = 1). Alternatively, we could relax that assumption and put a prior on *µ*_2_. However, this should be a strongly informative prior. Placing a weakly informative prior on *µ*_2_ would mean that we can no longer estimate its divergence time relative to the first pair, which is our primary goal. So, while it is possible to incorporate uncertainty in relative mutation rates, it is important to keep in mind that the data cannot inform these parameters, and thus the prior uncertainty in rates will be directly reflected in the posterior of divergence times.

##### Prior on the equilibrium-state frequency

Our method allows for a beta prior to be placed on the frequency of the green allele for each pair of populations, *π*_*i*_ *∼*Beta(·, *·*). However, if using SNP data, we advise fixing the frequency of the red and green states to be equal (i.e., *π* = 0.5). The reason for this is that there is no natural way of re-coding four-state nucleotides to two states, and so the relative transition rates, *u* and *v*, are not biologically meaningful. There will always be arbitrariness associated with how one decides to perform this re-coding, and unless *π* = 0.5, this arbitrariness will affect the likelihood and results. Constraining *π* to 0.5 makes the CTMC model a two-state analog of the “JC69” model (Jukes and Cantor, 1969). However, if the genetic markers are naturally biallelic, the frequencies of the two states can be meaningfully estimated, making the model a two-state general time-reversible model (Tavaré, 1986).

#### 2.3.2 Approximating the posterior with MCMC

We use Markov chain Monte Carlo (MCMC) algorithms to sample (approximately) from the joint posterior in Equation 11. To update the divergence model (*T*) during the chain, we use the Gibbs sampling algorithm (Algorithm 8) of Neal (2000). We also use univariate Metropolis-Hastings algorithms (Metropolis et al., 1953; Hastings, 1970) to update each parameter of the model during the MCMC. To improve mixing of the chain when there are strong correlations between divergence times, effective population sizes, and mutation rates we use multivariate Metropolis-Hastings algorithms. The details of these multivariate moves can be found in Appendix A.

### 2.4 Software implementation

The method outlined above is implemented in the open-source software package, `ecoevolity`, written in the `C++` language. The source code is freely available from https://github.com/phyletica/ecoevolity, and documentation is available at http://phyletica.org/ecoevolity/. The software package is accompanied by an extensive test suite, which, among other aspects, validates that the likelihood code returns the same values as SNAPP (Bryant et al., 2012), and all of our MCMC proposals sample from the expected prior distribution when data are ignored.

The `ecoevolity` package includes four programs:

1. `ecoevolity` for performing Bayesian inference under the model described above.
2. `sumcoevolity` for summarizing posterior samples collected by `ecoevolity` and performing simulations to calculate Bayes factors for all possible numbers of divergence events.
3. `simcoevolity` for simulating biallelic characters under the model described above.
4. `DPprobs` for Monte Carlo approximations of probabilities under the Dirichlet process; this can be useful for choosing a prior on the concentration parameter.

We have also developed a Python package, `pycoevolity`, to help with preprocessing data and summarizing posterior samples collected by `ecoevolity`. This includes assessing MCMC chain stationarity and convergence and plotting posterior distributions. The source code for `pycoevolity` is available at <u>https://github.com/phyletica/pycoevolity</u>.

All of our analyses were performed with Version 0.1 (commit 1d688a3) of the `ecoevolity` software package. The `TimeRootSizeMixer` algorithm implemented in this version of the software only updates one ancestral population size per proposal. In Version 0.2 (commit 884780e), the default behavior is for the `TimeRootSizeMixer` proposal to update the ancestral population size for all other pairs associated with the same divergence time (see above). While this tends to improve mixing slightly, it does not change the results we present here in a meaningful way. Our results can be reproduced exactly with Version 0.1. To help facilitate reproducibility, a detailed history of this project is available at https://github.com/phyletica/ecoevolity-experiments, including all of the data and scripts needed to produce our results.

### 2.5 Analyses of simulated data

#### 2.5.1 Validation analyses

Our first step to validate the new method was to verify that it behaves as expected when the model is correct (i.e., data are simulated and analyzed under the same model). We used the `simcoevolity` tool from the `ecoevolity` package, which simulates data under the model described above. All data were simulated under the following settings:

1. 𝒩 = 3
2. *n* = 10 (i.e., 10 alleles—5 diploid individuals—sampled from each population)
3. *α* = 1.414216, which corresponds with a prior mean of *k* = 2 divergence events
4. *τ ∼* Exponential(mean = 0.01)
5. *π* = 0.5
6. *µ* = 1

We simulated data under five different settings for the effective population sizes. The first setting was an idealized situation where all population sizes were known and equal, 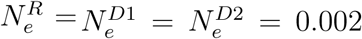. The four remaining scenarios differed in their distribution on the relative effective size of the root population:

1. 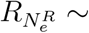 Gamma(shape = 2, mean = 1)
2. 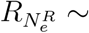 Gamma(shape = 10, mean = 1)
3. 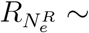 Gamma(shape = 100, mean = 1)
4. 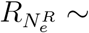 Gamma(shape = 1000, mean = 1)

For these four scenarios, the descendant populations were distributed as Gamma(shape = 5, mean = 0.002). The most difficult nuisance parameter to estimate for a pair of populations is the root population size, which can be correlated with the parameter of interest, the divergence time. Thus, our choice of simulation settings is designed to assess how uncertainty in the root population size affects inference.

Under each of the five scenarios we simulated 500 data sets of 100,000 characters and 500 data sets of 500,000 characters. This includes constant characters; the mean number of variable SNPs was approximately 5,500 and 27,500, respectively. We then analyzed all 5,000 simulated data sets in `ecoevolity` both with and without constant characters included. For all analyses, the prior for each parameter matched the distribution the true value was drawn from when the data were simulated. For analyses where 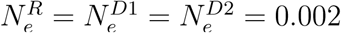, we ran three independent MCMC chains for 37,500 generations, sampling every 25th generation. For all other analyses, we ran the three chains for 75,000 generations, sampling every 50th generation. As a result, we collected 4503 samples for each analysis (1501 samples from each chain, including the initial state).

In order to assess the frequentist behavior of the posterior probabilities of divergence models inferred by `ecoevolity`, we simulated an additional 20,000 data sets of 100,000 characters under the setting where 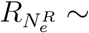 Gamma(shape = 100, mean = 1). All 20,500 data sets were analyzed with `ecoevolity` and binned based on the inferred posterior probability that *k* = 1. The mean posterior probability that *k* = 1 for each bin was plotted against the proportion of data sets within the bin for which the true divergence model was *k* = 1; the latter approximates the true probability that *k* = 1. If the new method is unbiased, in a frequentist sense, the inferred posterior probabilities that *k* = 1 within a bin should approximately equal the proportion of the data sets for which that is true (Huelsenbeck and Rannala, 2004; Oaks et al., 2013; Oaks, 2014).

#### 2.5.2 Assessing the effect of linked characters

The characters of most data sets being collected by high-throughput technologies do not all evolve along independent gene trees. Most consist of many putatively unlinked loci that each comprise sequences of linked nucleotides. For example, “RADseq” and “sequence capture” techniques generate thousands of loci that are approximately 50–300 nucleotides in length. This creates a question when using methods like `ecoevolity` that assume each character is independent: Is it better to violate the assumption of unlinked characters and use all of the data, or throw away much of the data to avoid linked characters?

To better adhere to the unlinked-character assumption, we could retain only a single site per locus. However, this results in a very large loss of data. Furthermore, to try and maximize the informativeness of the retained characters, most researchers retain only one *variable* character per locus. While this can be corrected for (see Equation 7), it still results in the loss of a very informative component of the data: The proportion of variable characters. Before throwing away so much information, we should determine whether it is in our best interest. In other words, does keeping all of the data and violating the assumption of unlinked characters result in better or worse inferences than throwing out much of our data?

To address this question, we simulated data sets composed of loci of linked sites that were 100, 500, and 1000 characters long. The characters for each locus were simulated along the same gene tree (i.e., no intra-locus recombination). Simulated data sets were analyzed with `ecoevolity` in one of three ways: (1) All characters were included, (2) only variable characters were included, and (3) only a maximum of one variable character per locus was included. Only the last option avoids violating the assumption of unlinked characters, but throws out the most data.

For all three locus lengths, we simulated 500 data sets with a total of 100,000 and 500,000 characters. The settings of the simulations performed with `simcoevolity`, and subsequent analyses with `ecoevolity`, correspond with the validation analyses described above where the relative size of the root population was distributed as Gamma(shape = 100, mean = 1). Furthermore, to assess the affect of linked characters on the posterior probabilities of divergence models, we simulated an additional 10,000 data sets with 1,000, 100-character loci (100,000 total characters each). As described above, to assess the frequentist behavior of the inferred posterior probabilities, we binned the results of the analyses of these 10,500 data sets based on the posterior probability that *k* = 1 and plotted the mean of each bin against the approximated true probability that *k* = 1.

#### 2.5.3 Assessing the effect of missing data

The method should be robust to missing data, because it is simply treated as a smaller sample of gene copies from a particular population for a particular locus. Because each character is assumed to have evolved along a coalescent gene tree, the identity of each gene copy within a population does not matter. Thus, some loci having fewer sampled gene copies from some populations should result in more variance in parameter estimates, but is not expected to create bias. To confirm this behavior, we simulated data sets with different probabilities of sampling each gene copy. Specifically, we simulated data sets for which the probability of sampling each gene copy was 90%, 75%, or 50%, which resulted in data sets with approximately 10%, 25%, or 50% missing data. For each sampling probability, we simulated 100 data sets with 500,000 unlinked characters; the settings were the same as described for the validation analyses above where 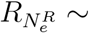 Gamma(shape = 100, mean = 1).

#### 2.5.4 Assessing the effect of biases in character-pattern acquisition

When analyzing the *Gekko* data (see below), we observed large discrepancies in the estimated divergence times depending on whether or not the constant characters were removed from the analysis. This was not observed in the analyses of simulated data, because the likelihood is appropriately corrected for the excluded constant characters. This suggests that there are additional character-pattern acquisition biases in the empirical data, for which are our method cannot correct. Such acquisition biases have been documented during the *de novo* assembly of RADseq loci (Harvey et al., 2015; Linck and Battey, 2017).

The loss of rare alleles during the acquisition and assembly of the data could explain the much larger divergence times estimated from the empirical data when constant characters are removed. After the constant characters, the rare alleles are “next in line” to inform the model that the population divergence was recent. If these patterns are being lost during data acquisition and assembly, and not accounted for in the likelihood calculation, this should create an upward bias in the divergence time estimates.

To explore whether data acquisition bias can explain the discrepancy we observed for the *Gekko* data, we simulated data sets where the probability of sampling singleton character patterns (i.e., one gene copy is different from all the others) was 80%, 60%, and 40%. For each, we simulated and analyzed 100 data sets with 500,000 unlinked characters; the settings were the same as described for the validation analyses above where *R*_*N*_*R ∼* Gamma(shape = 100, mean = 1).

#### 2.5.5 Comparison to ABC methods

We wanted to compare the performance of the new method to the existing approximate-likelihood Bayesian computation (ABC) method `dpp-msbayes` (Oaks, 2014). In order to do this, we had to simulate relatively small data sets that the ABC method could handle in a reasonable amount of time. Accordingly, we simulated data sets with 200 loci, each with 200 linked characters (40,000 total characters). For simulations and analyses of both `ecoevolity` and `dpp-msbayes`, the settings were

1. *𝒩* = 3
2. *n* = 10 (i.e., 10 alleles—5 diploid individuals—sampled from each population)
3. *α* = 1.414216, which corresponds with a prior mean of *k* = 2 divergence events
4. *τ ∼* Gamma(shape = 2, mean = 0.05)
5. *µ* = 1

For `dpp-msbayes`, we placed a Gamma(shape = 5, mean = 0.008) distribution on 4*N*_*e*_*µ* for the ancestral and both descendant populations of each pair. Accordingly, for `ecoevolity`, we used a Gamma(shape = 5, mean = 0.002) distribution on *N*_*e*_*µ* for both descendant populations of each pair. For the relative effective size of the ancestral population in `ecoevolity`, we used a Gamma(shape = 100, mean = 1) distribution; this induces a marginal prior distribution on 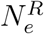 similar to that used for `dpp-msbayes`. For analyses with `dpp-msbayes`, we assumed a Jukes-Cantor model of nucleotide substitution, whereas for the `ecoevolity`, we assumed the two-state equivalent (i.e., *π* = 0.5).

Each method was applied to 500 data sets simulated under its own model. Thus, there were no model violations, except for the new method, for which the assumption of unlinked characters was violated by the 200-character loci. For the analysis of each simulated data set with `ecoevolity`, three independent MCMC chains were run for 75,000 generations, sampling every 50th generation. For the `dpp-msbayes` analyses, 500,000 samples were simulated from the joint prior distribution. To determine which samples to retain for the approximate posterior, for each pair we used the mean of four summary statistics across all the loci:

1. The number of segregating sites (*θ*_*W*_; Watterson, 1975),
2. the average number of pairwise differences across all gene copies (*π*; Nei and Li, 1979),
3. the net number of pairwise differences between the two populations (Equation 25 in Nei and Li, 1979), and
4. the standard deviation in the difference between *π* and *θ*_*W*_ (Tajima, 1989).

After standardizing these statistics, the 2,000 prior samples that were closest to the same statistics calculated from a simulated data set were retained as the approximate posterior.

### 2.6 Empirical application

Previous methods for estimating shared divergence times often over-cluster taxa (Oaks et al., 2013, 2014). Thus, a good empirical test of the new method would be pairs of populations that we expect diverged independently of one another. We analyzed restriction-site-associated sequence (RADseq) data from four pairs of populations of *Gekko* lizards (Table 2). Each pair of populations inhabit two different oceanic islands in the Philippines that were never connected during lower sea levels of glacial periods. Because these islands were never connected, the divergence between the populations of each pair is likely due to over-water dispersal, the timing of which should be idiosyncratic to each pair. We used previous phylogenetic results based on different genetic data (Siler et al., 2012, 2014) to help ensure that the pairs are independent (i.e., they do not overlap each other in the phylogeny of *Gekko*).

**Table 2.** A summary of the data collected from the pairs of *Cyrtodactylus* and *Gekko* populations from the Philippines. Each row represents a pair of populations sampled from two islands that were never connected during low sea levels of glacial periods.

| Species | Island 1 | Island 2 | Sample sizes | # loci | # sites | # variable | # polyallelic |
| --- | --- | --- | --- | --- | --- | --- | --- |
| *G. crombota-rossi* | Babuyan Claro | Calayan | 5 5 | 16,901 | 1,538,408 | 5737 | 50 |
| *G. mindorensis* | Lubang | Luzon | 5 4 | 18,137 | 1,651,186 | 12,092 | 68 |
| *G. mindorensis* | Maestre De Campo | Masbate | 3 3 | 15,993 | 1,455,238 | 11,845 | 27 |
| *G. sp. B-sp. A* | Camiguin Norte | Dalupiri | 5 5 | 15,199 | 1,383,596 | 5612 | 31 |

We analyzed the data with and without the constant characters. Also, there were a small number of sites that had more than two nucleotides represented (Table 2), which cannot be handled directly by our model of biallelic characters. We explored two ways of handling these sites: (1) excluding them, and (2) coding the first nucleotide in the alignment as 0 (“green”), and all other nucleotides for that site as 1 (“red”). Thus, between including/excluding the constant sites and removing/re-coding the polyallelic characters, we analyzed four versions of the RADseq data.

To be conservative in assessing the ability of the new method to distinguish divergence times among the pairs, we set *α* = 0.44, which places 50% of the prior probability on one divergence event (i.e., all four pairs sharing the same divergence). Furthermore, to assess the sensitivity of the results to *α*, we also used *α* = 3.77, which corresponds with a prior mean number of divergence events of three. Other settings that were shared by all analyses of the *Gekko* RADseq data include:

- 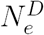 Gamma(shape = 4, mean = 0.004)
- 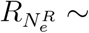 Gamma(shape = 100, mean = 1)
- *π* = 0.5
- *µ* = 1 for all four pairs

The ABC methods of inferring shared divergence events are very sensitive to the prior on divergence times (Oaks et al., 2013; Hickerson et al., 2014; Oaks et al., 2014; Oaks, 2014). To assess whether results of our new method are also sensitive to the prior on divergence times, we analyzed the data sets that included constant characters under the following priors:

1. *τ ∼* Exponential(mean = 0.005)
2. *τ ∼* Exponential(mean = 0.01)
3. *τ ∼* Exponential(mean = 0.05)
4. *τ ∼* Exponential(mean = 0.1)
5. *τ ∼* Exponential(mean = 0.2)

For the two versions of the *Gekko* data that lacked the constant characters, we used the following priors:

1. *τ ∼* Exponential(mean = 0.01)
2. *τ ∼* Exponential(mean = 0.05)
3. *τ ∼* Exponential(mean = 0.1)
4. *τ ∼* Exponential(mean = 0.2)
5. *τ ∼* Exponential(mean = 0.5)

For all analyses, we ran 10 independent MCMC chains for 150,000 generations, sampling every 100th generation. Convergence and mixing of the chains was assessed by the potential scale reduction factor (PSRF; the square root of Equation 1.1 in Brooks and Gelman, 1998) and effective sample size (ESS; Gong and Flegal, 2016) of the log-likelihood and all parameters. We also inspected the chains visually with the program Tracer version 1.6 (Rambaut et al., 2014).

The collection and assembly of the *Gekko* RADseq data are detailed by Oaks et al. (2018). The sequence reads are available on the NCBI Sequence Read Archive (Bioproject PRJNA486413, SRA Study SRP158258) and the assembled data matrices are available in our project repository (<u>https://github.com/phyletica/ecoevolity-experiments</u>).

#### 2.6.1 Empirical comparison to ABC

The *Gekko* data set is much too large to analyze with existing ABC methods for estimating co-divergences. In order to compare the results of the new full-likelihood method, `ecoevolity`, to the ABC method, `dpp-msbayes`, we randomly sampled (without replacement) 200 loci from three of the pairs of *Gekko* populations. The prior settings we used for the `ecoevolity` analysis of this reduced data set was:

- *α* = 1.414216, which corresponds with a prior mean number of divergence events of two
- *τ ∼* Exponential(mean = 0.1)
- 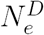 Gamma(shape = 5, mean = 0.002)
- 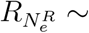 Gamma(shape = 100, mean = 1)
- *π* = 0.5
- *µ* = 1 for all three pairs

We used the same settings for the `dpp-msbayes` analysis, except to account for the different parameterization of effective population sizes, we used a Gamma(shape = 5, mean = 0.008) prior on 4*N*_*e*_*µ* for the ancestral and descendant populations.

For the `ecoevolity` analysis, we ran five independent MCMC chains for 12,000 generations, sampling every 10th generation. We assessed convergence and mixing using the same methods as we did for the analyses of the full *Gekko* data set, described above. For the `dpp-msbayes` analysis, we simulated 500,000 samples from the joint prior, and to get a sample from the approximate posterior, we retained the 5,000 samples with summary statistics most similar to those calculated from the reduced *Gekko* data set. For each pair, we used two summary statistics: The average number of pairwise differences across all gene copies (*π*) and between the gene copies from the two populations (*π*_*b*_) (Nei and Li, 1979). For each pair, we used the mean of both statistics across the 200 loci. The results from simulated data demonstrate that additional statistics that summarize information about effective population sizes are not informing the ABC method (see below).

## 3 Results

### 3.1 Analyses of simulated data

#### 3.1.1 Validation analyses

When there is no model misspecification, our new method has the desired frequentist behavior wherein 95% of the time the true value of a parameter falls within the 95% credible interval. We see this for divergence times (Fig. 2) and the effective sizes of descendant (Fig. S2) and ancestral (Fig. S3) populations. Our results also show that the estimated posterior probability of the single divergence model (*k* = 1) mirrors the probability that the model is correct (Fig. 3). We see the same behaviors whether or not the constant characters are excluded, demonstrating that our likelihood correction for excluded constant characters is working correctly (Equation 7).

**Figure 2.**
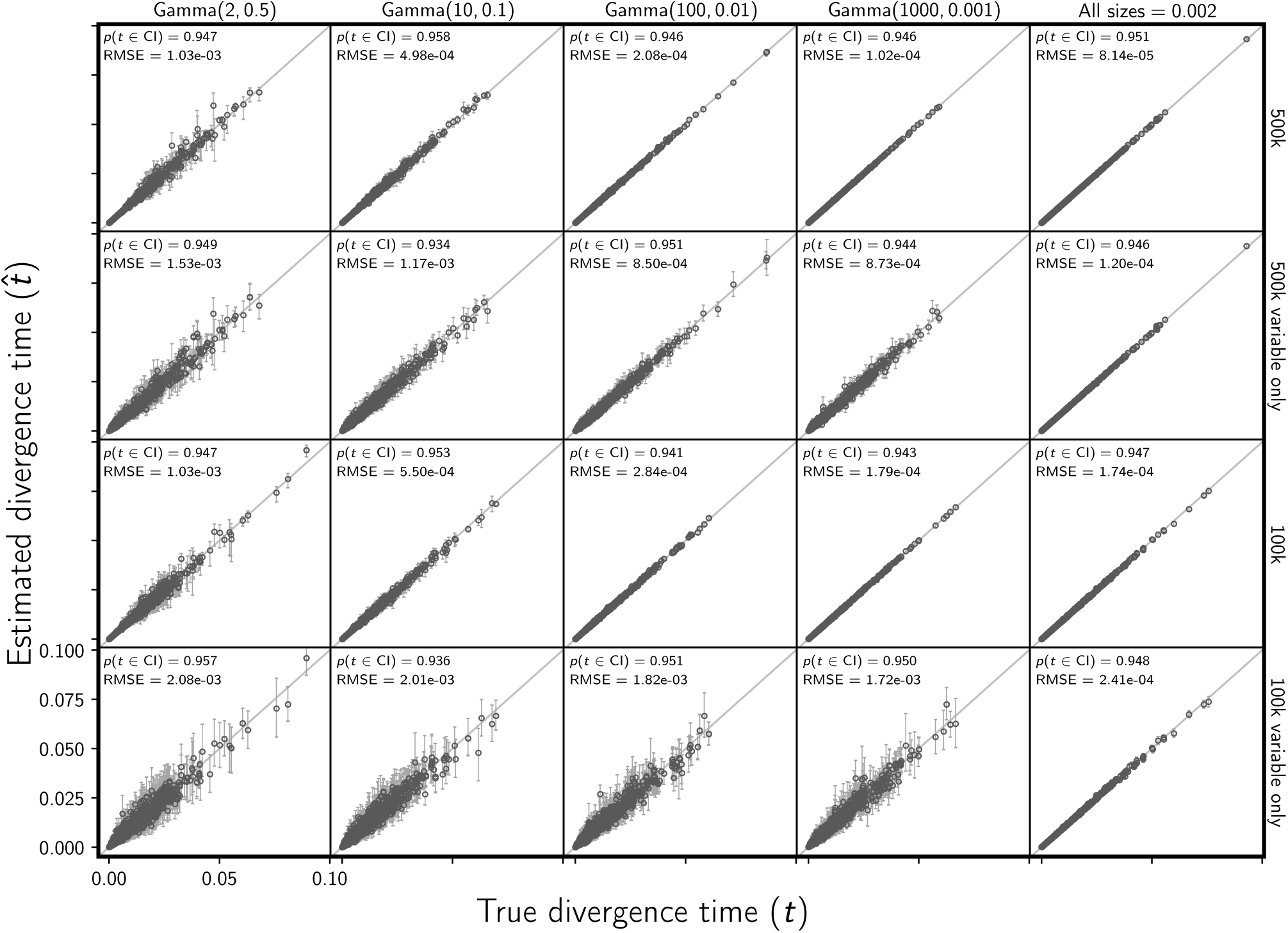
The accuracy and precision of divergence time estimates, in units of expected subsitutions per site, when data are simulated and analyzed under the same model (i.e., no model misspecification). The first four columns show the results from different distributions on the relative effective size of the ancestral population, decreasing in variance from left to right. The fifth column shows results when the effective size (*N*_*e*_*µ*) of all populations is fixed to 0.002. For the first two and last two rows, the simulated character matrix for each population had 500,000 and 100,000 characters, respectively. The first and third rows show the results of analyses using all characters, whereas the second and fourth rows show the results when only variable characters are used. Each plotted circle and associated error bars represent the posterior mean and 95% credible interval for the time that a pair of populations diverged. Each plot consists of 1500 estimates—500 simulated data sets, each with three pairs of populations. For each plot, the root-mean-square error (RMSE) and the proportion of estimates for which the 95% credible interval contained the true value—*p*(*t∈*CI)—is given. We generated the plot using matplotlib Version 2.0.0 (Hunter, 2007).

**Figure 3.**
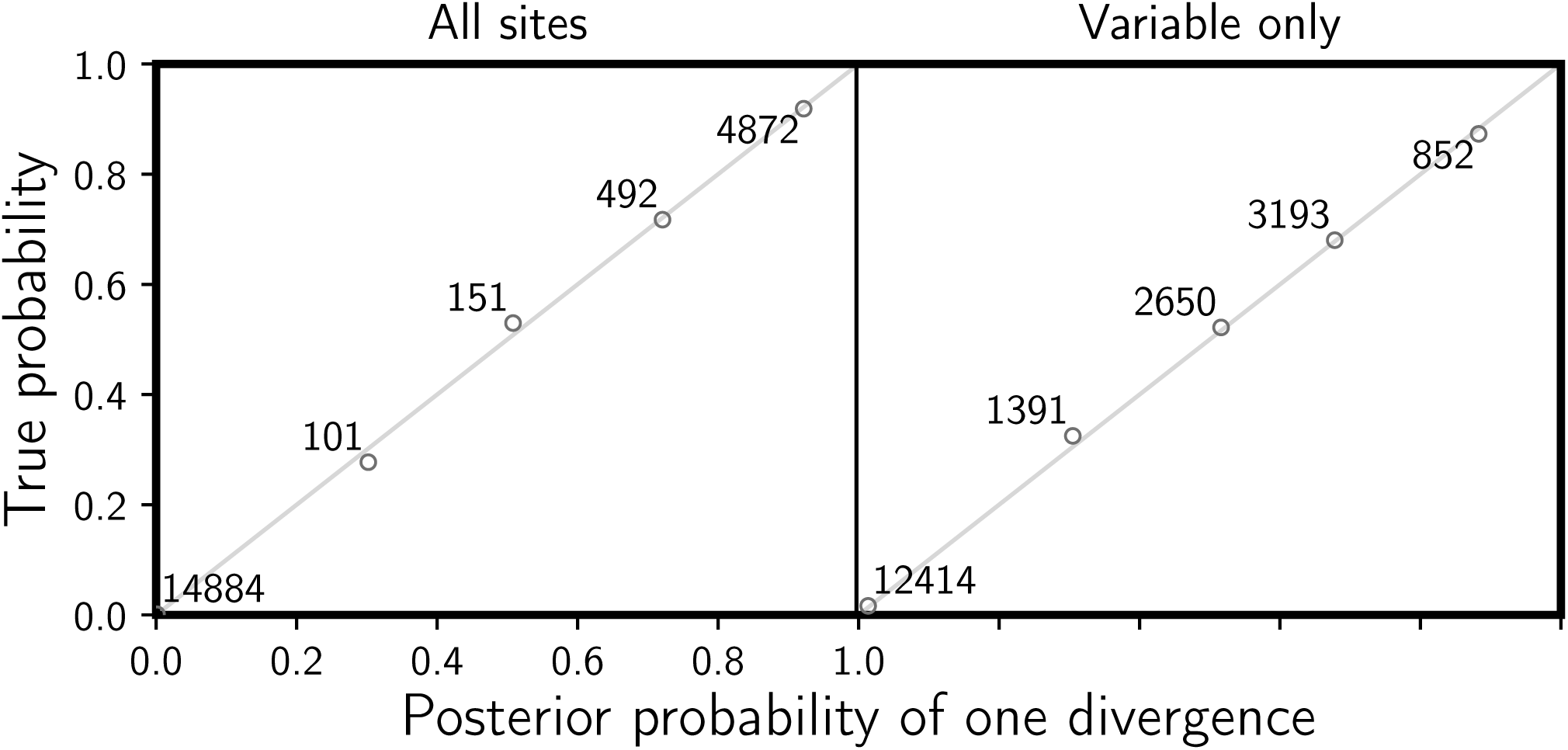
Assessing frequentist behavior of divergence-model posterior probabilities when there is no model misspecification. 20,500 data sets were simulated and analyzed under the same model and assigned to bins of width 0.2 based on the estimated posterior probability of a single, shared divergence event. The mean posterior probability of each bin is plotted against the proportion of data sets in the bin for which a single, shared divergence is the true model. The number of data sets within each bin is provided next to the corresponding plotted point. The left plot shows the results when all characters are analyzed, and the right plot shows the results when only the variable characters are analyzed. All simulated data sets had three pairs of populations, each with 100,000 characters. We generated the plot using matplotlib Version 2.0.0 (Hunter, 2007).

As expected, the precision of divergence time and population size estimates is greater when the constant characters are included and when there is greater prior information about the ancestral population size (Fig. 2, S2, & S3). The increase in precision associated with the fivefold increase in the number of sampled characters (100k to 500k) is relatively modest (Fig. 2, S2, & S3). Retaining the constant characters results in a much larger increase in precision than collecting five times more characters.

The true model and number of divergence events is included in the 95% credible set greater than 97% of the time for all the simulation conditions (Fig. 4 & S4). The frequency at which the correct number of events has the largest posterior probability, and the median posterior probability of the correct number of events, increases when constant characters are retained and as prior information about the ancestral population size increases (Fig. 4). We see the same patterns for inferring the correct divergence model (Fig. S4). As with the parameter estimates, the performance increase associated with the increase from 100k to 500k sampled characters is moderate; retaining the constant characters has a much larger effect (Fig. 4 & S4). When constant characters are used, the median posterior probability of the correct number of divergence events is high (over 0.89 for all simulation conditions; Fig. 4). Likewise, the median posterior probability of the correct divergence model is greater than 0.887 across all simulation conditions when constant characters are used (Fig. S4).

**Figure 4.**
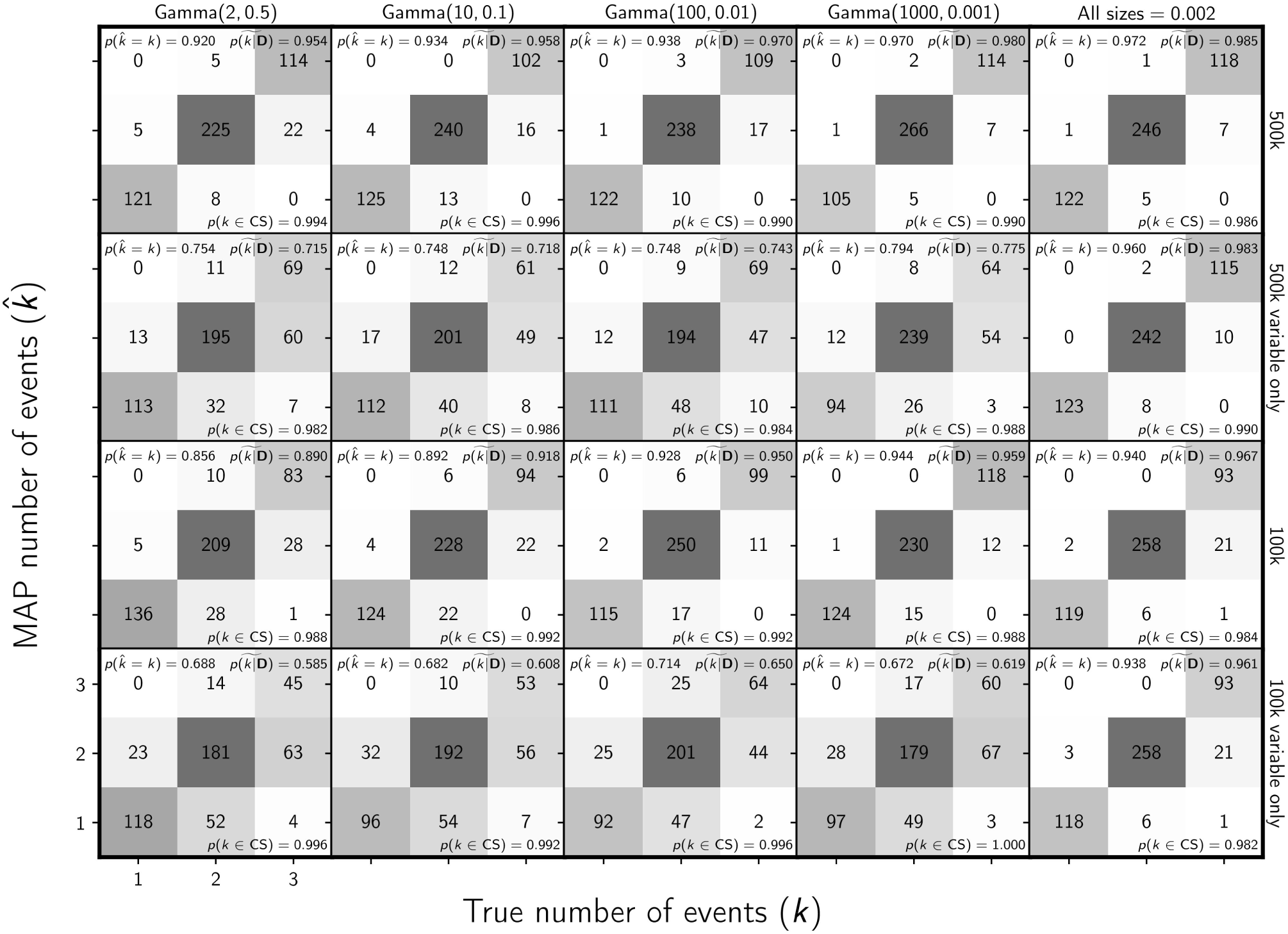
The ability of the new method to estimate the number of divergence events when data are simulated and analyzed under the same model (i.e., no model misspecification). The first four columns show the results from different distributions on the relative effective size of the ancestral population, decreasing in variance from left to right. The fifth column shows results when the effective size (*N*_*e*_*µ*) of all populations is fixed to 0.002. For the first two and last two rows, the simulated character matrix for each population had 500,000 and 100,000 characters, respectively. The first and third rows show the results of analyses using all characters, whereas the second and fourth rows show the results when only variable characters are used. Each plot shows the results of the analyses of 500 simulated data sets, each with three population pairs; the number of data sets that fall within each possible cell of true versus estimated numbers of events is shown, and cells with more data sets are shaded darker. The estimates are based on the number of events with the maximum *a posteriori* (MAP) probability. For each plot, the proportion of data sets for which the number of events with the largest posterior probability matched the true number of events—*p*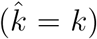—is shown in the upper left corner, the median posterior probability of the correct number of events across all data sets—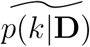—is shown in the upper right corner, and the proportion of data sets for which the true number of events was included in the 95% credible set—*p*(*k∈*CS)—is shown in the lower right. We generated the plot using matplotlib Version 2.0.0 (Hunter, 2007).

For the data sets simulated with 100,000 and 500,000 characters, the number of variable characters ranged from 515–21,676 and 4,670–105,373, respectively, with an average of approximately 5,500 and 27,500 variable characters, respectively (Fig. S5 & S6). As expected, the variance in the number variable characters increases with the variance in the prior distribution of the relative effective size of the root population (Fig. S5 & S6).

The MCMC chains for all analyses converged very quickly; we conservatively removed the first 401 samples, resulting in 3300 samples from the posterior (1100 samples from three chains) for each analysis. To assess convergence and mixing, we plotted histograms of the potential scale reduction factor across the three independent chains and the effective sample size for the log-likelihood and divergence times (Fig. S7, S8, S9, & S10). Mixing was poorer when there was more prior uncertainty in the root population size (Fig. S9 & S10). However, given the expected frequentist behavior for how often the true parameter values were contained within the 95% confidence intervals (Fig. 2, S2, & S3), and the weak relationship between the ESS and estimation error (Fig. S11), we do not expect MCMC mixing had a large effect on our simulation results under the most extreme levels of uncertainty in the root population size that we simulated.

#### 3.1.2 Assessing the effect of linked characters

The accuracy of divergence time estimates did not appear to be affected by the model violation of linked characters (Fig. 5 & S12). However, as the length of loci increases, we do see an underestimation of posterior uncertainty (i.e., the true divergence time is contained within the 95% credible interval less frequently than 95% of the time; Fig. 5 & S12). This makes sense given that there is less coalescent variation in the data than the model expects if all the characters had evolved along independent gene trees. Importantly, this effect of underestimating posterior uncertainty is small for data sets with 100bp loci, suggesting this violation of the model has little impact for high-throughput data sets with short loci, like those collected via RADseq. As expected, analyzing only one variable site per locus removes this underestimation of posterior uncertainty (see the last row of Fig. 5 & S12), but at a large cost of much greater posterior uncertainty in parameter estimates due to the loss of data. We see the same behavior for estimating the effective sizes of the ancestral and descendant populations (Fig. S13–16).

**Figure 5.**
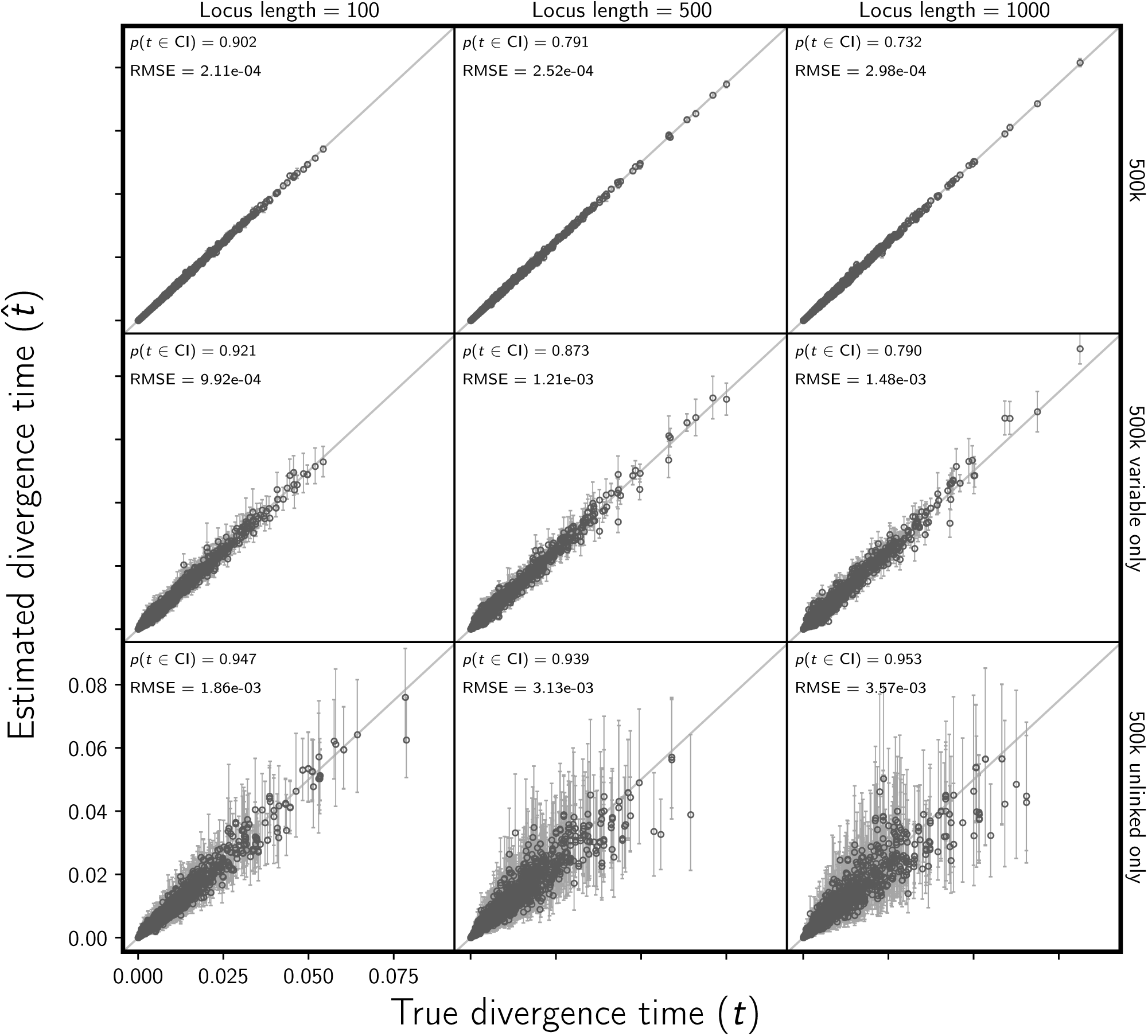
Assessing the effect of linked sites on the the accuracy and precision of divergence time estimates (in units of expected subsitutions per site). The columns, from left to right, show the results when loci are simulated with 100, 500, and 1000 linked sites. For each simulated data set, each of three population pairs has 500,000 sites total. The rows show the results when (top) all sites, (middle) all variable sites, and (bottom) at most one variable site per locus are analyzed. For each plot, the root-mean-square error (RMSE) and the proportion of estimates for which the 95% credible interval contained the true value—*p*(*t∈*CI)—is given. We generated the plot using matplotlib Version 2.0.0 (Hunter, 2007).

The cost of removing data to avoid violating the assumption of unlinked characters is also very pronounced for estimating the divergence model and number of divergence events. The method better estimates the correct model and number of events, both with much higher posterior probability, when the constant characters are retained (Fig. 6, S17, S18, & S19). The median posterior probability of both the correct number of divergence events and the correct model is over 0.95 for all 500k-character data sets, even when loci were 1000bp long (Fig. 6 & S17). However, our results show that linked characters do introduce bias in the estimated posterior probability of the one divergence model (*k* = 1) (Fig. 7). However, the bias is moderate and makes the method conservative in the sense that it tends to underestimate the probability of shared divergence (Fig. 7).

**Figure 6.**
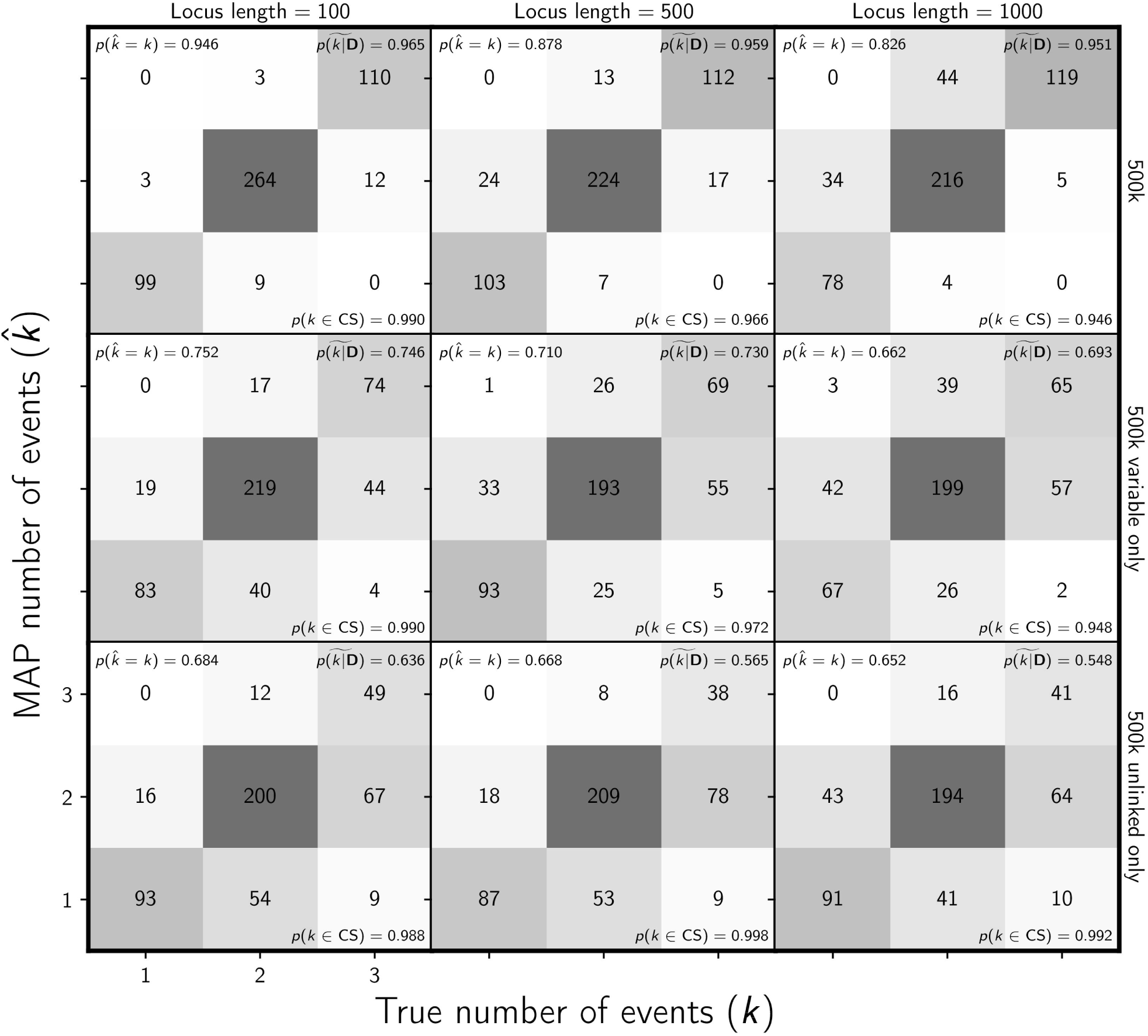
Assessing the effect of linked sites on the ability of the new method to estimate the number of divergence events. The columns, from left to right, show the results when loci are simulated with 100, 500, and 1000 linked sites. For each simulated data set, each of three population pairs has 500,000 sites total. The rows show the results when (top) all sites, (middle) all variable sites, and (bottom) at most one variable site per locus are analyzed. The number of data sets that fall within each possible cell of true versus estimated numbers of events is shown, and cells with more data sets are shaded darker. The estimates are based on the number of events with the maximum *a posteriori* (MAP) probability. For each plot, the proportion of data sets for which the number of events with the largest posterior probability matched the true number of events—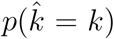—is shown in the upper left corner, the median posterior probability of the correct number of events across all data sets—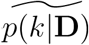—is shown in the upper right corner, and the proportion of data sets for which the true number of events was included in the 95% credible set—*p*(*k∈*CS)—is shown in the lower right. We generated the plot using matplotlib Version 2.0.0 (Hunter, 2007).

**Figure 7.**
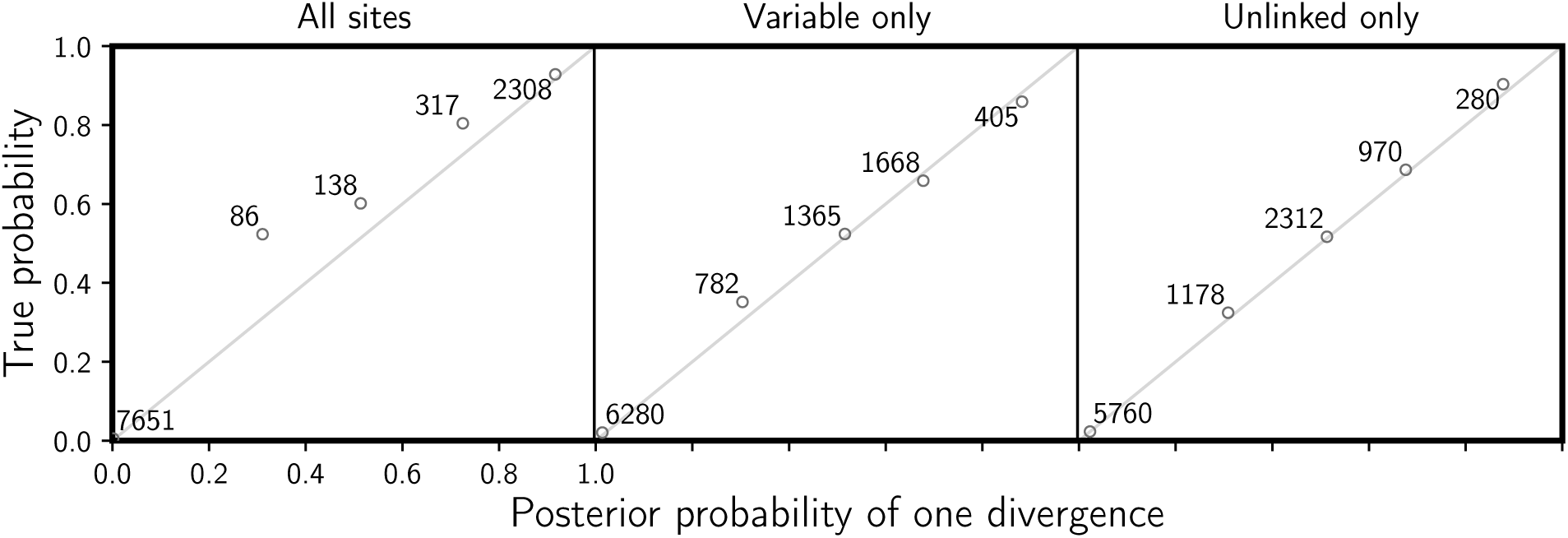
Assessing the effect of linked sites on the frequentist behavior of divergence-model posterior probabilities. 10,500 data sets were simulated such that each of three population pairs has 1000 loci, each with 100 linked sites (100,000 sites total). Each simulated data set is assigned to a bin of width 0.2 based on the estimated posterior probability of a single, shared divergence event. The mean posterior probability of each bin is plotted against the proportion of data sets in the bin for which a single, shared divergence is the true model. The number of data sets within each bin is provided next to the corresponding plotted point. The plots show the results when (left) all characters, (middle) all variable characters, and (right) at most one variable character per locus is analyzed. We generated the plot using matplotlib Version 2.0.0 (Hunter, 2007).

For simulated data sets with loci of length 100, 500, and 1000 base pairs, there were an average of 5.4, 27.1, and 54.1 variable characters per locus, respectively. As expected, the number of variable characters per 100k and 500k data set was very similar to the simulated unlinked-character data sets, with an average of about 5,500 variable characters per 100k data set (Fig. S20) and 27,100 variable characters per 500k data set (Fig. S21). When at most one variable character is sampled per locus, the number of remaining characters is usually close or equal to the number of loci; 1000, 200, and 100 characters for the 100k data sets with 100, 500, and 1000 bp loci, respectively, and 5000, 1000, and 500 characters for 500k data sets with 100, 500, and 1000 bp loci, respectively (Fig. S20 & S21).

#### 3.1.3 Assessing the effect of missing data

As predicted by coalescent theory, our results show that random missing data has little effect on the performance of the method with respect to estimating divergence times (Fig. 8), effective population sizes (Fig. S22 & S23), the number of divergence events (Fig. 9), or the divergence model (Fig. S24).

**Figure 8.**
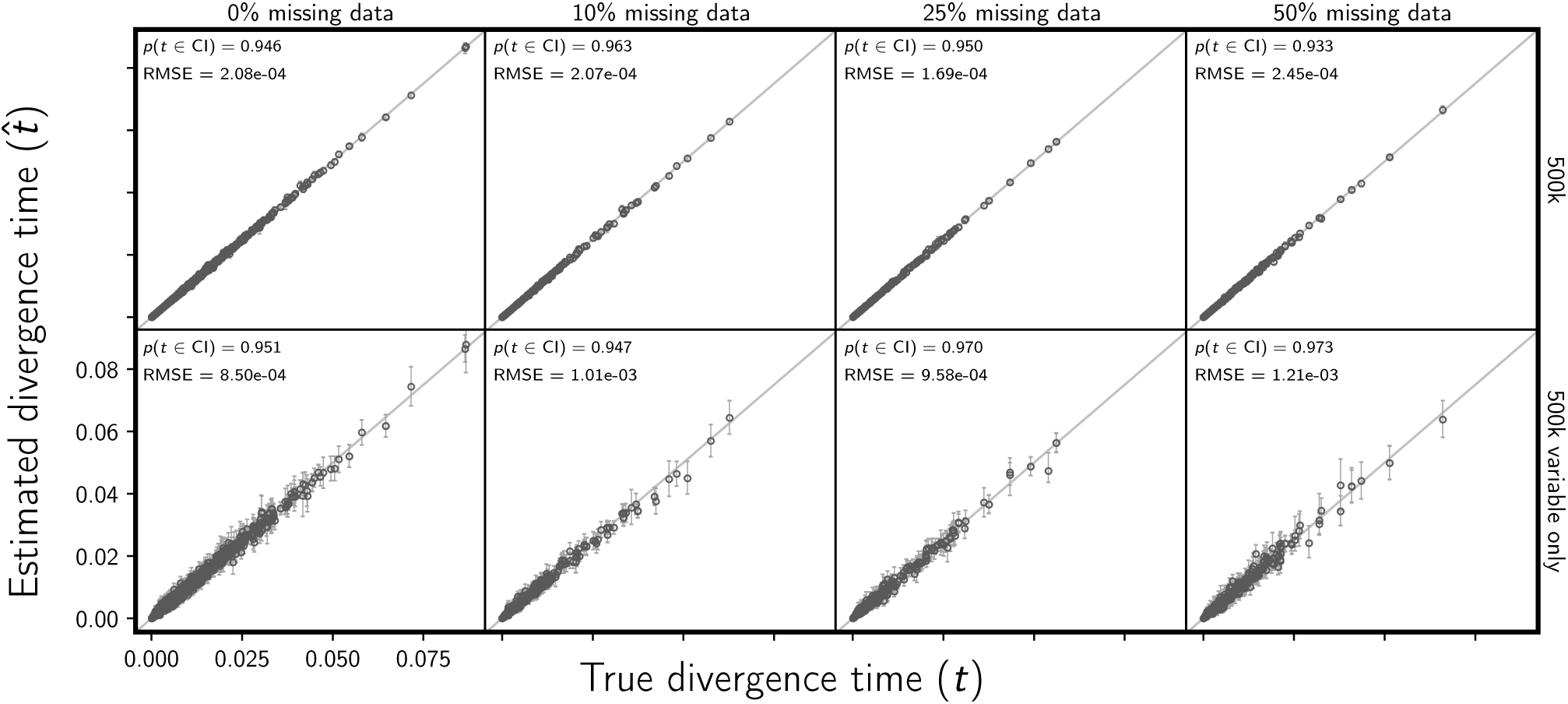
Assessing the effect of missing data on the the accuracy and precision of divergence time estimates (in units of expected subsitutions per site). The columns, from left to right, show the results when each simulated 500,000-character matrix has approximately 0%, 10%, 25%, and 50% missing cells. For comparison, the first column shows the results of the 500 data sets from Figure 2; the remaining columns show the results of 100 data sets. The rows show the results when (top) all sites and (bottom) only variable sites are analyzed. For each plot, the root-mean-square error (RMSE) and the proportion of estimates for which the 95% credible interval contained the true value—*p*(*t∈*CI)—is given. All simulated data sets had three pairs of populations. We generated the plot using matplotlib Version 2.0.0 (Hunter, 2007).

**Figure 9.**
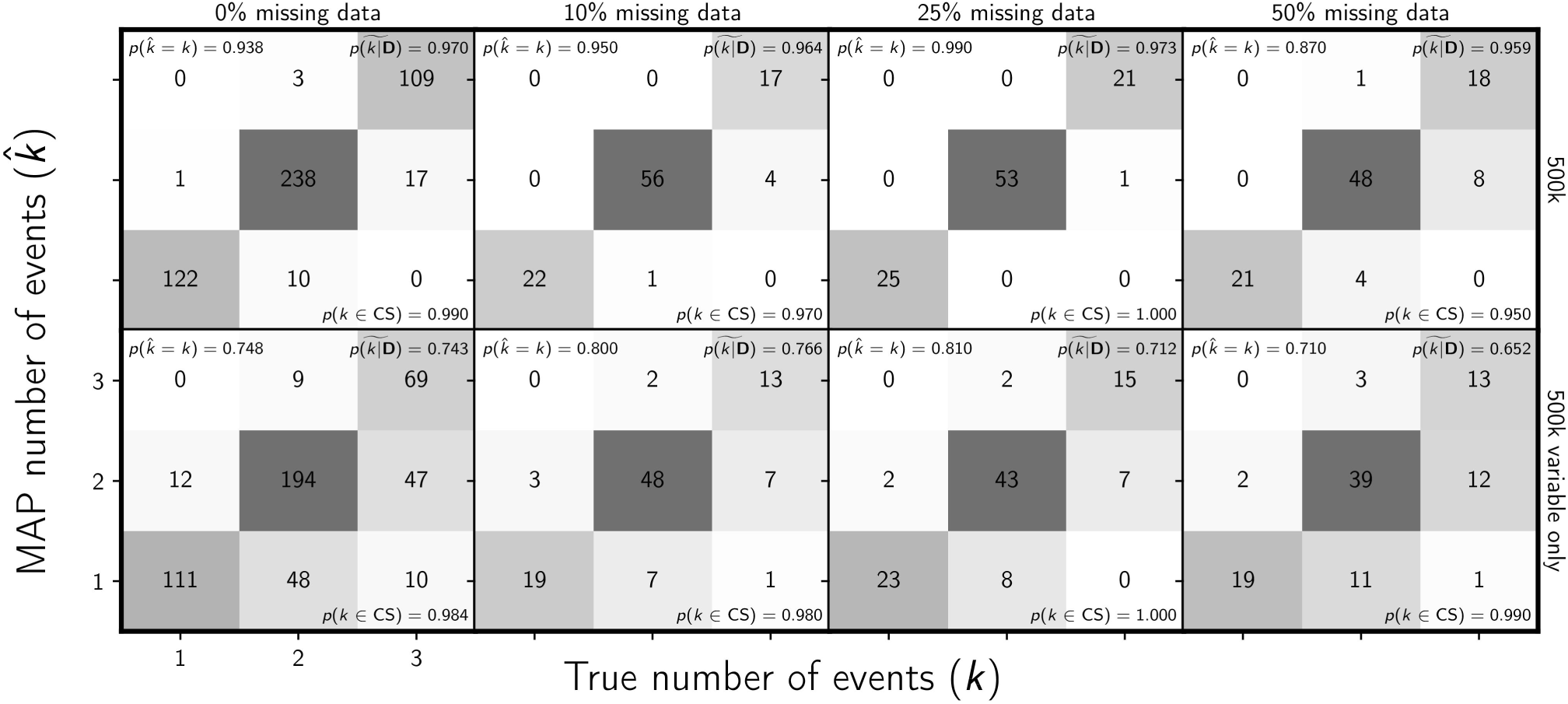
Assessing the effect of missing data on the ability of the new method to estimate the number of divergence events. The columns, from left to right, show the results when each simulated 500,000-character matrix has approximately 0%, 10%, 25%, and 50% missing cells. For comparison, the first column shows the results of the 500 data sets from Figure 4; the remaining columns show the results of 100 data sets. The rows show the results when (top) all sites and (bottom) only variable sites are analyzed. The number of data sets that fall within each possible cell of true versus estimated numbers of events is shown, and cells with more data sets are shaded darker. The estimates are based on the number of events with the maximum *a posteriori* (MAP) probability. For each plot, the proportion of data sets for which the number of events with the largest posterior probability matched the true number of events—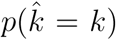—is shown in the upper left corner, the median posterior probability of the correct number of events across all data sets—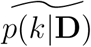—is shown in the upper right corner, and the proportion of data sets for which the true number of events was included in the 95% credible set—*p*(*k∈*CS)—is shown in the lower right. All simulated data sets had three pairs of populations. We generated the plot using matplotlib Version 2.0.0 (Hunter, 2007).

#### 3.1.4 Assessing the effect of biases in character-pattern acquisition

Biased character acquisition against singleton character patterns does create bias in estimates of divergence times (Fig. 10) and population sizes (Fig. S25 & S26), and the bias increases as the probability of missing a character with a singleton pattern increases. Notably, this bias is smaller when the constant characters are retained in the data set (Fig. 10, S25 & S26).

**Figure 10.**
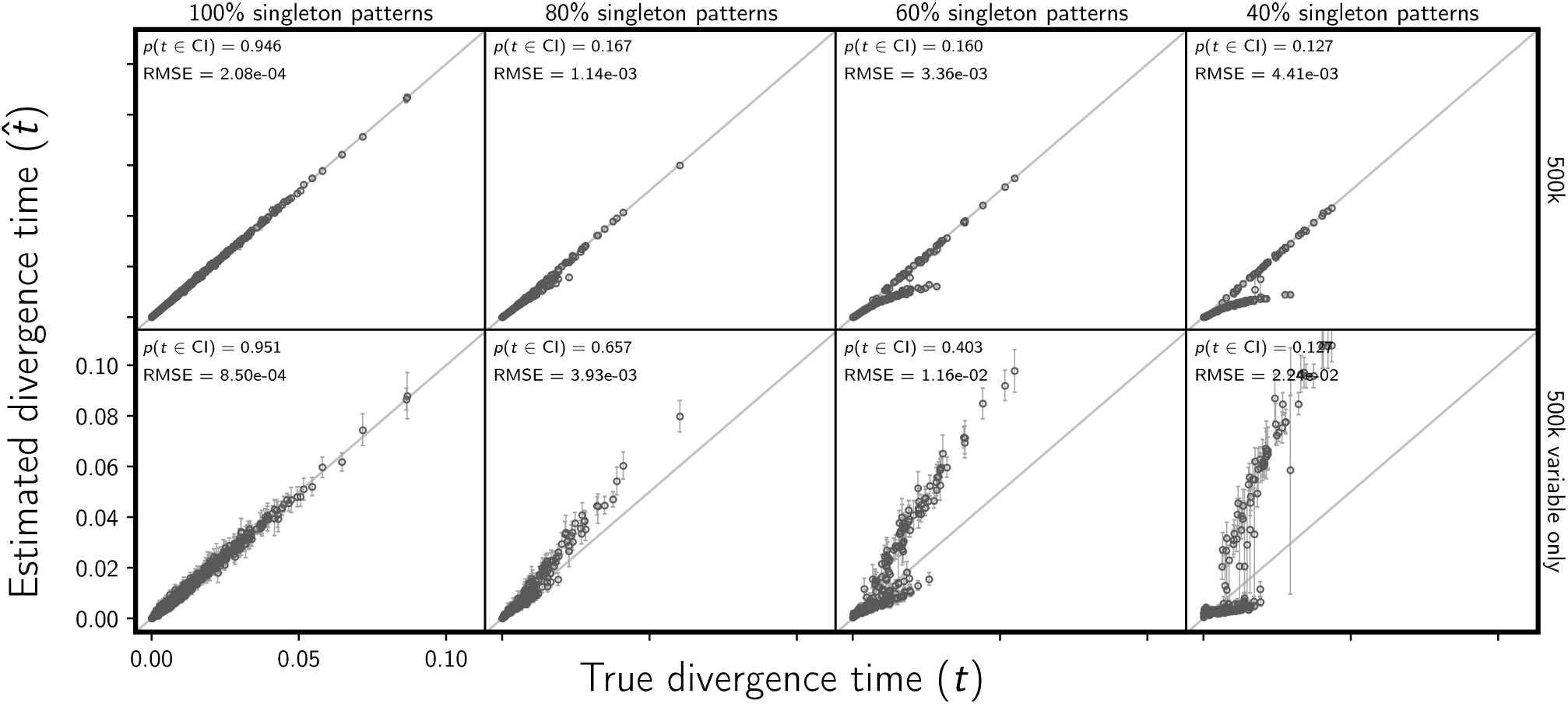
Assessing the effect of an acquisition bias against rare allele patterns on the accuracy and precision of divergence time estimates (in units of expected subsitutions per site). The columns, from left to right, show the results when each simulated 500,000-character data set has a probability of 100%, 80%, 60%, and 40% of sampling each simulated singleton pattern. E.g., each character matrix analyzed in the far right column is missing approximately 60% of characters where all but one gene copy has the same allele. For comparison, the first column shows the results of the 500 data sets from Figure 2; the remaining columns show the results of 100 data sets. The rows show the results when (top) all sites and (bottom) only variable sites are analyzed. For each plot, the root-mean-square error (RMSE) and the proportion of estimates for which the 95% credible interval contained the true value—*p*(*t∈*CI)—is given. All simulated data sets had three pairs of populations. We generated the plot using matplotlib Version 2.0.0 (Hunter, 2007).

However, in the face of data-acquisition bias, the method still estimates the number of divergence events and the divergence model well, especially when constant characters are used (Fig. 11 & S27). Even when the probability of sampling a character with a singleton pattern is 0.4, the median posterior probability of the correct number of divergence events is 0.948 (Fig. 11), and the median posterior probability of the correct model is 0.945 (Fig. S27).

**Figure 11.**
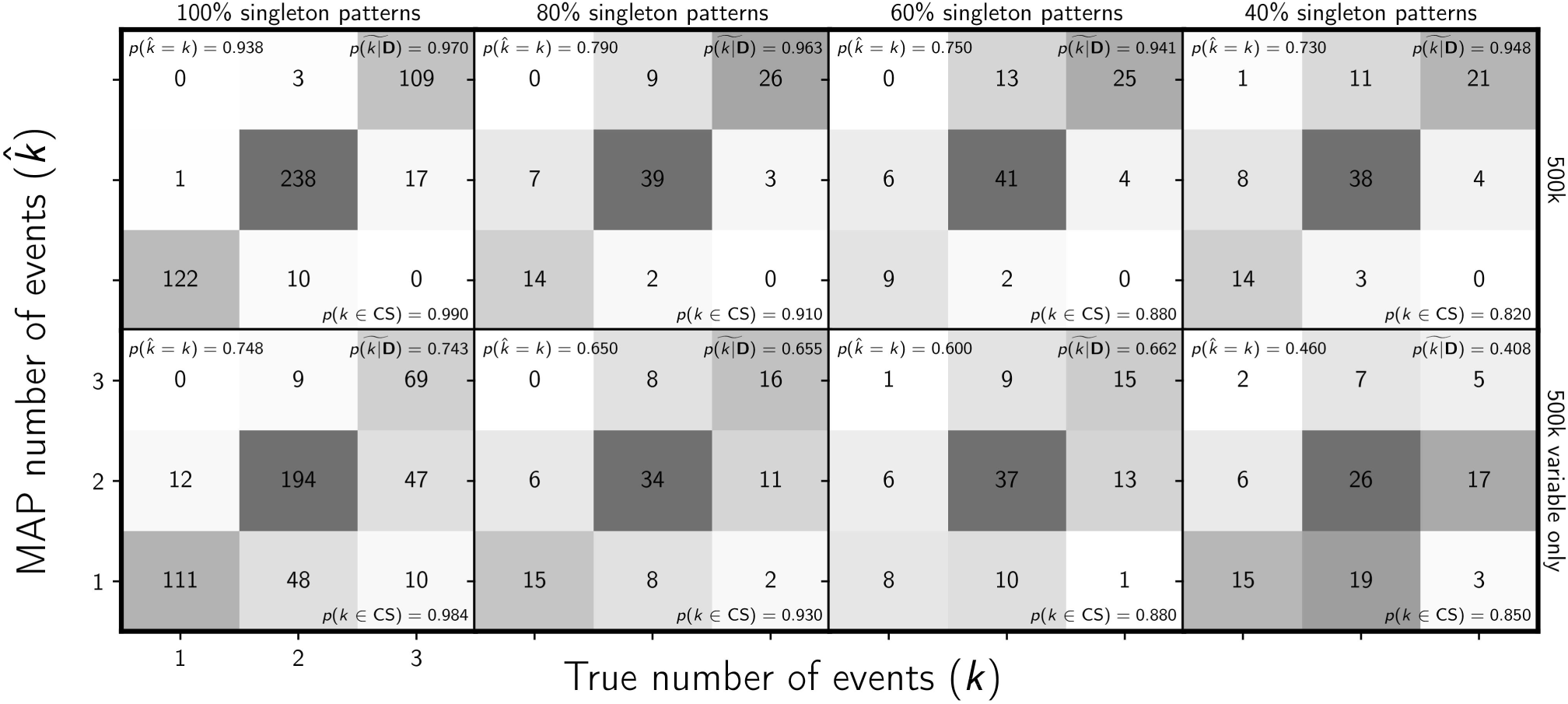
Assessing the effect of an acquisition bias against rare allele patterns on the ability of the new method to estimate the number of divergence events. The columns, from left to right, show the results when each simulated 500,000-character data set has a probability of 100%, 80%, 60%, and 40% of sampling each simulated singleton pattern. E.g., each character matrix analyzed in the far right column is missing approximately 60% of characters where all but one gene copy has the same allele. For comparison, the first column shows the results of the 500 data sets from Figure 4; the remaining columns show the results of 100 data sets. The rows show the results when (top) all sites and (bottom) only variable sites are analyzed. The number of data sets that fall within each possible cell of true versus estimated numbers of events is shown, and cells with more data sets are shaded darker. The estimates are based on the number of events with the maximum *a posteriori* (MAP) probability. For each plot, the proportion of data sets for which the number of events with the largest posterior probability matched the true number of events—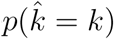—is shown in the upper left corner, the median posterior probability of the correct number of events across all data sets—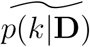—is shown in the upper right corner, and the proportion of data sets for which the true number of events was included in the 95% credible set—*p*(*k∈*CS)—is shown in the lower right. All simulated data sets had three pairs of populations. We generated the plot using matplotlib Version 2.0.0 (Hunter, 2007).

#### 3.1.5 Comparison to ABC methods

The new full-likelihood method, `ecoevolity`, does a much better job of estimating divergence times (Fig. 12) and effective population sizes (Fig. S28 & S29), than the approximate-likelihood Bayesian method, `dpp-msbayes`. This is despite the simulated data sets being “tailored” for the ABC method (i.e., loci of 200 linked base pairs). Notably, the new method does not underestimate the older divergence times like the ABC method, which suffers from saturated population-genetic summary statistics that assume an infinite-sites model of mutation (Fig. 12). Also, the ABC approach gleans no information from the data about effective population sizes; the posterior distribution nearly matches the prior for all analyses (Fig. S28 & S29). This is despite the fact that three of the four statistics used for the ABC approach summarize information about population sizes.

**Figure 12.**
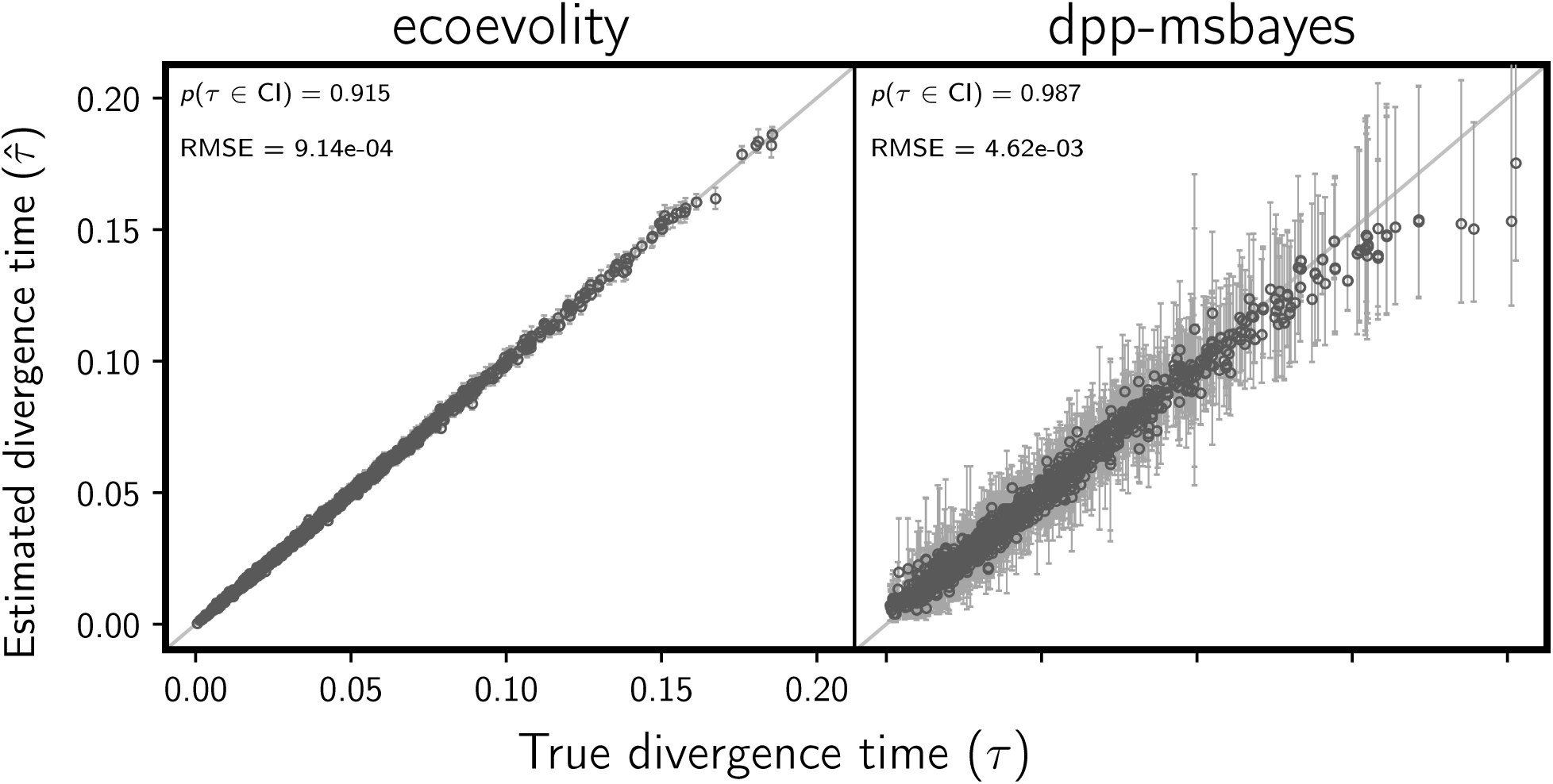
Comparing the accuracy and precision of divergence-time estimates between (left) the new full-likelihood Bayesian method, `ecoevolity`, and (right) the approximate-likelihood Bayesian method, `dpp-msbayes`. Each plotted circle and associated error bars represent the posterior mean and 95% credible interval for the time that a pair of populations diverged. Each plot consists of 1500 estimates—500 simulated data sets, each with three pairs of populations. The simulated character matrix for each population pair consisted of 200 loci, each with 200 linked sites (40,000 characters total). For each plot, the root-mean-square error (RMSE) and the proportion of estimates for which the 95% credible interval contained the true value—*p*(*t∈*CI)— is given. We generated the plot using matplotlib Version 2.0.0 (Hunter, 2007).

The new method also does a better job of estimating the number of divergence events (Fig. 13), with a median posterior probability of the correct number of events of 0.942, compared to 0.7 for the ABC method. Similarly, the new method is better at estimating the divergence model (Fig. S30), with a median posterior probability of the correct model of 0.942, compared to 0.685 for the ABC method. Importantly, the new method underestimates the number of events much less frequently (Fig. 13 & S30), which should lead to fewer erroneous interpretations of shared processes of divergence.

**Figure 13.**
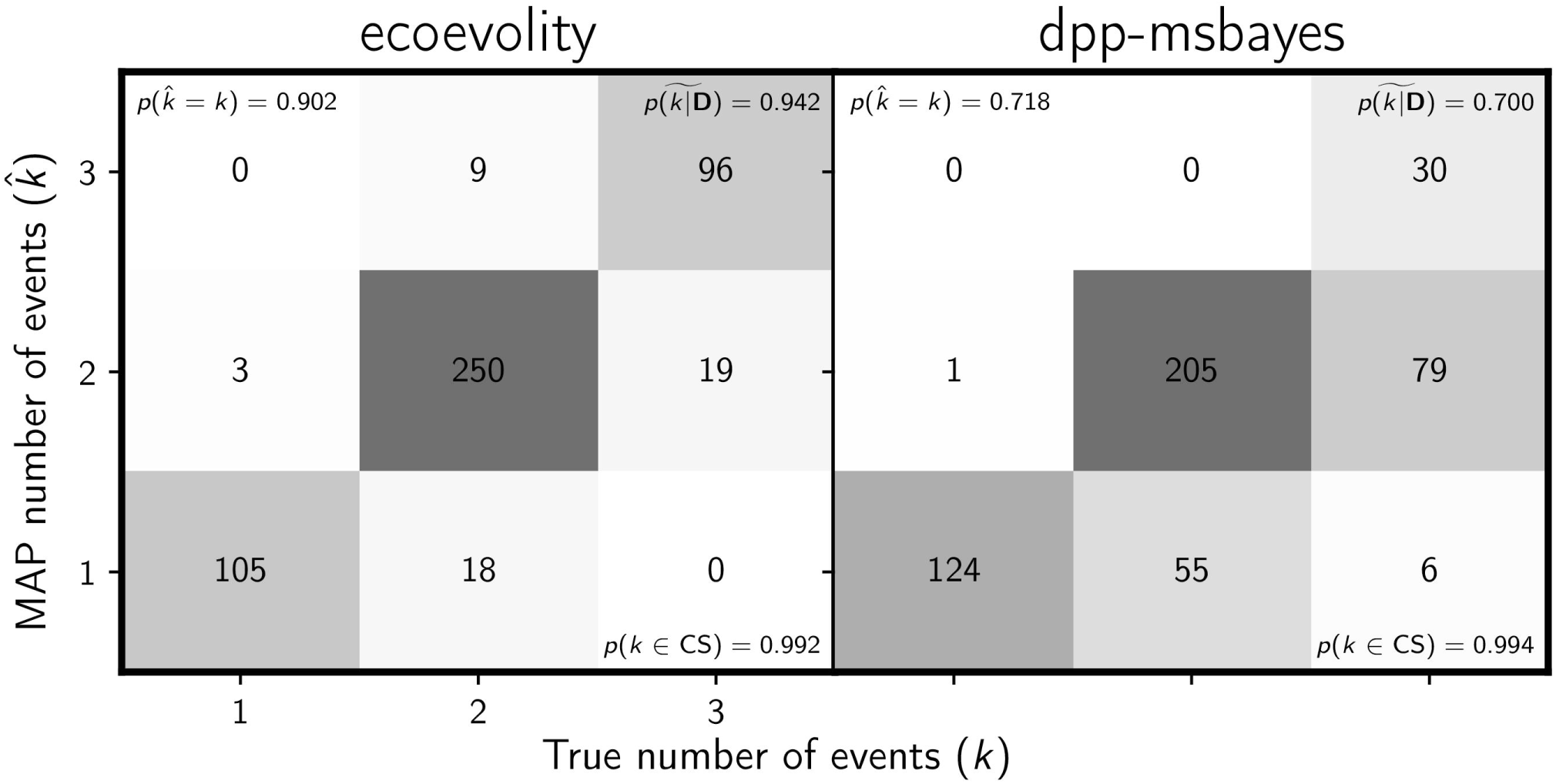
Comparing the ability to estimate the number of divergence events between (left) the new full-likelihood Bayesian method, `ecoevolity`, and (right) the approximate-likelihood Bayesian method, `dpp-msbayes`. Each plot shows the results of the analyses of 500 simulated data sets; the number of data sets that fall within each possible cell of true versus estimated numbers of events is shown, and cells with more data sets are shaded darker. Each simulated data set contained three pairs of populations, and the simulated character matrix for each pair consisted of 200 loci, each with 200 linked sites (40,000 characters total). The estimates are based on the number of events with the maximum *a posteriori* (MAP) probability. For each plot, the proportion of data sets for which the number of events with the largest posterior probability matched the true number of events—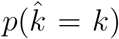—is shown in the upper left corner, the median posterior probability of the correct number of events across all data sets—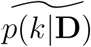—is shown in the upper right corner, and the proportion of data sets for which the true number of events was included in the 95% credible set—*p*(*k∈*CS)—is shown in the lower right. We generated the plot using matplotlib Version 2.0.0 (Hunter, 2007).

It is difficult to compare the computational effort between the two approaches, given that `ecoevolity` is collecting autocorrelated samples from the full posterior, whereas `dpp-msbayes` is collecting independent samples from a different distribution we hope is similar to the posterior. Nonetheless, the comparison is aided by the fact that the heavy computation of both methods is coded in `C`/`C++`. To compare the overall amount of computation required by the two approaches we look at the average time it takes to analyze a simulated data set on a single processor (2.6GHz Intel Xeon CPU E5-2660 v3). This was 38.8 days for `dpp-msbayes` (3,350,465 seconds) and only 33.4 minutes (2004.5 seconds) for `ecoevolity`. The majority of the runtime for the ABC method is spent simulating samples from the prior distribution. While this step can be parallelized, the likelihood computations of `ecoevolity` can also be multi-threaded. Regardless of the difficulties associated with comparing the approaches, the 1,671-fold difference in computation time clearly demonstrates the full-likelihood method is much more efficient than ABC.

### 3.2 Empirical application

When the new method is applied to all of the RADseq sites from the four pairs of *Gekko* populations, the results strongly support that all of the pairs diverged independently (Fig. 14). The results are very robust to the priors on divergence times (*τ*) and the concentration parameter (*α*) of the Dirichlet process, and to whether the polyallelic SNPs are recoded as binary (Fig. 14) or removed (Fig. S31). Likewise, the estimates of divergence times and effective population sizes are nearly identical regardless of the prior on *τ* or *α*, or whether polyallelic SNPs are recoded or removed (Fig. 15, S32, S33, & S34).

**Figure 14.**
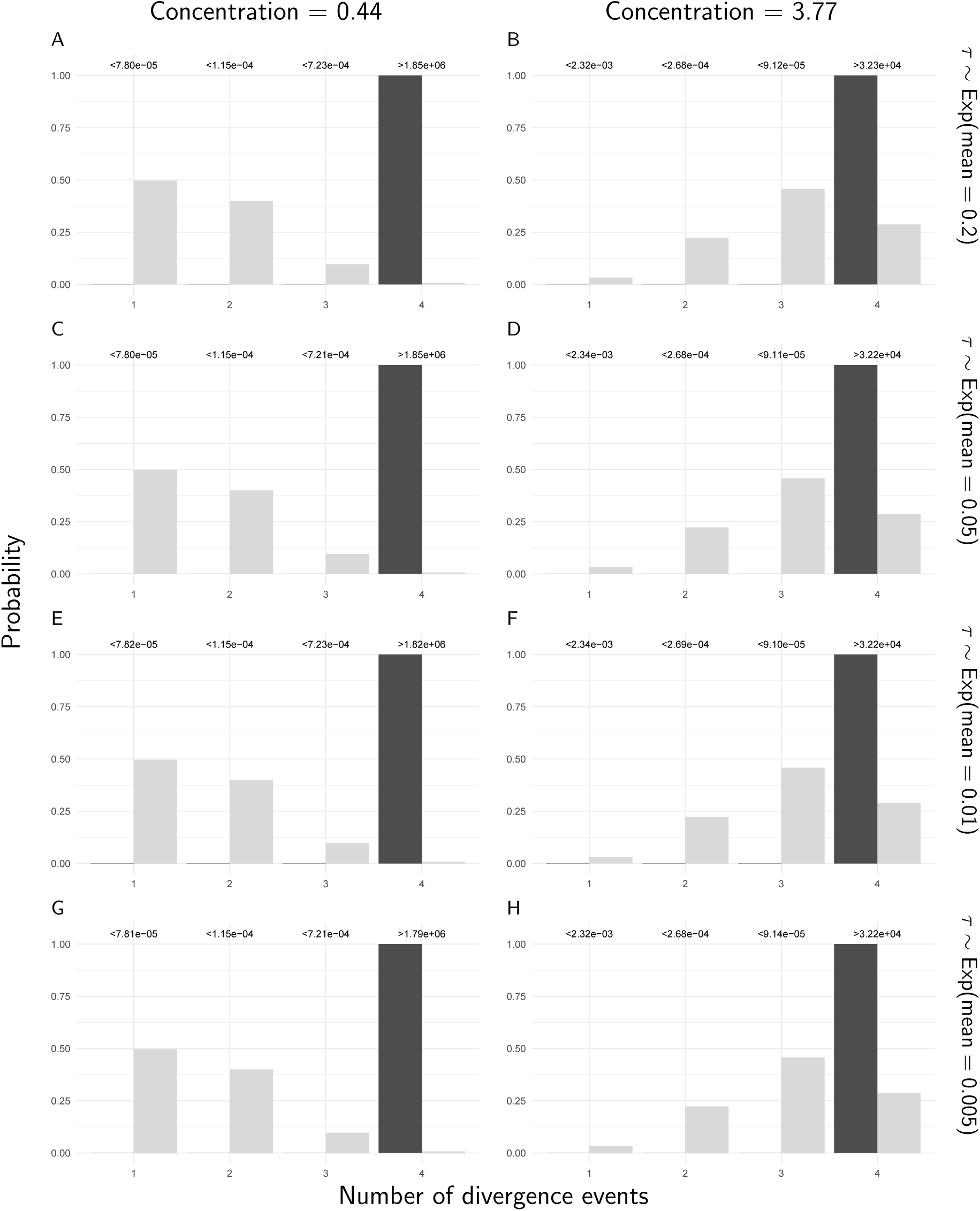
The prior (light bars) and approximated posterior (dark bars) probabilities of the number of divergence events across *Gekko* pairs of populations, under eight different combinations of prior on the divergence times (rows) and the concentration parameter of the Dirichlet process (columns). For these analyses, constant characters were included, and all characters with more than two alleles were recoded as biallelic. The Bayes factor for each number of divergence times is given above the corresponding bars. Each Bayes factor compares the corresponding number of events to all other possible numbers of divergence events. We generated the plots with ggplot2 Version 2.2.1 (Wickham, 2009).

**Figure 15.**
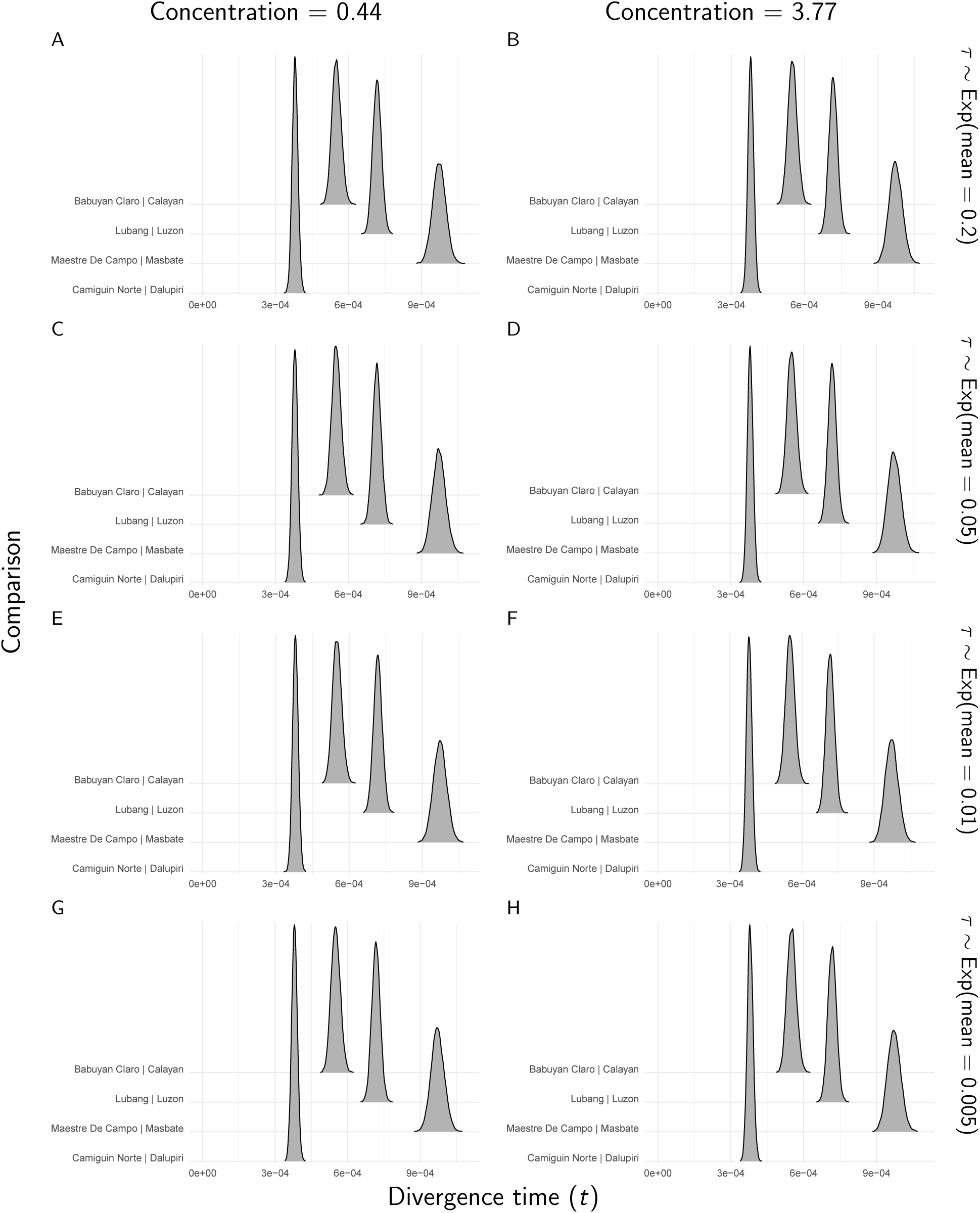
The approximate marginal posterior densities of divergence times for each *Gekko* pair of populations, under eight different combinations of prior on the divergence times (rows) and the concentration parameter of the Dirichlet process (columns). For these analyses, constant characters were included, and all characters with more than two alleles were recoded as biallelic. We generated the plots with ggridges Version 0.4.1 (Wilke, 2018) and ggplot2 Version 2.2.1 (Wickham, 2009).

However, when only variable SNPs are analyzed, the behavior is much different. First, the estimated divergence times and population sizes are clearly far too large and more sensitive to the priors on the divergence times and the concentration parameter (Fig. S35 & S36). While the true values of these parameters are obviously unknown, given the variability of these data (Table 2), and other data from these species (Siler et al., 2012, 2014), these values are clearly nonsensical. The posterior probabilities of the number of divergences are also much more sensitive to the *τ* and *α* priors, with some combinations yielding results for which three divergence events are preferred, although Bayes factors always preferred four divergences (Fig. S37). These findings are similar when the polyallelic SNPs are removed (Fig. S38–40).

The large overestimation of divergence times and population sizes is consistent with our findings from the data sets simulated with an acquisition bias against rare allele patterns (Fig. 10, S25 & S26). In these simulation-based analyses, we also saw dramatic overestimation of these parameters when constant characters were excluded. It appears that some variable character patterns are being lost during the acquisition and assembly of the RADseq data, and the model is sensitive to these missing variable sites, especially when only variable characters are analyzed.

#### 3.2.1 Empirical comparison to ABC

Figure 16 shows the dramatic difference between the results of the new full-likelihood method, `ecoevolity`, and the ABC method, `dpp-msbayes`, when analyzing a random subset of the *Gekko* data. For `dpp-msbayes`, there is strong support for a single, shared divergence (Fig. 16c), and the marginal posterior distributions of divergence times are almost completely overlapping among the three pairs of populations (Fig. 16d). In contrast, `ecoevolity` strongly supports three independent divergences (Fig. 16a), and the marginal posterior divergence-time distributions are almost completely non-overlapping (Fig. 16b). Furthermore, the computing time for `ecoevolity` and `dpp-msbayes` was 7.6 minutes and 49.3 days, respectively.

**Figure 16.**
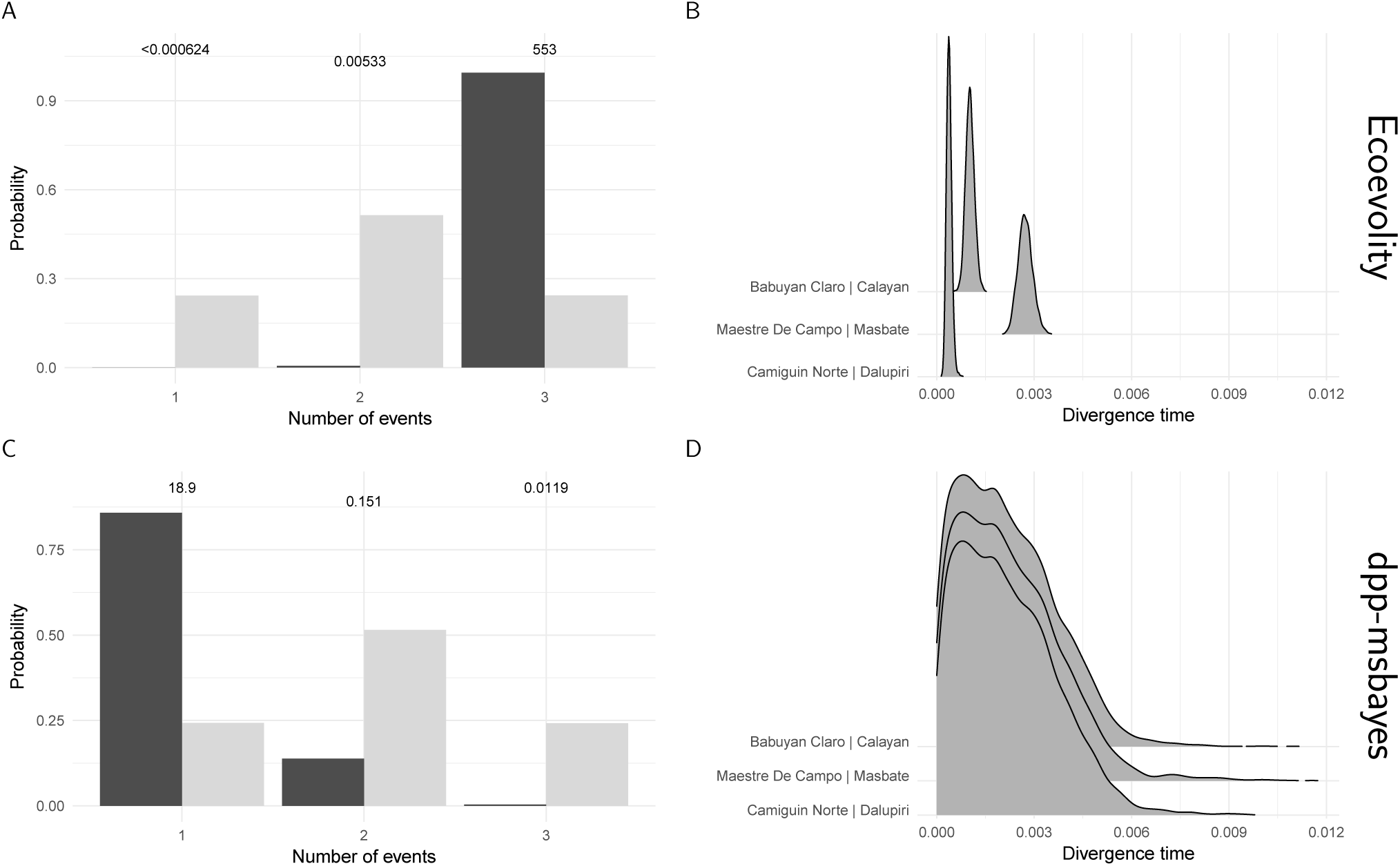
The results of the (A & B) new method full-likelihood Bayesian method, `ecoevolity`, and (C & D) approximate-likelihood Bayesian method `dpp-msbayes`, when applied to 200 RAD-seq loci randomly sampled from three of the pairs of *Gekko* populations. Plots B & D show the estimated marginal posterior densities of divergence times for each *Gekko* pair of populations. Plots A & C show the approximated prior (light bars) and posterior (dark bars) probabilities of the number of divergence events across the pairs of *Gekko* populations. The Bayes factor for each number of divergence times is given above the corresponding bars. Each Bayes factor compares the corresponding number of events to all other possible numbers of divergence events. We generated the plots with ggridges Version 0.4.1 (Wilke, 2018) and ggplot2 Version 2.2.1 (Wickham, 2009).

## 4 Discussion

Previous approaches to estimating shared divergence times based on approximate-likelihood Bayesian computation (ABC) are very sensitive to prior assumptions about divergence times and often over-cluster divergences with strong support (Oaks et al., 2013; Hickerson et al., 2014; Oaks et al., 2014; Oaks, 2014). Here, we introduced a new approach that increases the power and robustness of these inferences by leveraging all of the information in genomic data within a full-likelihood, Bayesian framework. The full-likelihood approach is much better at estimating divergence times (Fig. 12) and nuisance parameters (Fig. S28 & S29) than ABC. It is also better able to estimate the correct number of divergence events with more confidence, and is much less biased toward underestimating the number of divergence events (Fig. 13). This is especially important, because most biogeographers that use these methods are interested in testing for shared events. The increased power of the method to detect variation in divergence times and avoid spurious estimates of shared divergences will lead to fewer erroneous interpretations of shared processes of divergence.

The efficiency associated with using all of the information in the data makes the method very promising for empirical applications. For example, increasing the number of characters from 100,000 to 500,000 resulted in only modest improvements in precision (Fig. 2, S2, & S3). This suggests that the benefit of collecting more characters begins to plateau when data sets are small relative to the number of characters commonly collected via modern high-throughput sequencing technologies (e.g., the simulated 100k data sets had only 5,500 SNPs on average). Even with only 200 short (200 bp) loci, the median posterior probability of the correct number of divergence events was 0.94 (Fig. 13). Also, directly calculating the likelihood of the population history from genomic data avoids the computation necessary for approximating the likelihood via simulations. As a result, the new method, `ecoevolity`, provides better approximations of the posterior over 1000 times faster than the ABC method, `dpp-msbayes`.

### 4.1 To exclude linked characters, or not?

The increased precision and robustness associated with retaining constant characters creates an interesting question when analyzing DNA sequence data from reduced-representation genomic libraries: Is it better to analyze all the data and violate the assumption that the characters are unlinked, or suffer a large loss of data to avoid violating that assumption? Several of our results suggest retaining all the data is preferable. First of all, the method is much better at estimating the divergence times, effective population sizes, and the correct number of events with high posterior probability when analyzing linked sequences of characters compared to when only one variable character per locus is analyzed (Fig. 5 and 6, and Fig. S12–16 and 18). Second, retaining all the data makes the method more robust to dataacquisition biases (Fig. 10 and 11 and Fig. S25 and 26), which are common in alignments from reduced-representation genomic libraries (Harvey et al., 2015; Linck and Battey, 2017). Third, the results from the *Gekko* RADseq data are reasonable and robust to prior assumptions when all data are analyzed, but nonsensical and sensitive to prior assumptions when only variable characters are analyzed. Our simulations suggest this is due to the filtering of the character patterns that occurred when assembling these data.

Perhaps most striking is how much better the method estimates the number of divergence events when all the data are used. For example, across the 500,000-character data sets, the median posterior probability of the correct number of divergence events is over 0.94 regardless of the linked characters or pattern-acquisition biases we simulated (Fig. 4, 6, & 11). For comparison, these values are as low as 0.41 when constant characters are removed (Fig. 11).

We caution against generalizing our findings of favorable performance with linked loci to other methods that assume unlinked characters. However, Chifman and Kubatko (2014) found quartet inference of splits in multi-species coalescent trees from SNP data was also robust to the violation of unlinked characters. Our results show the amount of data that is discarded to avoid linked characters can far outweigh the effects of violating the assumption of unlinked characters. When analyzing linked loci with a method that assumes unlinked characters, using simulations to assess the effect of linkage on the method may be worth the effort in order to bring more data to bear.

### 4.2 Philippine *Gekko*

We purposefully selected a challenging empirical test case for the new method. Each of the four pairs of populations of *Gekko* occur on two different oceanic islands that were never connected. Thus, we do not expect shared divergence times across the pairs. However, based on previous findings (Siler et al., 2012, 2014), all of these pairs likely diverged very recently. This is a challenging region of parameter space for this type of method: Very similar and recent divergence times that are nonetheless independent. Our results strongly support independent divergences, despite all four pairs diverging very recently. (Fig. 14 & 15).

We found similar results when we analyzed a random subset of 200 loci from three of the pairs of *Gekko* populations (Fig. 16a&b). In contrast, when we analyzed these 200 loci with the ABC method, `dpp-msbayes`, we found strong support for the opposite conclusion that all three pairs diverged at the same time (Fig. 16c&d). In addition, `ecoevolity` took approximately 9,300-fold less computing time than `dpp-msbayes`. These results demonstrate that using the likelihood from genomic data provides enough information to efficiently and unambiguously separate divergences across very narrow timescales, and avoids erroneous inferences of shared divergences.

### 4.3 Caveats

This method is subject to the caveats associated with all model-based statistical methods, however, there are two caveats that are worth emphasizing with specific reference to the types of models we explored here. First, it is important to keep in mind that when modeling the divergence of two populations, the time of the divergence and the mutation rate are inextricably linked. Thus, we cannot learn about the relative rates of mutation among pairs of populations when also trying to estimate their divergence times. Unlike previous methods (Hickerson et al., 2006; Huang et al., 2011; Oaks, 2014), we allow priors to be placed on mutation rates, to allow uncertainty to be incorporated into the model. However, the priors on the mutation rates need to be informative if one hopes to be able to estimate the divergence times.

Second, the new method does not allow migration after populations diverge. This is a weakness compared to ABC approaches to this problem (Huang et al., 2011; Oaks, 2014). However, given the biases and sensitivity to priors exhibited by the ABC methods even when migration is ignored (Oaks et al., 2013, 2014; Oaks, 2014), modeling migration with these methods is not advisable without thorough simulation-based analyses to assess their statistical behavior.

## 5 Conclusions

We introduced a new Bayesian model-choice method for estimating shared divergence times across taxa. By using the full likelihood and genome-scale data, the new method is more accurate, precise, robust, and efficient than existing methods based on approximate likelihoods. This new tool will allow biologists to leverage comparative genomic data to test hypotheses about the effects of environmental change on diversification.

## 6 Funding

This work was supported by the National Science Foundation (grant numbers DBI 1308885 and DEB 1656004 to JRO).

## 7 Acknowledgments

I thank Mark Holder and Vladimir Minin for discussions that helped preserve my sanity while working in units of time and population size under the coalescent. I also thank David Bryant and Mark Holder for helpful advice on approaching the Hastings ratios of the multivariate Metropolis-Hastings proposals. Support and ideas from Adam Leaché and his lab group greatly improved this work. Early testing of the software by Matt McElroy helped identify bugs. I also thank the members of the Phyletica Lab (the phyleticians) for constructive feedback on multiple drafts of this paper. I am grateful to David Bryant, one anonymous referee, and Associate Editor, Laura Kubatko, for constructive reviews that greatly improved this work. Most of the computational work for this project was performed on the Auburn University Hopper Cluster. This paper is contribution number 879 of the Auburn University Museum of Natural History.

## Appendix A Multivariate Metropolis-Hastings Moves

Here, we describe the multivariate Metropolis-Hastings moves that improve mixing of the MCMC chain when there are strong correlations between divergence times, effective population sizes, and mutation rates. The probability of accepting a Metropolis-Hastings proposal is determined by the product of three terms, the first two of which are the ratios of the likelihood and prior probability densities of the proposed state to the current state of the model. The third term, the Hastings ratio (HR), accounts for any difference in the probability of the proposed move versus the probability of the move that would exactly reverse the proposed state back to the current state of the model. Below, we detail the Hastings ratios for two of our multivariate moves.

### A.1 TimeRootSizeMixer proposal

One case of poor mixing can occur for pairs that diverge long enough ago such that only a single coalescence occurs within the root for most loci. In this scenario there is very little information in the character patterns about the size of the ancestral population, and so the divergence time and root population size become highly correlated (i.e., an older divergence time and smaller root size explain the data equally well as a younger divergence time and larger root size). We used expectations under the coalescent to design a proposal to better sample this correlated region of parameter space. To simplify notation, throughout this section we will use *N* in place of 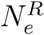 to denote the effective size of the ancestral population.
when coalescence of gene lineages is complete within a sampled pair of populations, only two lineages coalesce within the ancestral population. In this case, the expected height of the root of a gene tree is equal to *τµ* + 2*Nµ*, in units of time determined by *µ* (e.g., expected substitutions per site if *µ* = 1). The purpose of our move is to keep the expected root height of the gene trees of the proposed state equal to the current state,

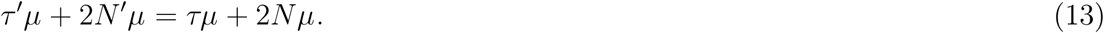

The mutation rate cancels, giving us the following relationship to uphold during our proposal,

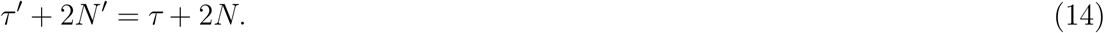

This relationship will allow us to jointly and efficiently explore the space of *τ* and *N* when there is little information in the data to tease them apart.

For population pair *i*, we first d raw a u niform r andom d eviate, *u ∼* Uniform(*-λ, λ*), where *λ* is a tuning parameter that can be adjusted to improve the acceptance rate of the proposal. Next, we propose a new value for the effective population size of the root population

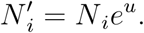

Now, we use the relationship in Equation 14 to determine the corresponding proposed value for the population divergence time,

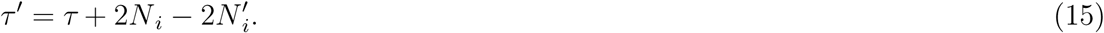

The uniform deviate to reverse this move is simply *u*^*′*^ = *-u*.

To get the Hastings ratio for this move, we use the formula of Green (1995),

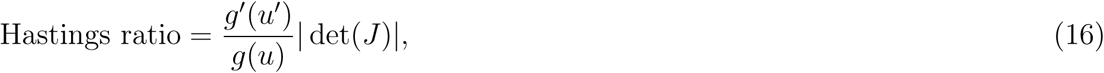

which is the ratio of the probability of drawing the random deviate that would reverse the proposed move to the probability of drawing the random deviate of the proposed move, multiplied by the absolute value of the determinant of a Jacobian matrix. Because the forward and reverse random deviates are uniform, 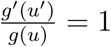, and the Hastings ratio reduces to just the Jacobian term,

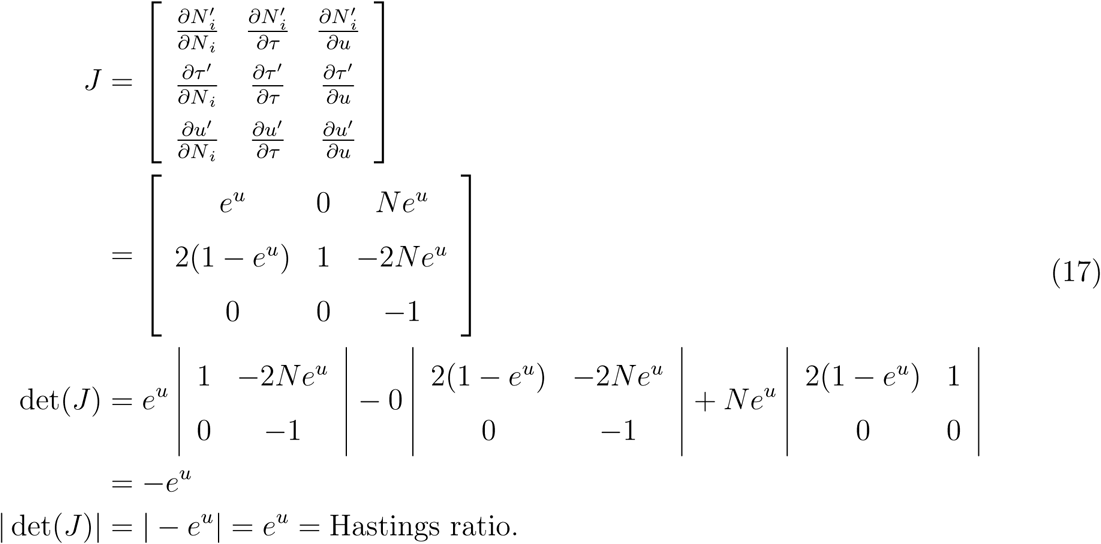

Notice that the change to *τ* also changes the divergence times of all the pairs that currently share this divergence time with pair *i*. So, the efficiency of this move can be hindered when there is a lot of sharing of divergence times. However, we can easily extend this move to change the sizes of the ancestral population of pairs *j, k, … n* that share their divergence time with pair *i*. To do this, we, again, adhere to the relationship in Equation 14:

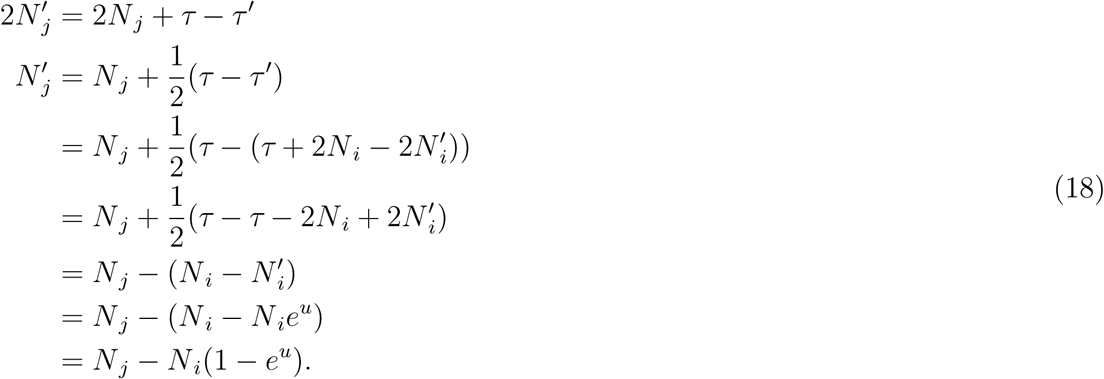

The Jacobian term then becomes

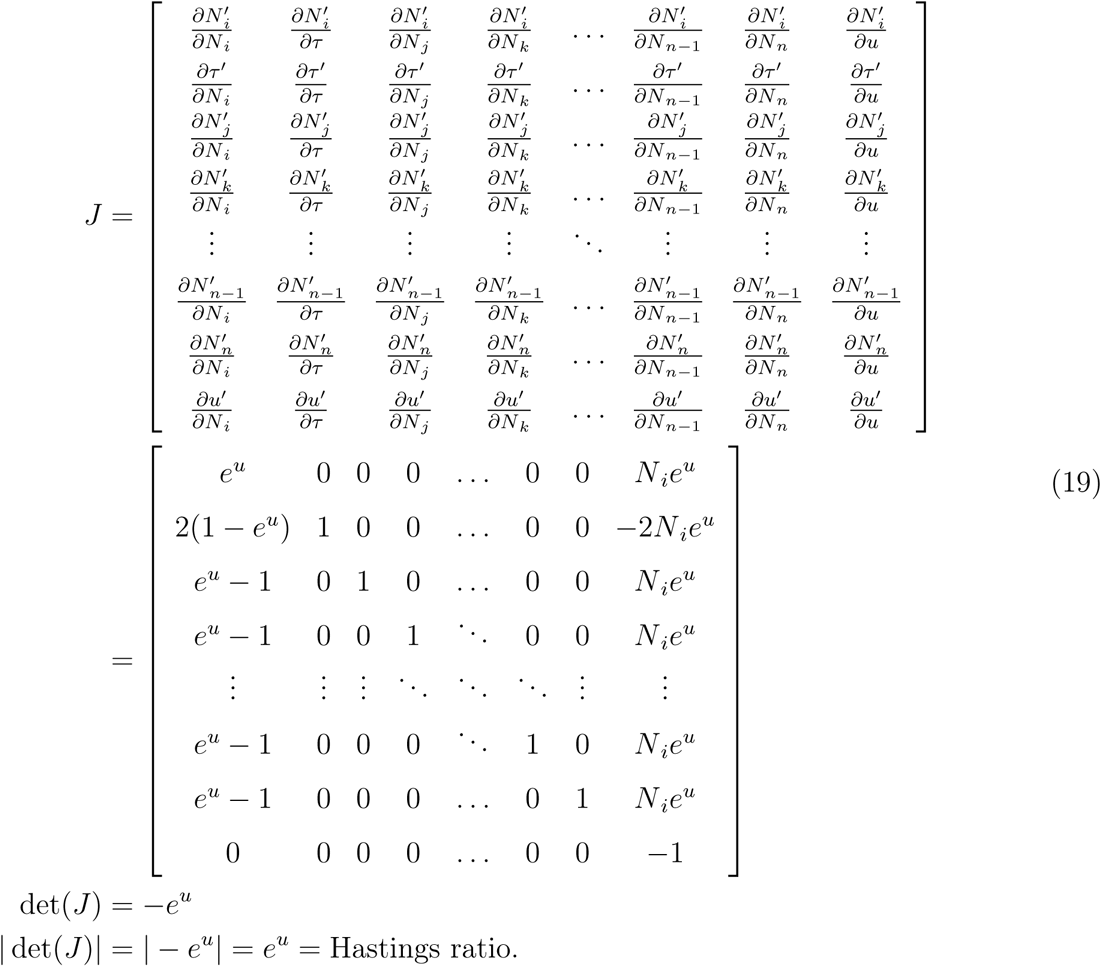

### A.2 TimeSizeRateMixer proposal

This proposal is designed to improve mixing of the MCMC chain when there are strong posterior correlations among divergence time, population size, and mutation rate parameters. It does so by jointly scaling these parameters according to the direction (positive or negative) of the posterior correlations we often observed when analyzing simulated data. The divergence time was often positively correlated with the effective sizes of the descendant populations, and negatively correlated with the mutation rate and effective population size of the ancestral population.

For a given divergence time, *τ*_*i*_, we first draw a random uniform deviate, *u ∼*Uniform(–*λ, λ*), where *λ* is, again, a tuning parameter to adjust the proposal’s acceptance rate. We use this random deviate to propose a new value for the divergence time,

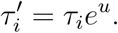

Next, we visit each population pair that is associated with this divergence time, and propose the following updates to the pair’s parameters, if they are being estimated (i.e., not fixed):

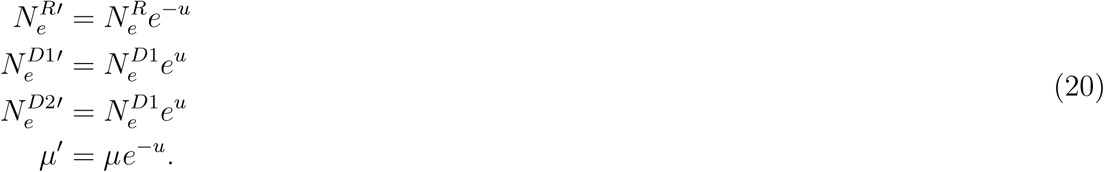

When doing so, we keep track of the total number of parameters that have been updated, denoted as *n*, and how many of these were scaled by *e*^*u*^, denoted *m*; the remaining *n–m* parameters were scaled by *e*^*-u*^.

Given *n* and *m*, we can again use Green’s 1995 formula (Equation 16 above) to determine the Hastings ratio for this proposal. Once again, 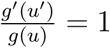, because the random deviates are uniform. Using *θ*_1_, *…, θ*_*m*_ and *θ*_*m*+1_, *…, θ*_*n*_ to denote the parameters that have been scaled by *e*^*u*^ and *e*^*-u*^, respectively, the Jacobian term is

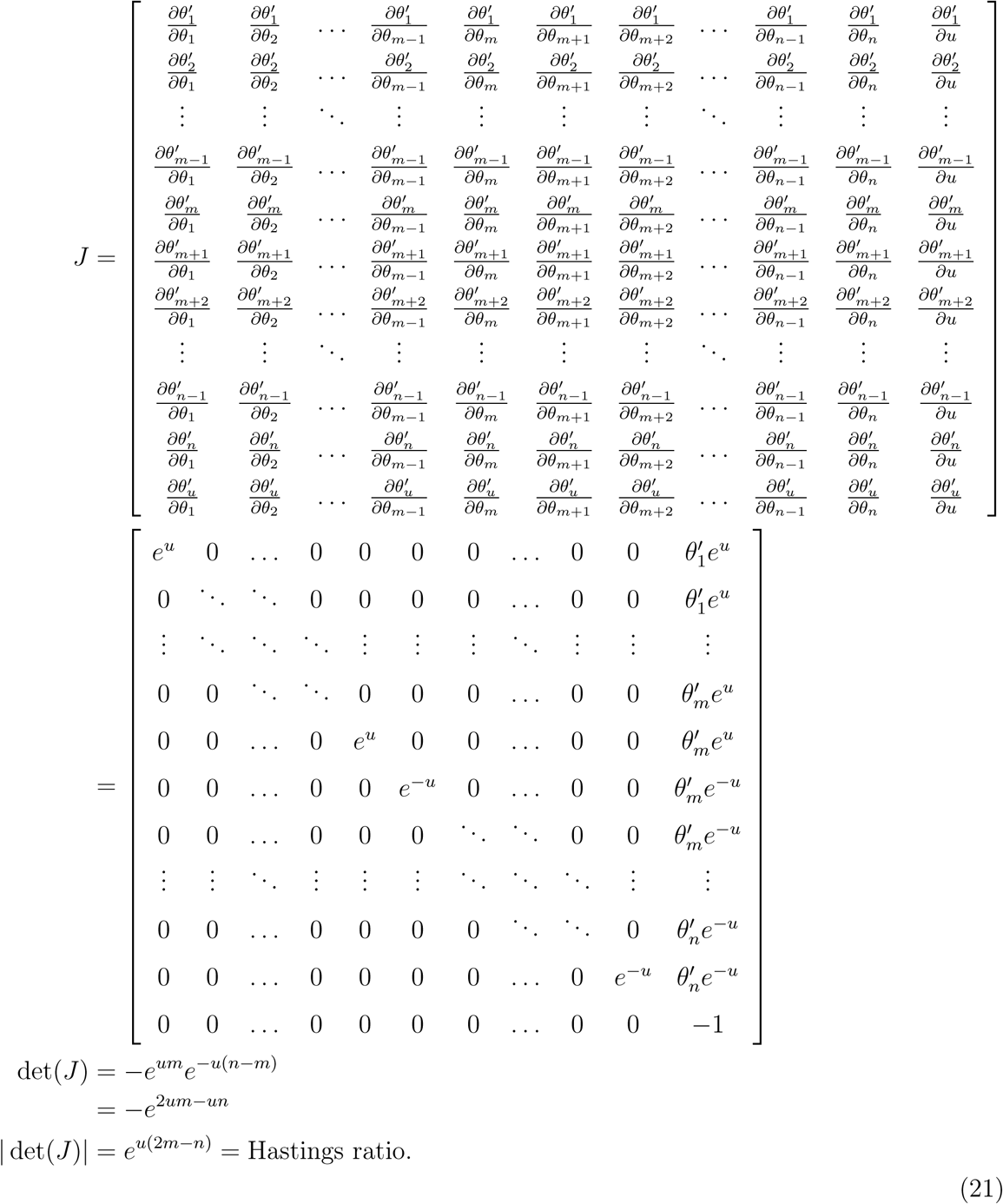

## Supporting Information

**Figure S1.**
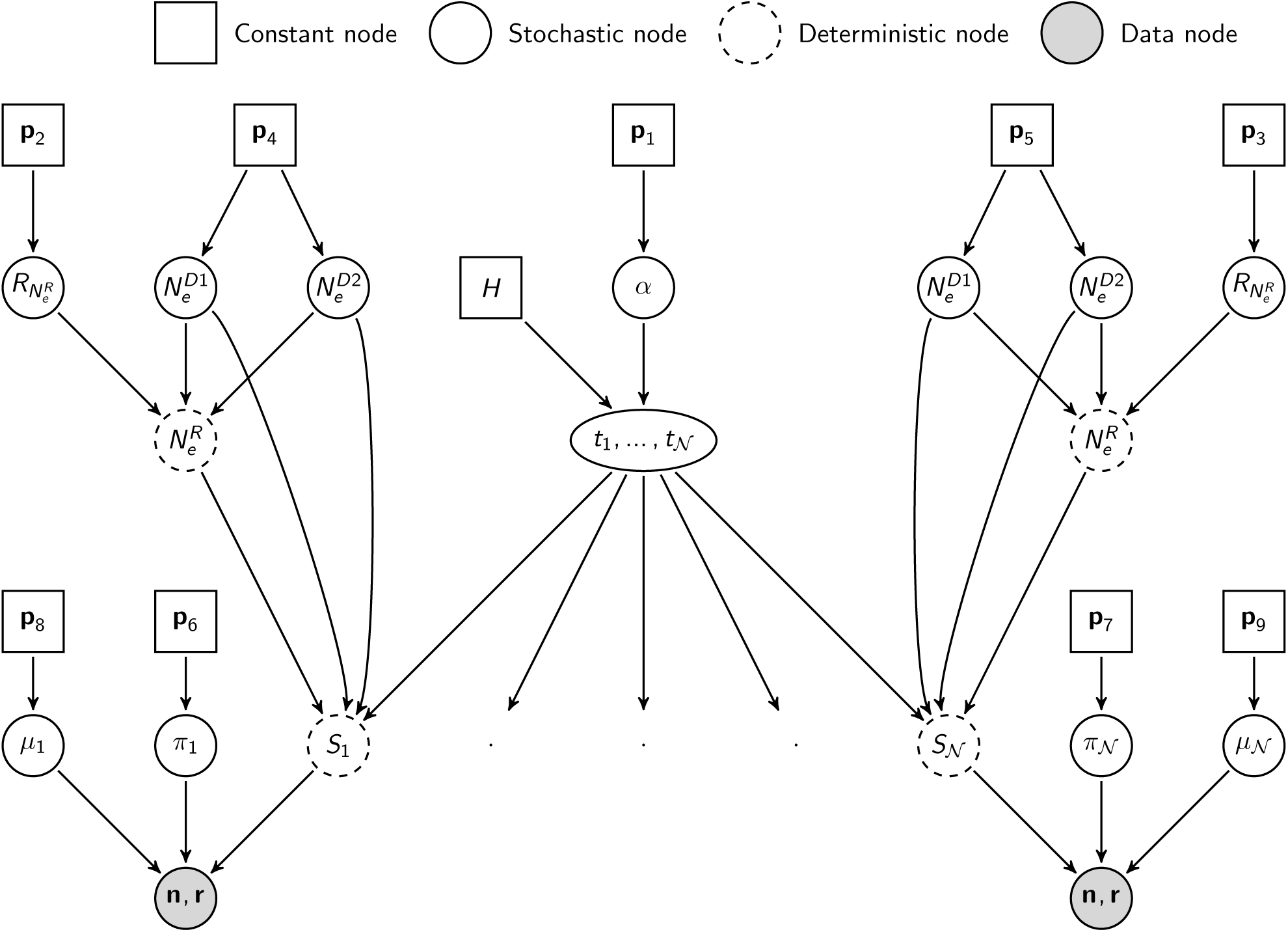
A directed graph representation of the model implemented in `ecoevolity`. Each constant node represents the parameters of a prior for which the prior distribution and values of the parameters can be choosen by the investigator.

**Figure S2.**
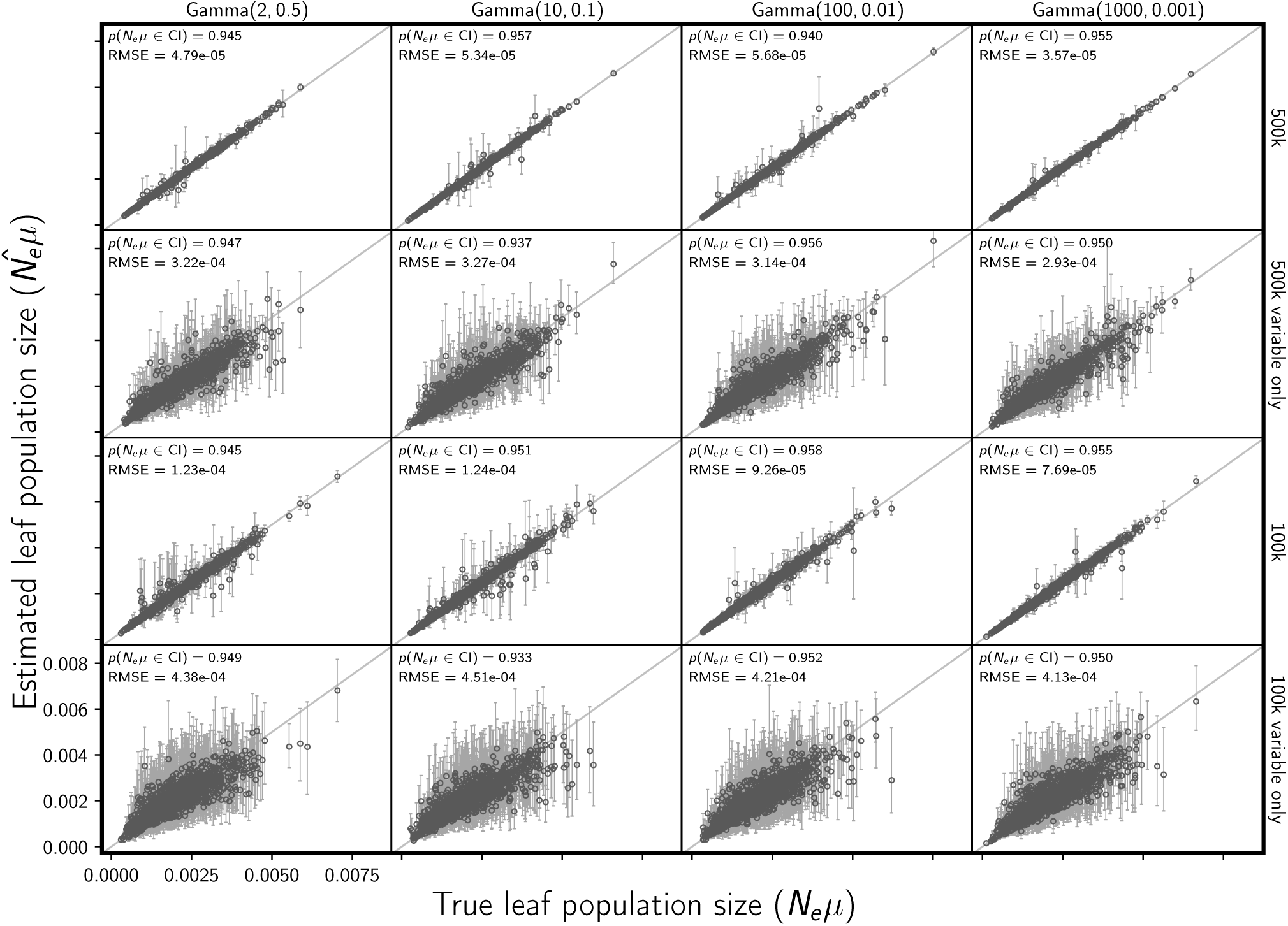
The accuracy and precision of estimates of the descendant (leaf) population sizes (scaled by the mutation rate), when data are simulated and analyzed under the same model (i.e., no model misspecification). The columns show the results from different distributions on the relative effective size of the ancestral population, decreasing in variance from left to right. For the first two and last two rows, the simulated character matrix for each population had 500,000 and 100,000 characters, respectively. The first and third rows show the results of analyses using all characters, whereas the second and fourth rows show the results when only variable characters are used. Each plotted circle and associated error bars represent the posterior mean and 95% credible interval. Each plot consists of 3000 estimates—500 simulated data sets, each with three pairs of populations. For each plot, the root-mean-square error (RMSE) and the proportion of estimates for which the 95% credible interval contained the true value—*p*(*N*_*e*_*µ∈*CI)—is given. We generated the plot using matplotlib Version 2.0.0 (Hunter, 2007).

**Figure S3.**
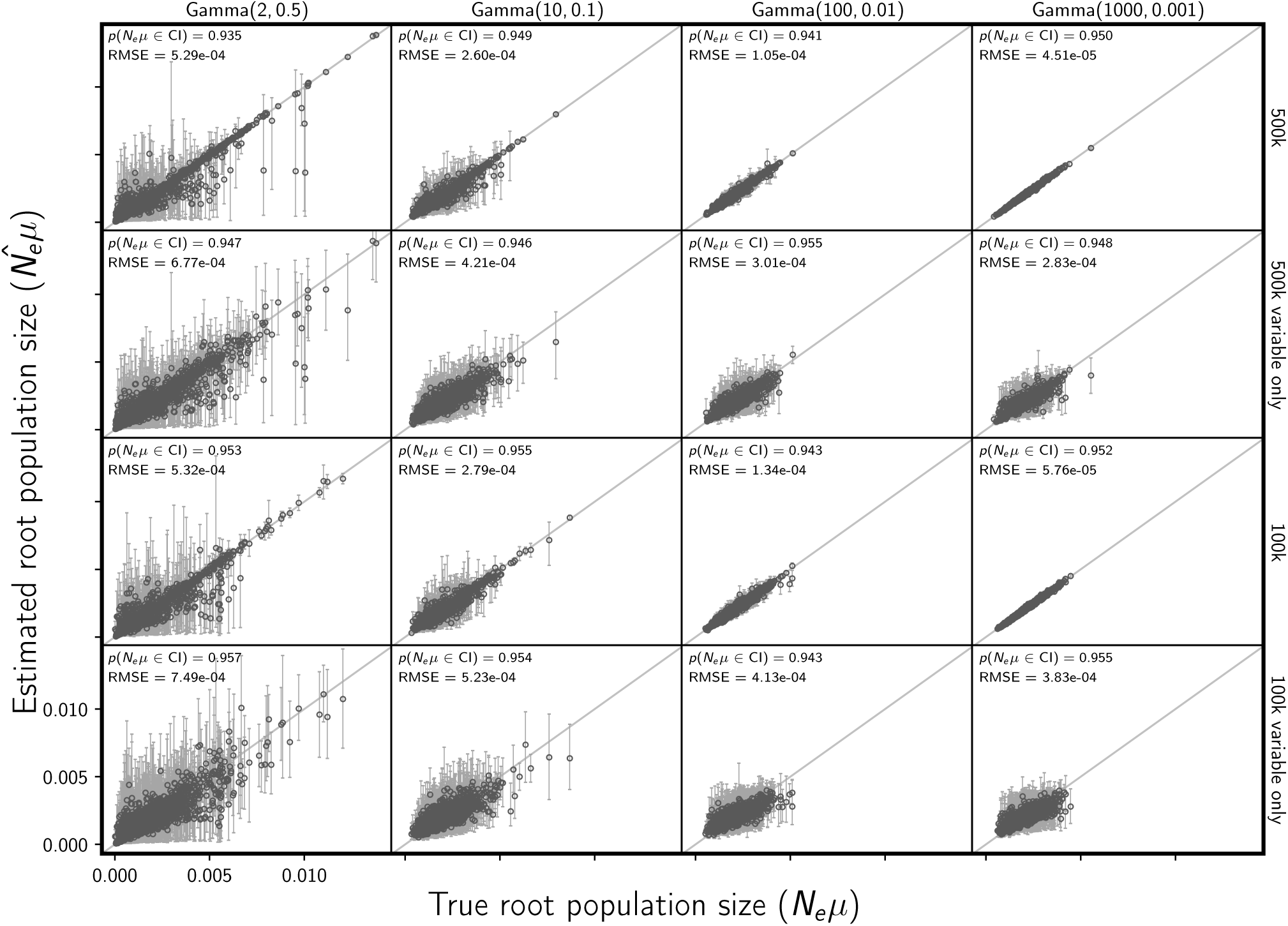
The accuracy and precision of estimates of the ancestral (root) population size (scaled by the mutation rate), when data are simulated and analyzed under the same model (i.e., no model misspecification). The columns show the results from different distributions on the relative effective size of the ancestral population, decreasing in variance from left to right. For the first two and last two rows, the simulated character matrix for each population had 500,000 and 100,000 characters, respectively. The first and third rows show the results of analyses using all characters, whereas the second and fourth rows show the results when only variable characters are used. Each plotted circle and associated error bars represent the posterior mean and 95% credible interval. Each plot consists of 1500 estimates—500 simulated data sets, each with three pairs of populations. For each plot, the root-mean-square error (RMSE) and the proportion of estimates for which the 95% credible interval contained the true value—*p*(*N*_*e*_*µ∈*CI)—is given. We generated the plot using matplotlib Version 2.0.0 (Hunter, 2007).

**Figure S4.**
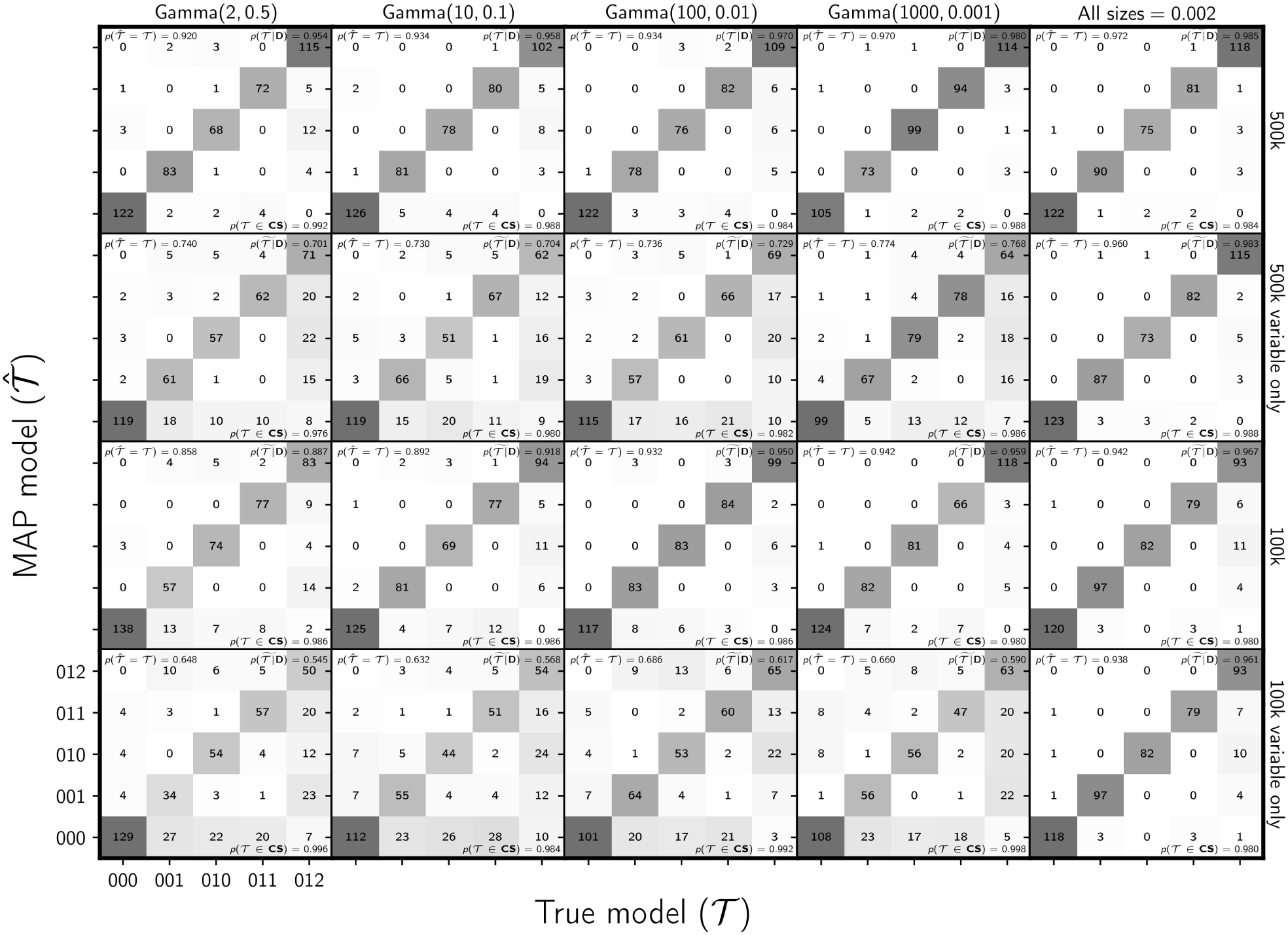
The ability of the new method to estimate the model of divergence when data are simulated and analyzed under the same model (i.e., no model misspecification). The first four columns show the results from different distributions on the relative effective size of the ancestral population, decreasing in variance from left to right. The fifth column shows results when the effective size (*N*_*e*_*µ*) of all populations is fixed to 0.002. For the first two and last two rows, the simulated character matrix for each population had 500,000 and 100,000 characters, respectively. The first and third rows show the results of analyses using all characters, whereas the second and fourth rows show the results when only variable characters are used. Each plot shows the results of the analyses of 500 simulated data sets, each with three population pairs; the number of data sets that fall within each possible cell of true versus estimated model is shown, and cells with more data sets are shaded darker. Each model is represented along the plot axes by three integers that indicate the divergence category of each pair of populations (e.g., 011 represents the model in which the second and third pair diverge at the same time, but separately from the first). The estimates are based on the model with the maximum *a posteriori* (MAP) probability. For each plot, the proportion of data sets for which the model with the largest posterior probability matched the true model—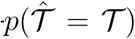—is shown in the upper left corner, the median posterior probability of the correct model across all data sets—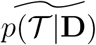—is shown in the upper right corner, and the proportion of data sets for which the true model was included in the 95% credible set— *p*(𝒯*∈*CS)—is shown in the lower right. We generated the plot using matplotlib Version 2.0.0 (Hunter, 2007).

**Figure S5.**
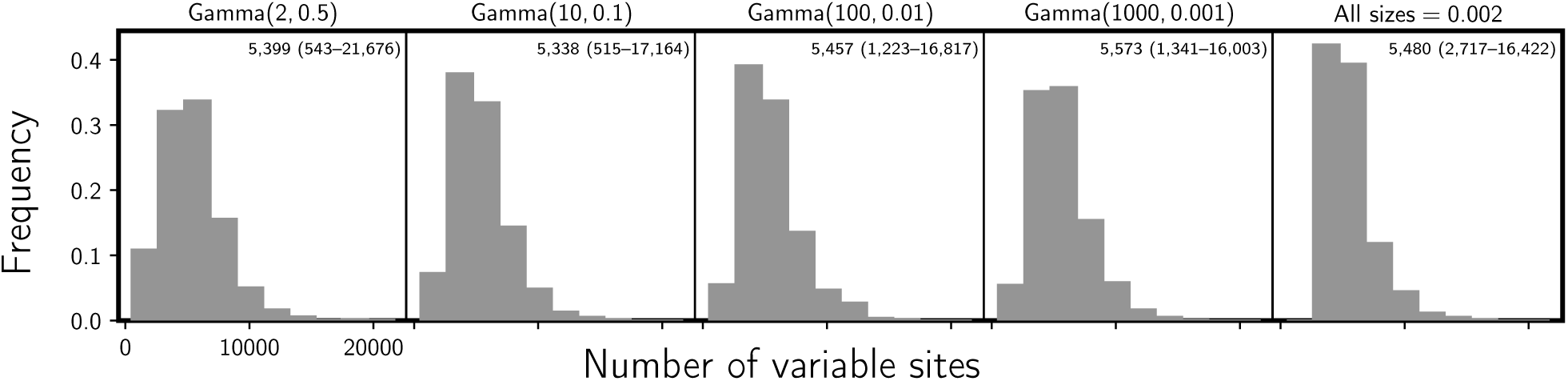
The number of variable characters in the simulated data sets with 100,000 unlinked and unfiltered characters per pair of populations. 500 data sets were simulated for each setting on the relative size of the ancestral population (indicated above each plot). The mean and range across the 500 data sets is indicated in the upper right corner of each plot. We generated the plot using matplotlib Version 2.0.0 (Hunter, 2007).

**Figure S6.**
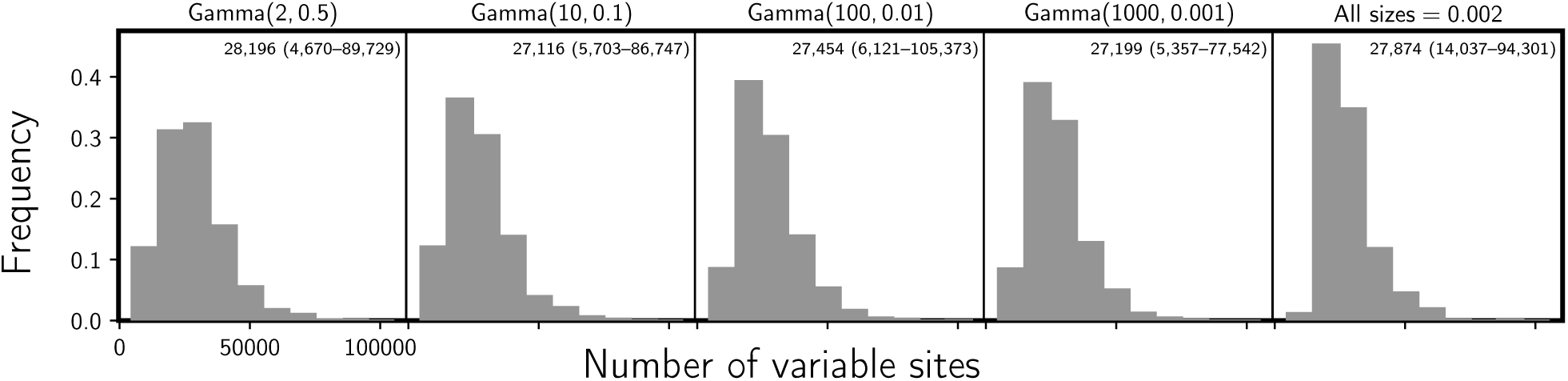
The number of variable characters in the simulated data sets with 500,000 unlinked and unfiltered characters per pair of populations. 500 data sets were simulated for each setting on the relative size of the ancestral population (indicated above each plot). The mean and range across the 500 data sets is indicated in the upper right corner of each plot. We generated the plot using matplotlib Version 2.0.0 (Hunter, 2007).

**Figure S7.**
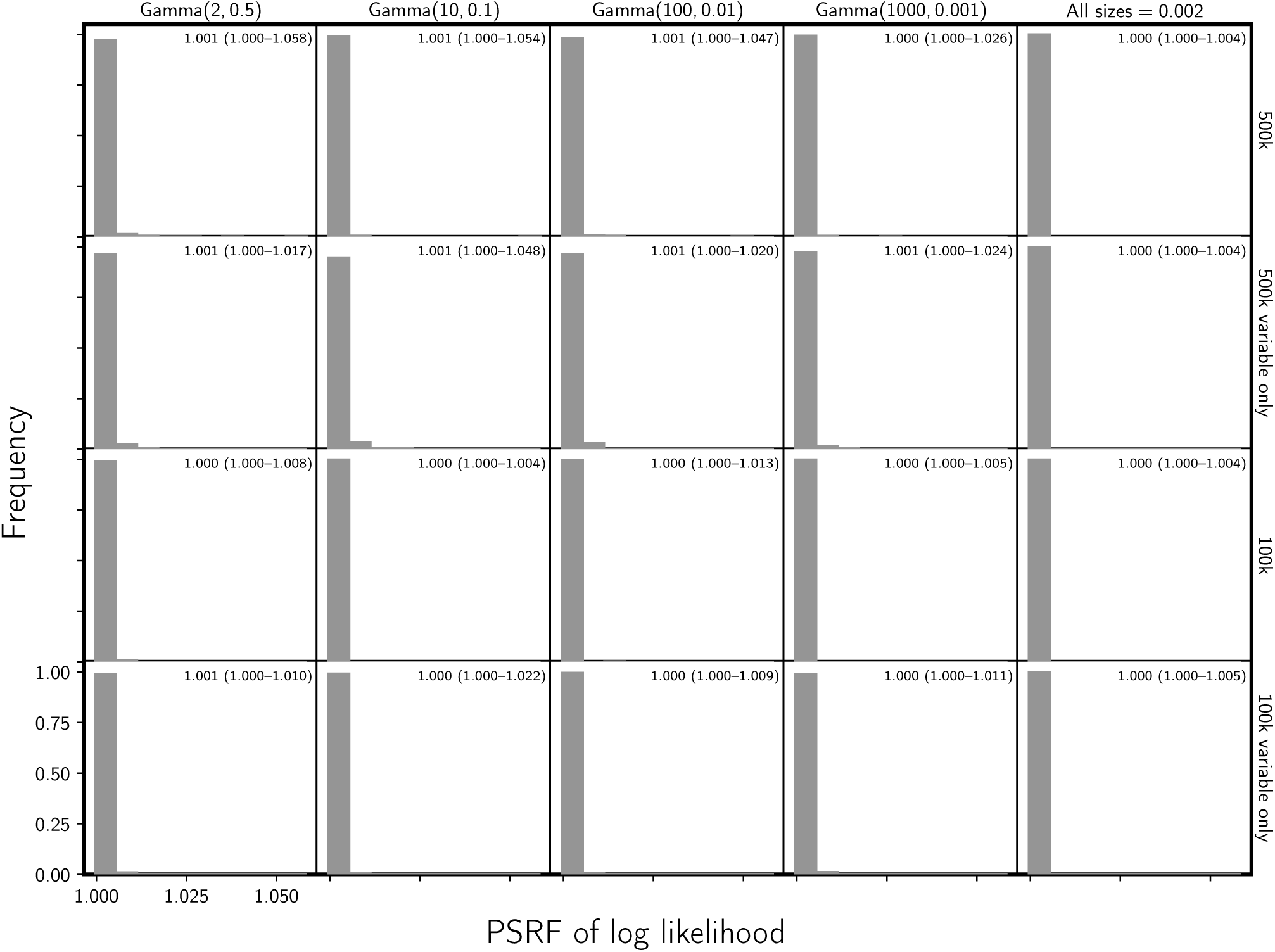
Histograms of the potential scale reduction factor (the square root of Equation 1.1 in Brooks and Gelman 1998) for the log likelihood across the three MCMC chains run for each simulated data set. The first four columns show the results from different distributions on the relative effective size of the ancestral population, decreasing in variance from left to right. The fifth column shows results when the effective size (*N*_*e*_*µ*) of all populations is fixed to 0.002. For the first two and last two rows, the simulated character matrix for each population had 500,000 and 100,000 characters, respectively. The first and third rows show the results of analyses using all characters, whereas the second and fourth rows show the results when only variable characters are used. The mean and range across the 500 data sets is indicated in the upper right corner of each plot. We generated the plot using matplotlib Version 2.0.0 (Hunter, 2007).

**Figure S8.**
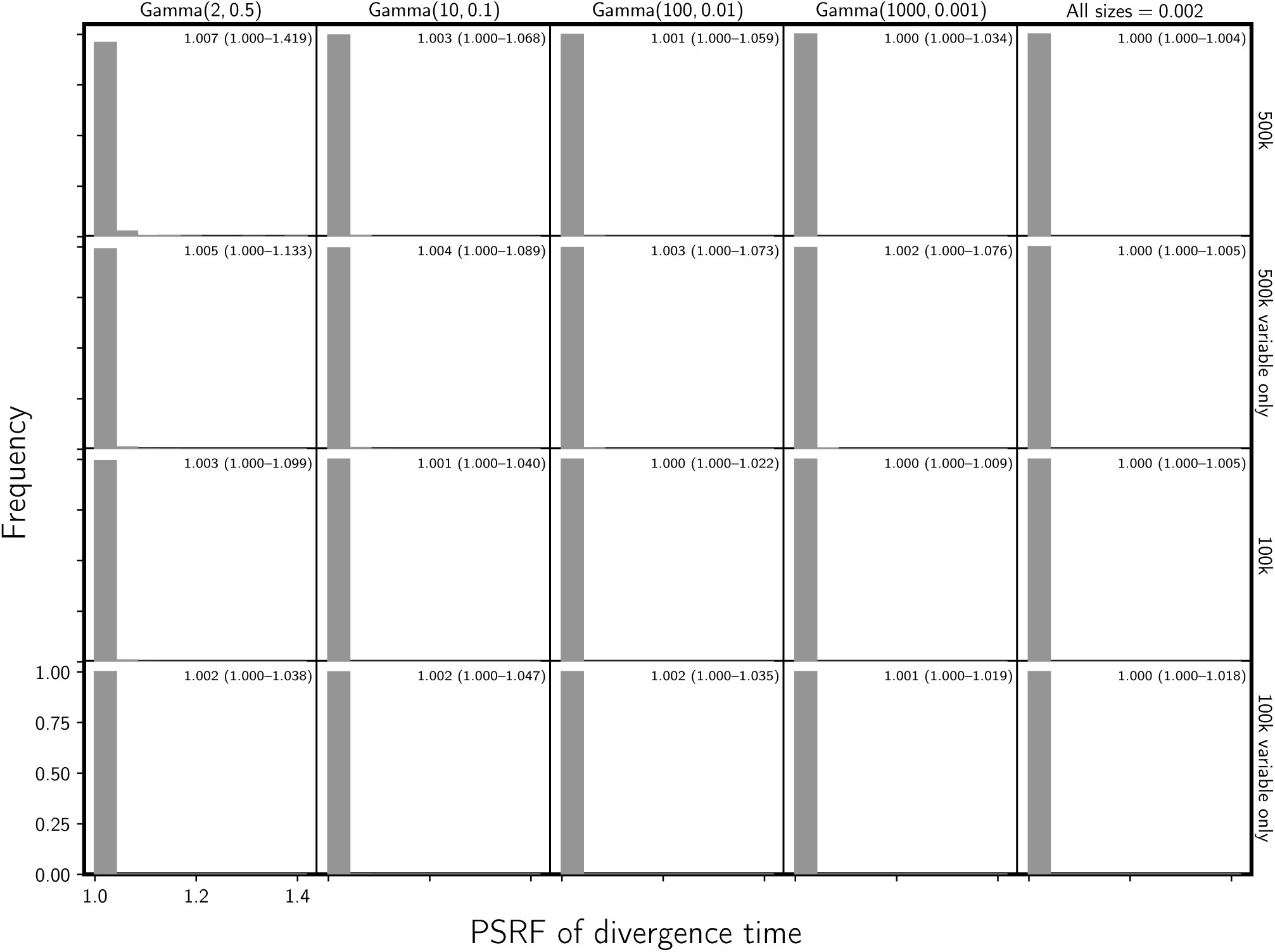
Histograms of the potential scale reduction factor (the square root of Equation 1.1 in Brooks and Gelman 1998) for the divergence times across the three MCMC chains run for each simulated data set. The first four columns show the results from different distributions on the relative effective size of the ancestral population, decreasing in variance from left to right. The fifth column shows results when the effective size (*N*_*e*_*µ*) of all populations is fixed to 0.002. For the first two and last two rows, the simulated character matrix for each population had 500,000 and 100,000 characters, respectively. The first and third rows show the results of analyses using all characters, whereas the second and fourth rows show the results when only variable characters are used. The mean and range across the 500 data sets is indicated in the upper right corner of each plot. We generated the plot using matplotlib Version 2.0.0 (Hunter, 2007).

**Figure S9.**
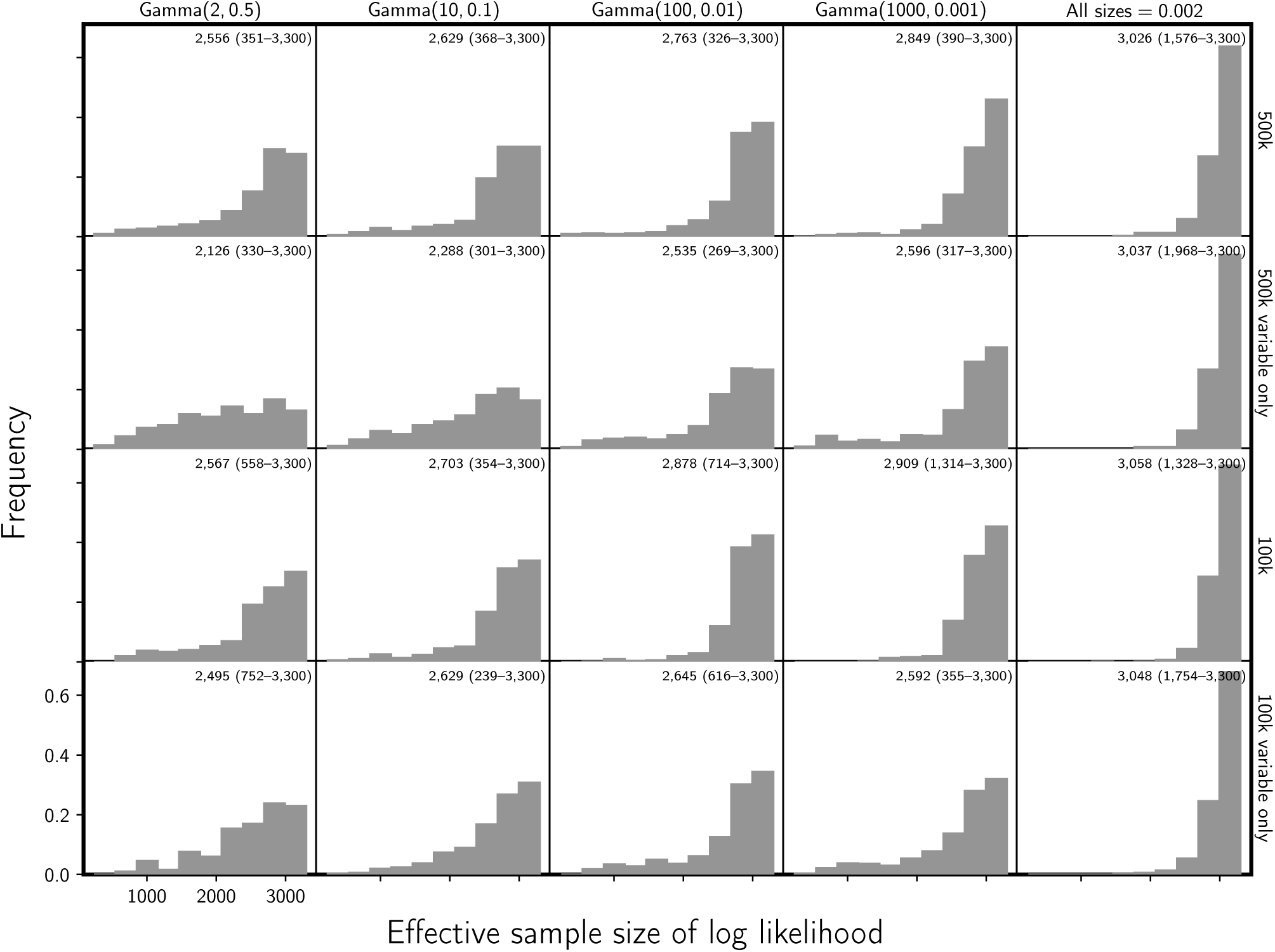
Histogram of the estimated effective sample sizes (Gong and Flegal, 2016) for the log likelihood across the three MCMC chains run for each simulated data set. The first four columns show the results from different distributions on the relative effective size of the ancestral population, decreasing in variance from left to right. The fifth column shows results when the effective size (*N*_*e*_*µ*) of all populations is fixed to 0.002. For the first two and last two rows, the simulated character matrix for each population had 500,000 and 100,000 characters, respectively. The first and third rows show the results of analyses using all characters, whereas the second and fourth rows show the results when only variable characters are used. The mean and range across the 500 data sets is indicated in the upper right corner of each plot. We generated the plot using matplotlib Version 2.0.0 (Hunter, 2007).

**Figure S10.**
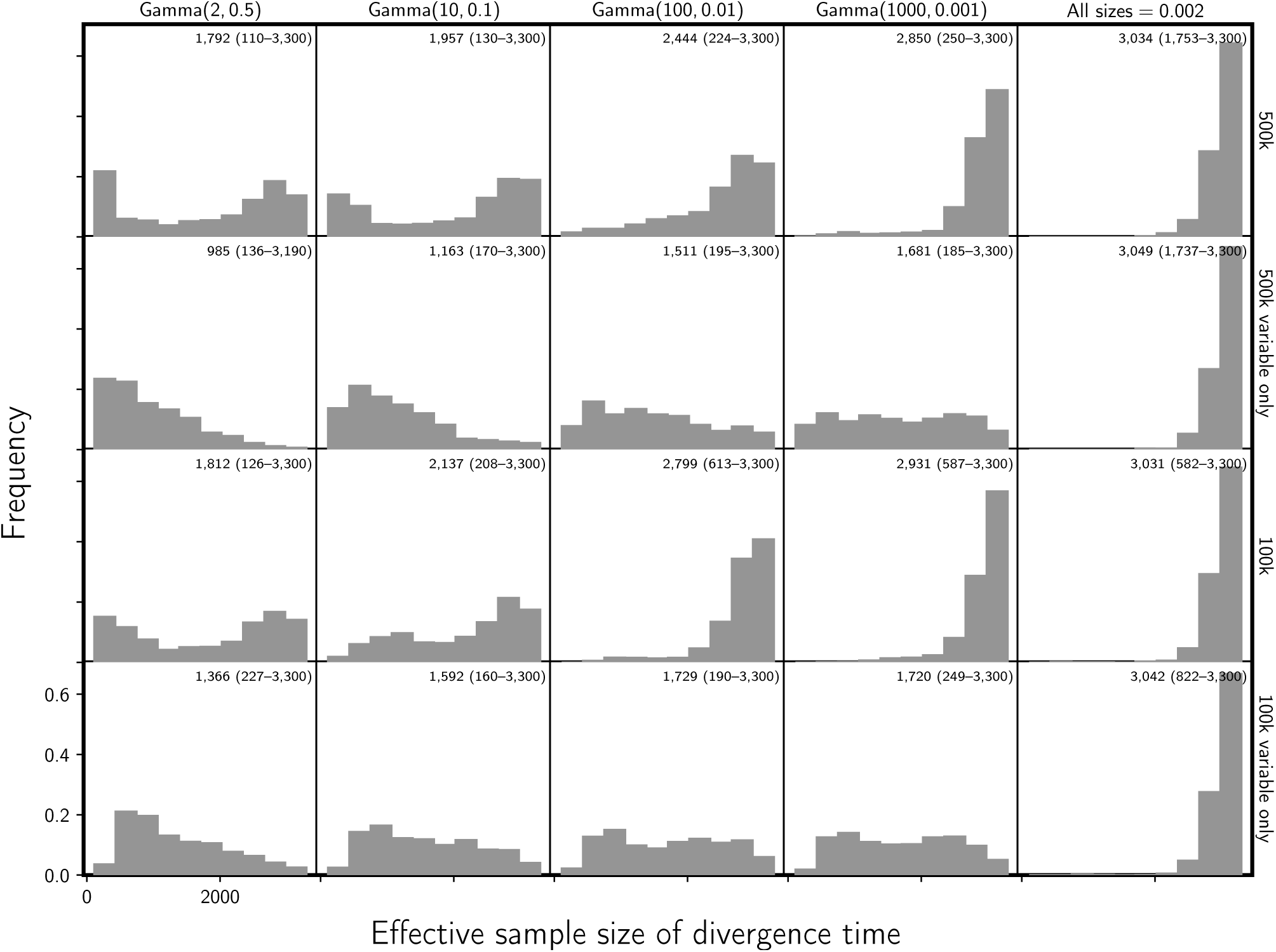
Histogram of the estimated effective sample sizes (Gong and Flegal, 2016) for the divergence times across the three MCMC chains run for each simulated data set. The first four columns show the results from different distributions on the relative effective size of the ancestral population, decreasing in variance from left to right. The fifth column shows results when the effective size (*N*_*e*_*µ*) of all populations is fixed to 0.002. For the first two and last two rows, the simulated character matrix for each population had 500,000 and 100,000 characters, respectively. The first and third rows show the results of analyses using all characters, whereas the second and fourth rows show the results when only variable characters are used. The mean and range across the 500 data sets is indicated in the upper right corner of each plot. We generated the plot using matplotlib Version 2.0.0 (Hunter, 2007).

**Figure S11.**
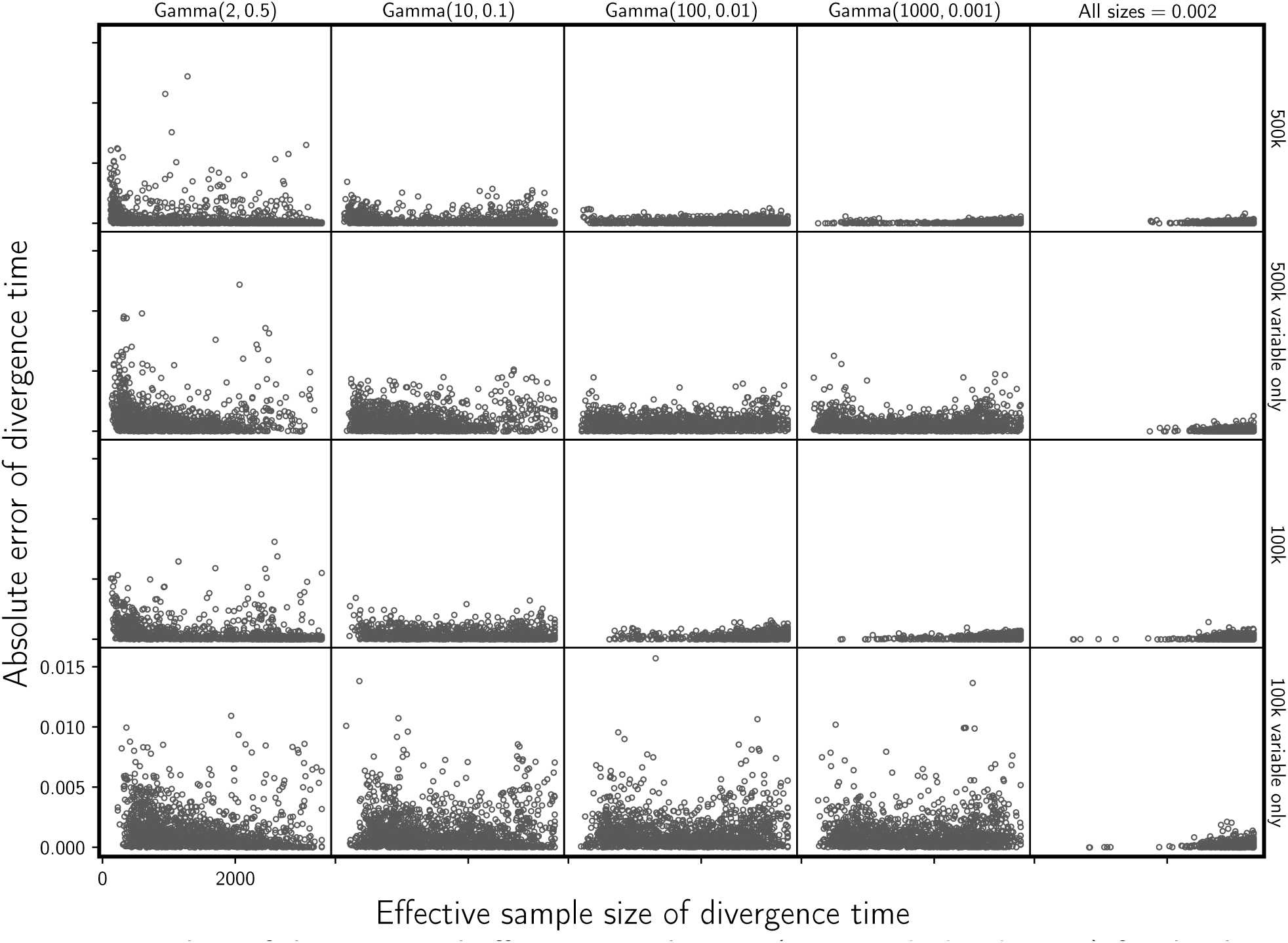
Plots of the estimated effective sample sizes (Gong and Flegal, 2016) for the divergence times against absolute error of the divergence-time estimates. The first four columns show the results from different distributions on the relative effective size of the ancestral population, decreasing in variance from left to right. The fifth column shows results when the effective size (*N*_*e*_*µ*) of all populations is fixed to 0.002. For the first two and last two rows, the simulated character matrix for each population had 500,000 and 100,000 characters, respectively. The first and third rows show the results of analyses using all characters, whereas the second and fourth rows show the results when only variable characters are used. We generated the plot using matplotlib Version 2.0.0 (Hunter, 2007).

**Figure S12.**
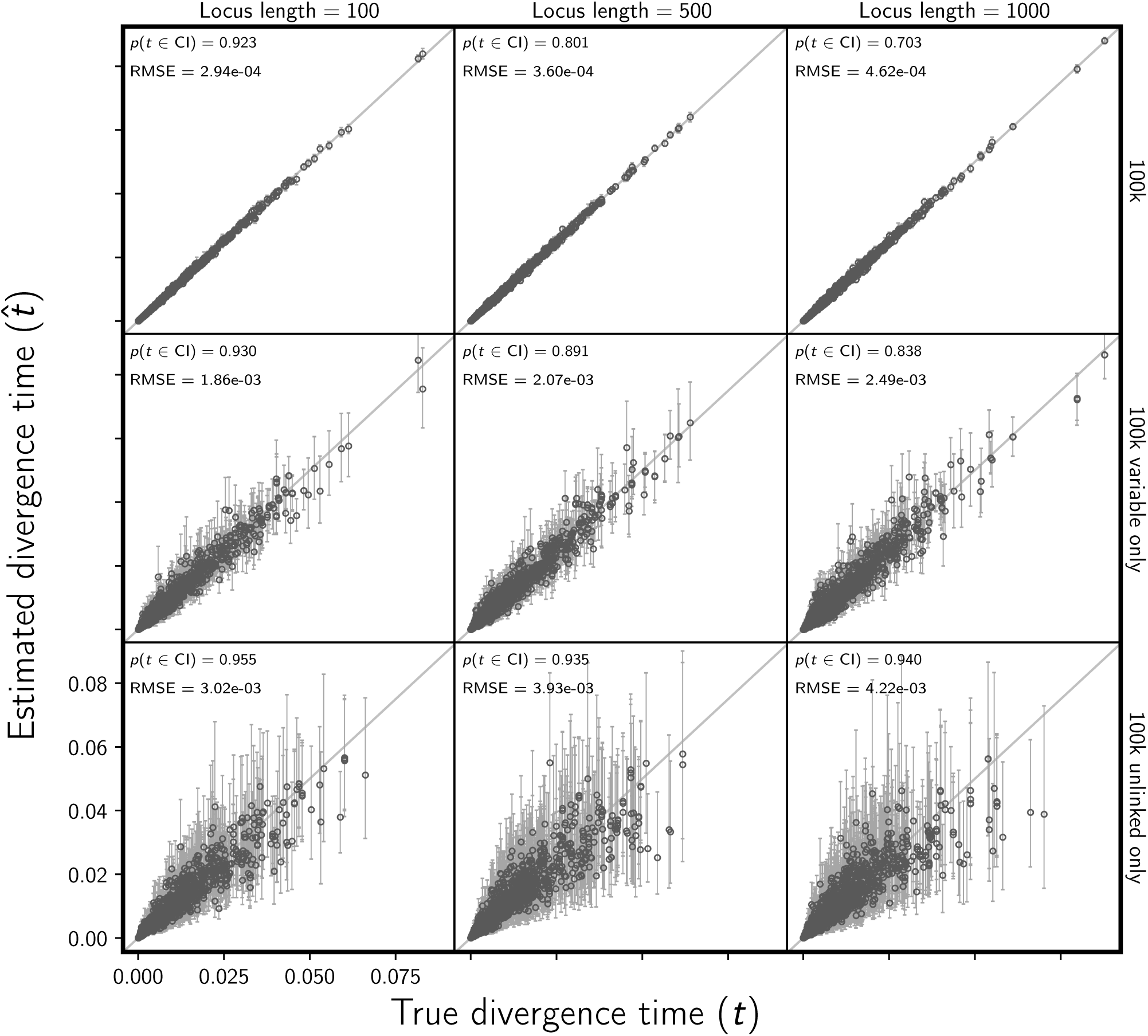
Assessing the effect of linked sites on the the accuracy and precision of divergence time estimates (in units of expected subsitutions per site). The columns, from left to right, show the results when loci are simulated with 100, 500, and 1000 linked sites. For each simulated data set, each of three population pairs has 100,000 sites total. The rows show the results when (top) all sites, (middle) all variable sites, and (bottom) at most one variable site per locus are analyzed. For each plot, the root-mean-square error (RMSE) and the proportion of estimates for which the 95% credible interval contained the true value—*p*(*t∈*CI)—is given. We generated the plot using matplotlib Version 2.0.0 (Hunter, 2007).

**Figure S13.**
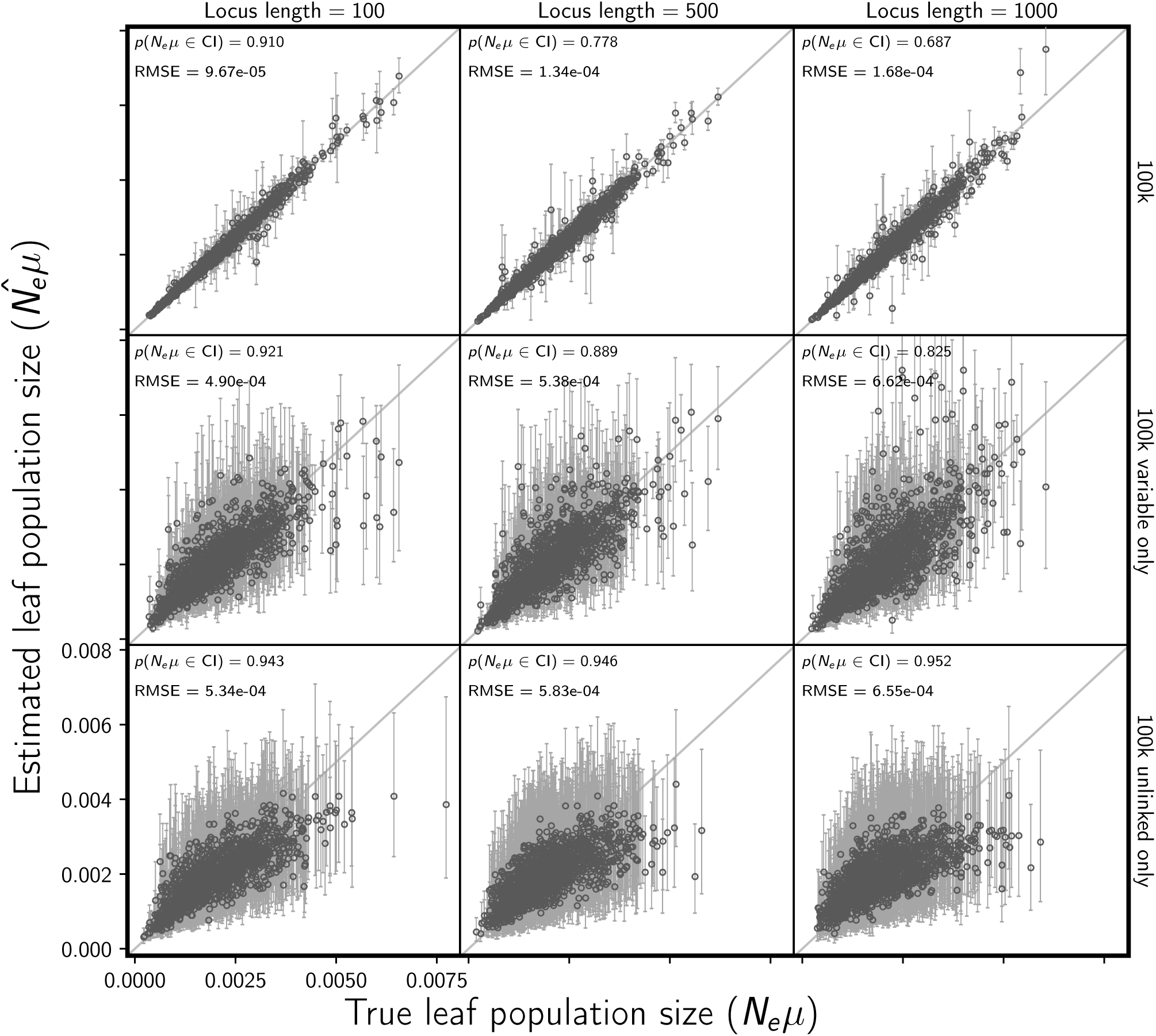
Assessing the effect of linked sites on the accuracy and precision of estimates of descendant (leaf) population sizes (scaled by the mutation rate). The columns, from left to right, show the results when loci are simulated with 100, 500, and 1000 linked sites. For each simulated data set, each of three population pairs has 100,000 sites total. The rows show the results when (top) all sites, (middle) all variable sites, and (bottom) at most one variable site per locus are analyzed. For each plot, the root-mean-square error (RMSE) and the proportion of estimates for which the 95% credible interval contained the true value—*p*(*N*_*e*_*µ∈*CI)—is given. We generated the plot using matplotlib Version 2.0.0 (Hunter, 2007).

**Figure S14.**
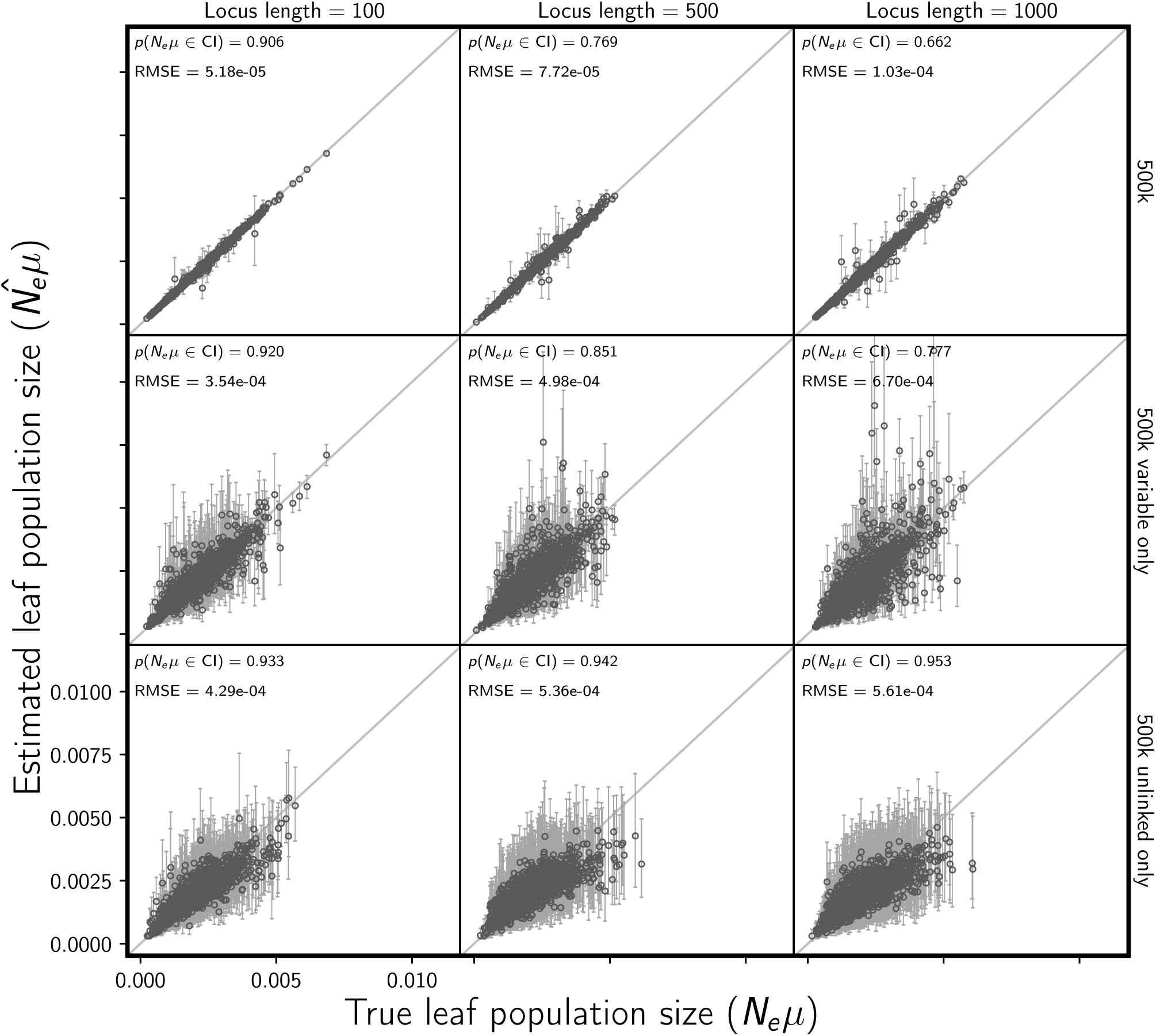
Assessing the effect of linked sites on the accuracy and precision of estimates of descendant (leaf) population sizes (scaled by the mutation rate). The columns, from left to right, show the results when loci are simulated with 100, 500, and 1000 linked sites. For each simulated data set, each of three population pairs has 500,000 sites total. The rows show the results when (top) all sites, (middle) all variable sites, and (bottom) at most one variable site per locus are analyzed. For each plot, the root-mean-square error (RMSE) and the proportion of estimates for which the 95% credible interval contained the true value—*p*(*N*_*e*_*µ∈*CI)—is given. We generated the plot using matplotlib Version 2.0.0 (Hunter, 2007).

**Figure S15.**
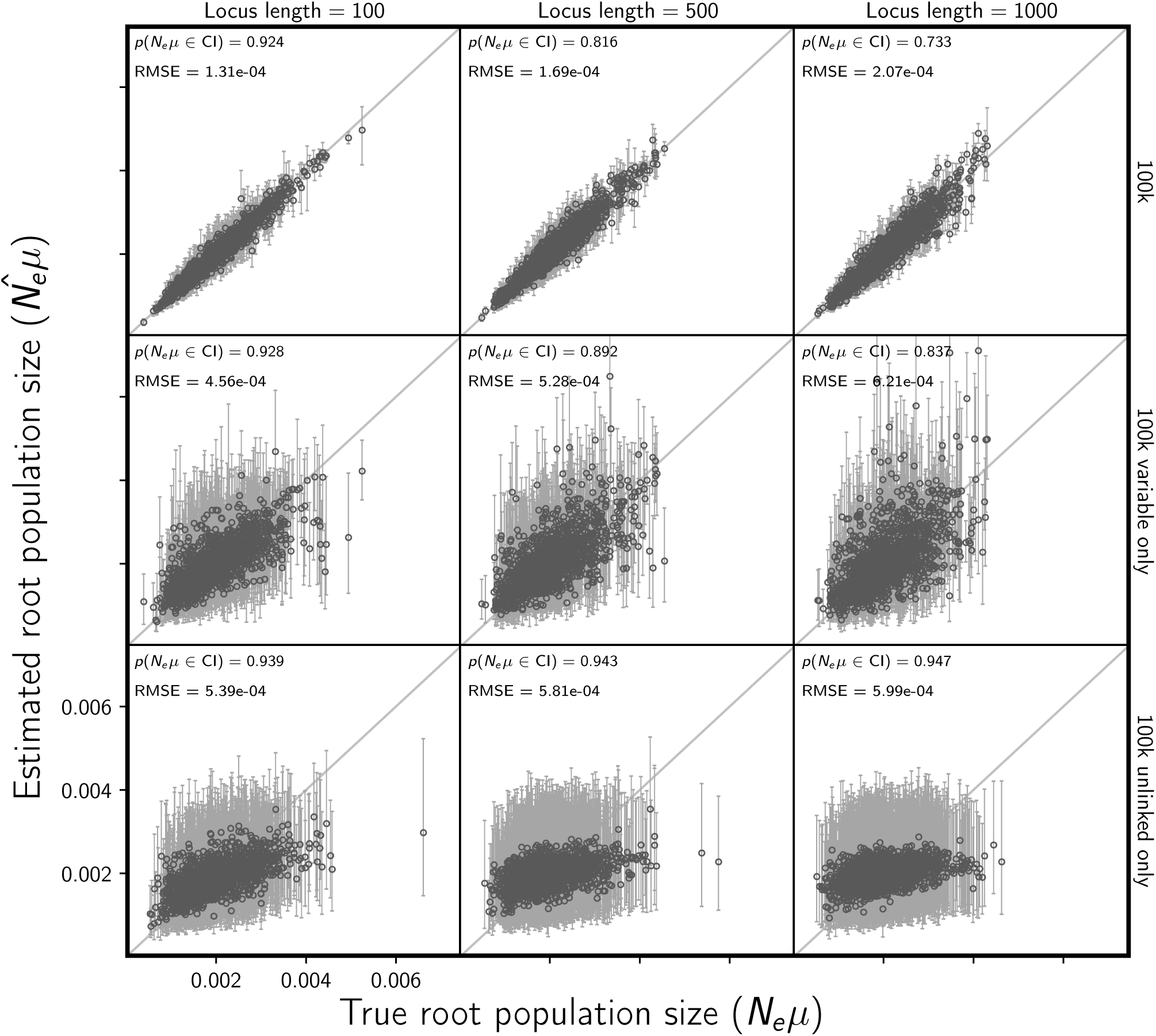
Assessing the effect of linked sites on the accuracy and precision of estimates of ancestral (root) population size (scaled by the mutation rate). The columns, from left to right, show the results when loci are simulated with 100, 500, and 1000 linked sites. For each simulated data set, each of three population pairs has 100,000 sites total. The rows show the results when (top) all sites, (middle) all variable sites, and (bottom) at most one variable site per locus are analyzed. For each plot, the root-mean-square error (RMSE) and the proportion of estimates for which the 95% credible interval contained the true value—*p*(*N*_*e*_*µ∈*CI)—is given. We generated the plot using matplotlib Version 2.0.0 (Hunter, 2007).

**Figure S16.**
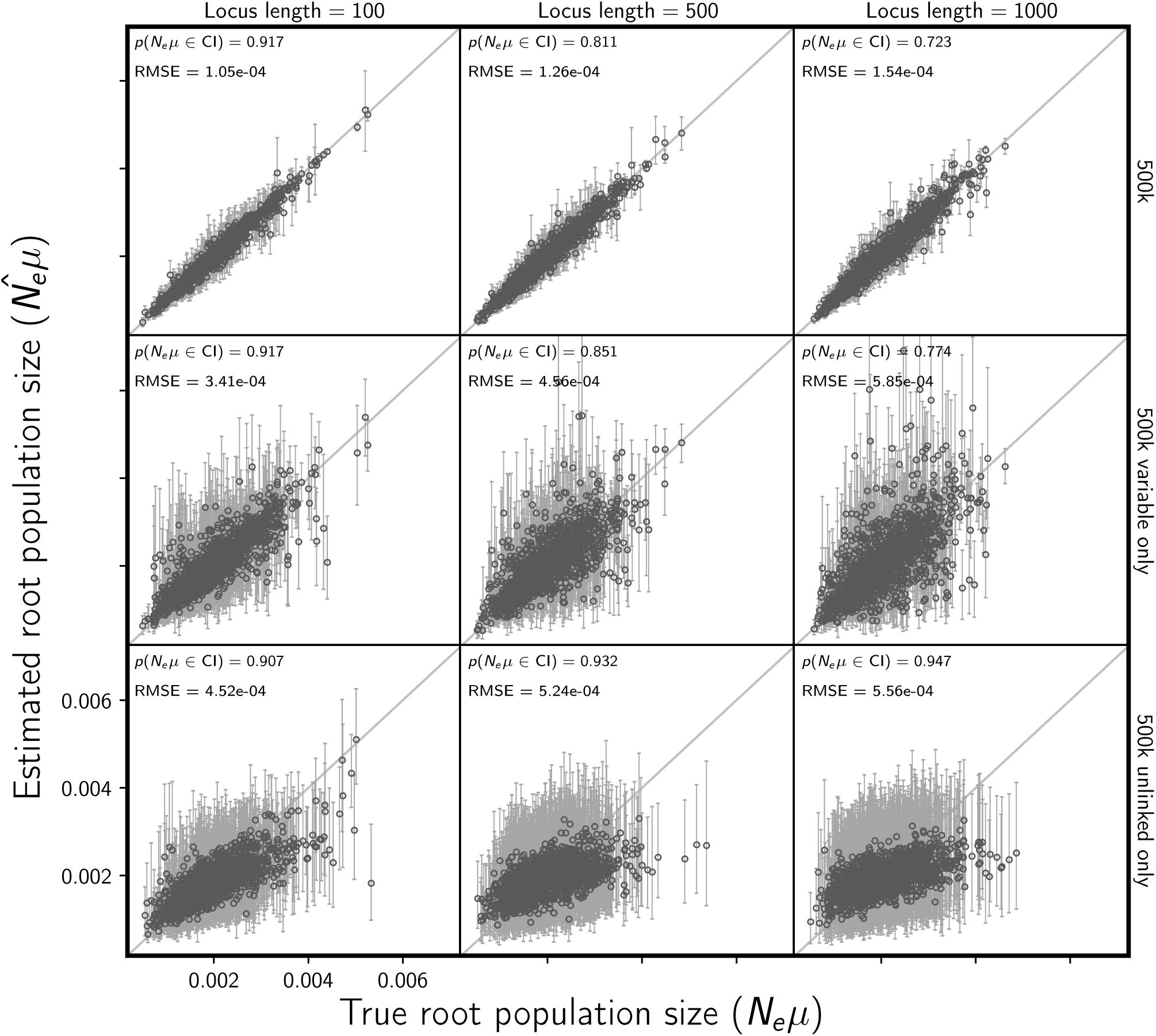
Assessing the effect of linked sites on the accuracy and precision of estimates of ancestral (root) population size (scaled by the mutation rate). The columns, from left to right, show the results when loci are simulated with 100, 500, and 1000 linked sites. For each simulated data set, each of three population pairs has 500,000 sites total. The rows show the results when (top) all sites, (middle) all variable sites, and (bottom) at most one variable site per locus are analyzed. For each plot, the root-mean-square error (RMSE) and the proportion of estimates for which the 95% credible interval contained the true value—*p*(*N*_*e*_*µ∈*CI)—is given. We generated the plot using matplotlib Version 2.0.0 (Hunter, 2007).

**Figure S17.**
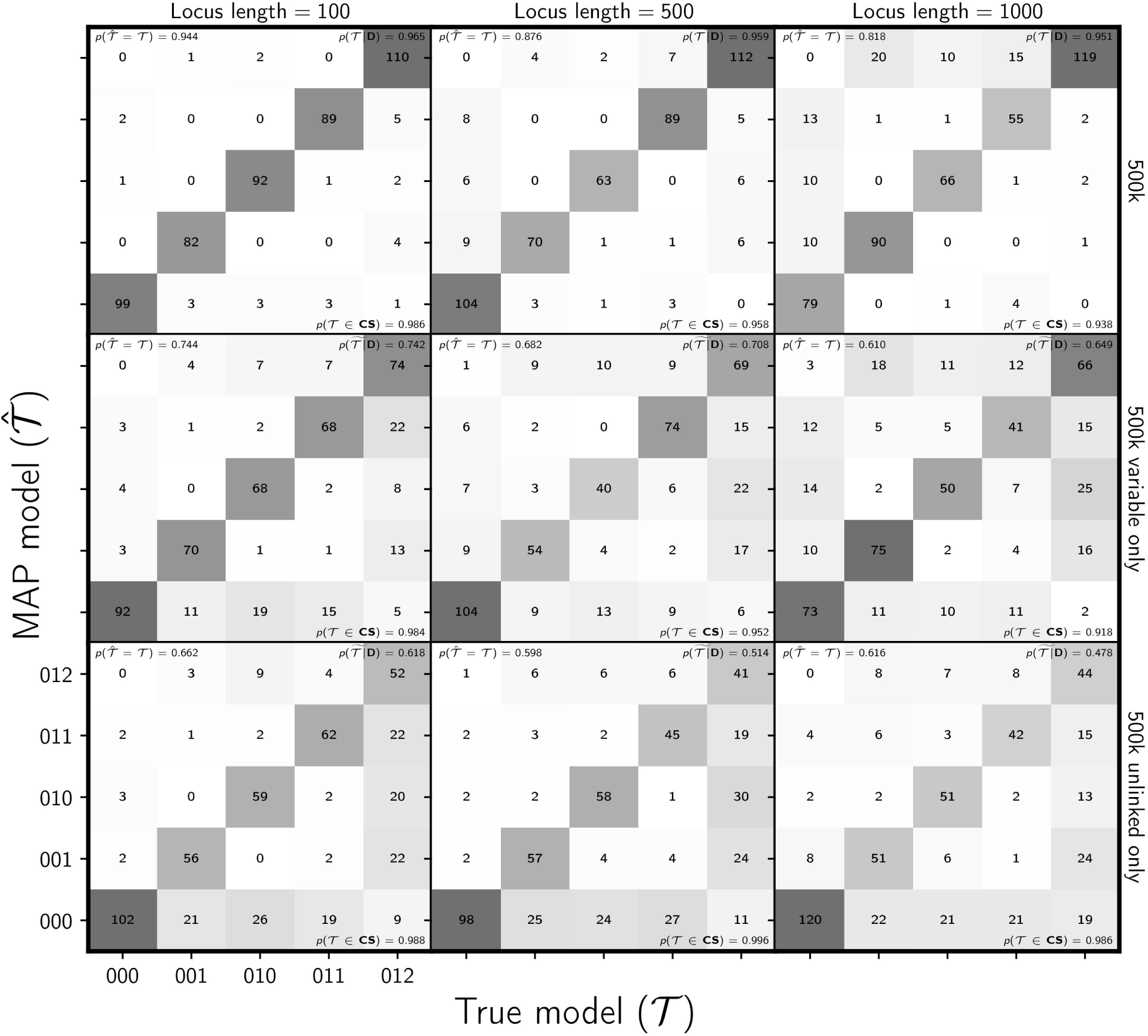
Assessing the effect of linked sites on estimating the divergence model. The columns, from left to right, show the results when loci are simulated with 100, 500, and 1000 linked sites. For each simulated data set, each of three population pairs has 500,000 sites total. The rows show the results when (top) all sites, (middle) all variable sites, and (bottom) at most one variable site per locus are analyzed. The number of data sets that fall within each possible cell of true versus estimated model is shown, and cells with more data sets are shaded darker. Each model is represented along the plot axes by three integers that indicate the divergence category of each pair of populations (e.g., 011 represents the model in which the second and third pair diverge at the same time, but separately from the first). The estimates are based on the model with the maximum *a posteriori* (MAP) probability. For each plot, the proportion of data sets for which the model with the largest posterior probability matched the true model—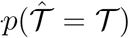—is shown in the upper left corner, the median posterior probability of the correct model across all data sets—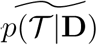—is shown in the upper right corner, and the proportion of data sets for which the true model was included in the 95% credible set—*p*(𝒯*∈*CS)—is shown in the lower right. We generated the plot using matplotlib Version 2.0.0 (Hunter, 2007).

**Figure S18.**
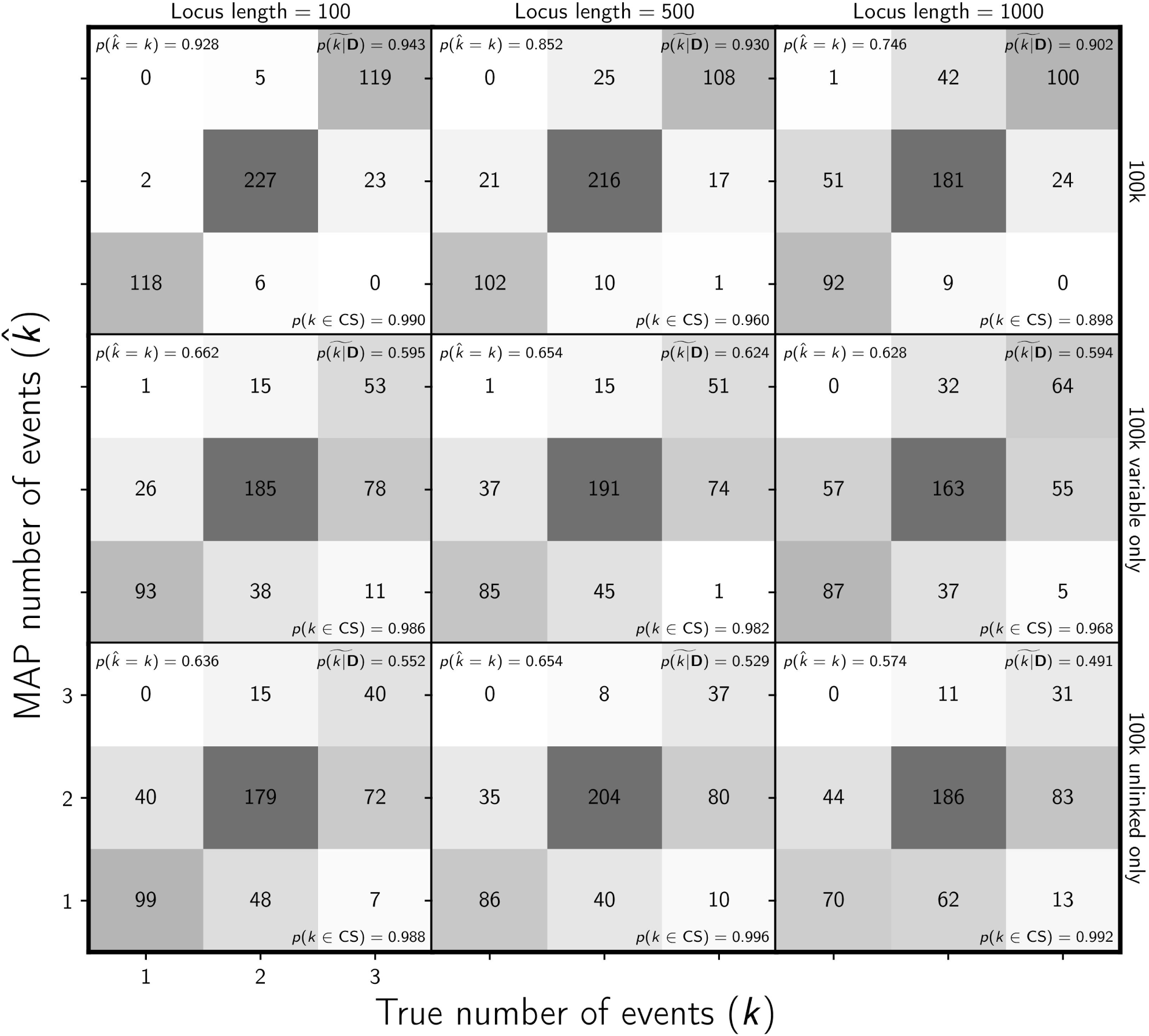
Assessing the effect of linked sites on estimating the number of divergence events. The columns, from left to right, show the results when loci are simulated with 100, 500, and 1000 linked sites. For each simulated data set, each of three population pairs has 100,000 sites total. The rows show the results when (top) all sites, (middle) all variable sites, and (bottom) at most one variable site per locus are analyzed. The number of data sets that fall within each possible cell of true versus estimated numbers of events is shown, and cells with more data sets are shaded darker. The estimates are based on the number of events with the maximum *a posteriori* (MAP) probability. For each plot, the proportion of data sets for which the number of events with the largest posterior probability matched the true number of events—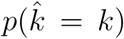—is shown in the upper left corner, the median posterior probability of the correct number of events across all data sets—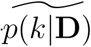—is shown in the upper right corner, and the proportion of data sets for which the true number of events was included in the 95% credible set—*p*(*k∈*CS)—is shown in the lower right. We generated the plot using matplotlib Version 2.0.0 (Hunter, 2007).

**Figure S19.**
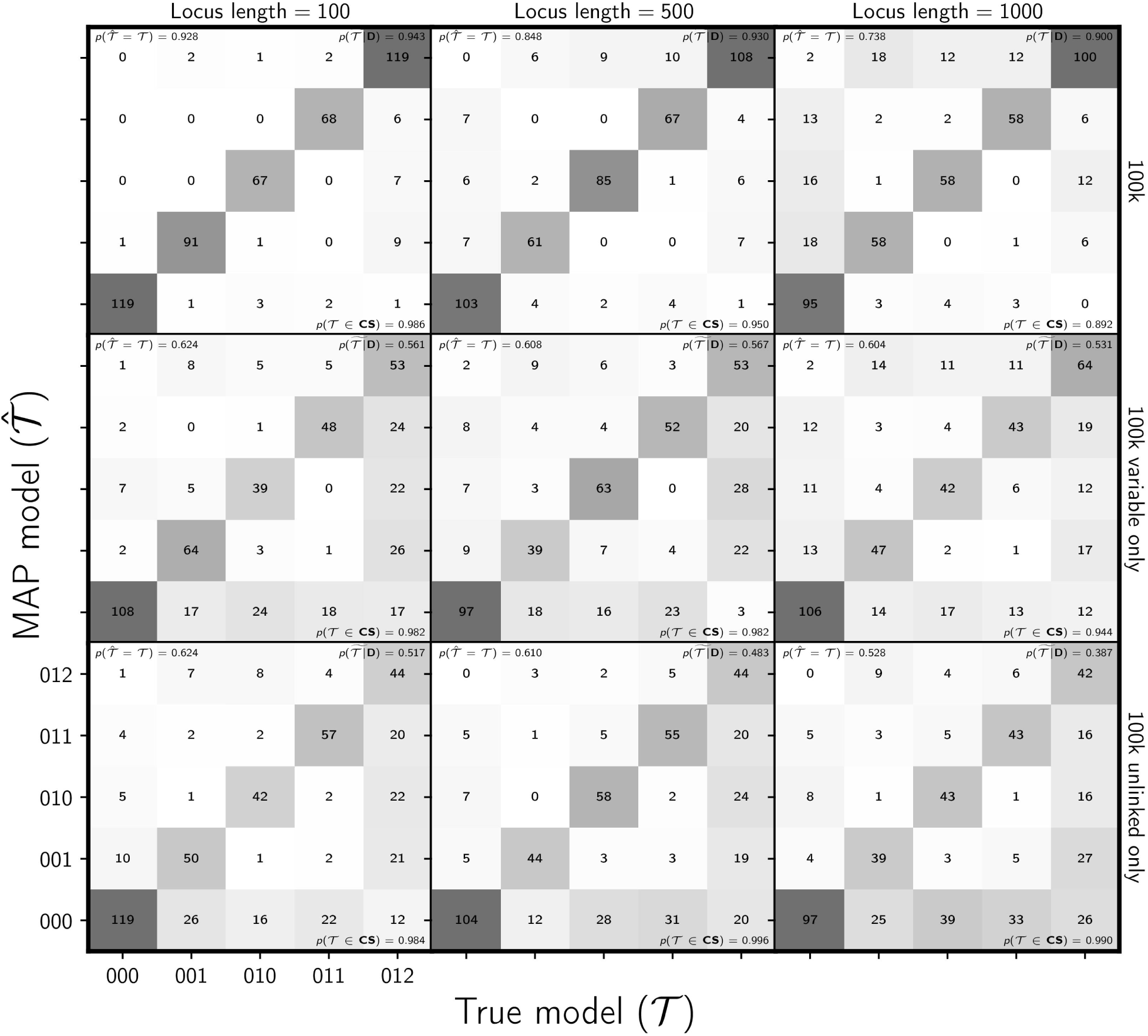
Assessing the effect of linked sites on estimating the divergence model. The columns, from left to right, show the results when loci are simulated with 100, 500, and 1000 linked sites. For each simulated data set, each of three population pairs has 100,000 sites total. The rows show the results when (top) all sites, (middle) all variable sites, and (bottom) at most one variable site per locus are analyzed. The number of data sets that fall within each possible cell of true versus estimated model is shown, and cells with more data sets are shaded darker. Each model is represented along the plot axes by three integers that indicate the divergence category of each pair of populations (e.g., 011 represents the model in which the second and third pair diverge at the same time, but separately from the first). The estimates are based on the model with the maximum *a posteriori* (MAP) probability. For each plot, the proportion of data sets for which the model with the largest posterior probability matched the true model—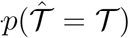—is shown in the upper left corner, the median posterior probability of the correct model across all data sets—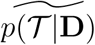—is shown in the upper right corner, and the proportion of data sets for which the true model was included in the 95% credible set—*p*(*𝒯∈*CS)—is shown in the lower right. We generated the plot using matplotlib Version 2.0.0 (Hunter, 2007).

**Figure S20.**
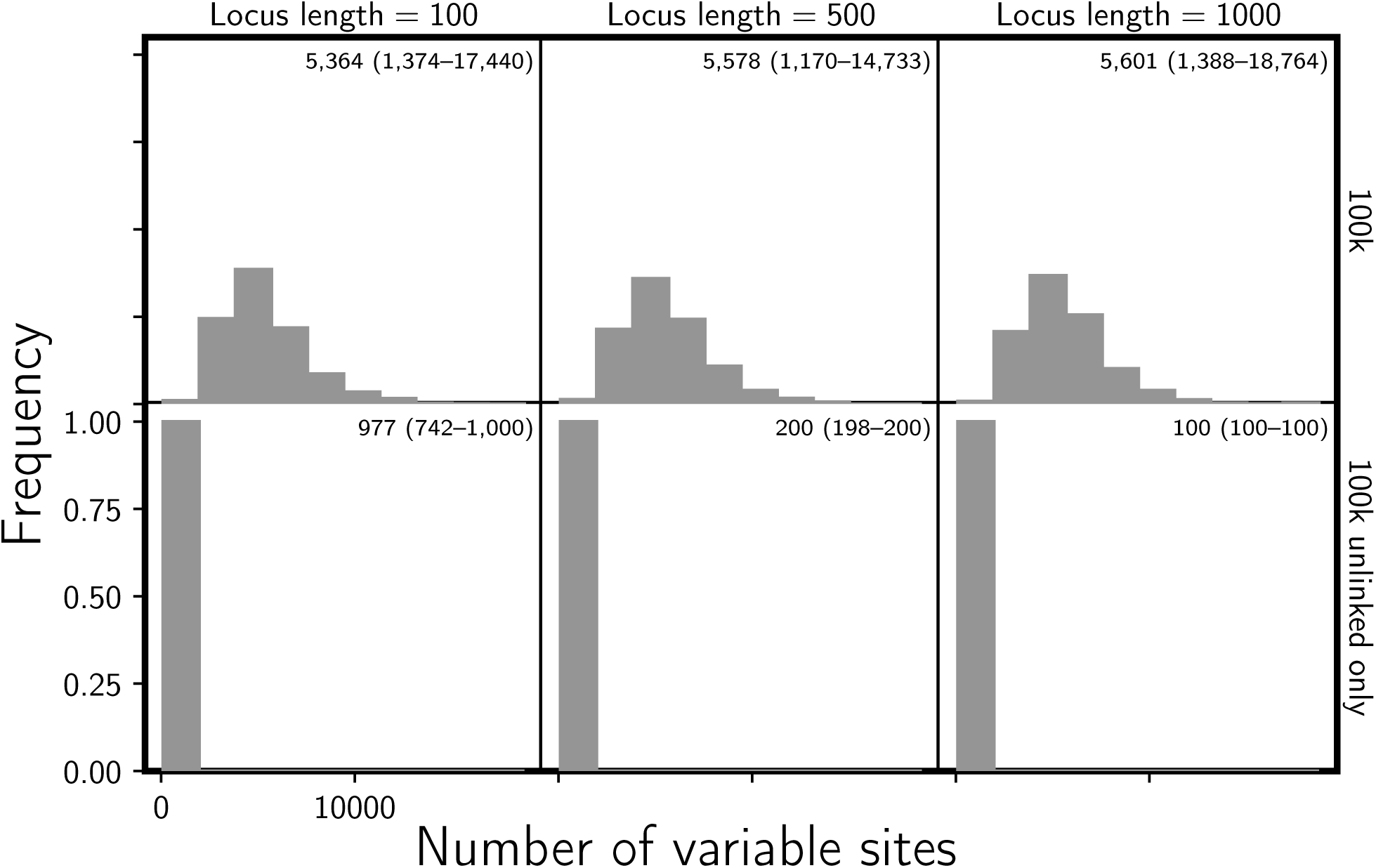
The number of variable characters in the simulated data sets with 100,000 characters per pair of populations that were linked in loci of length (left column) 100, (middle column) 500, and (right column) 1000 sites. The first row shows all the variable sites, whereas the second row shows when at most one variable site per locus is randomly choosen. The mean and range across the 500 data sets is indicated in the upper right corner of each plot. We generated the plot using matplotlib Version 2.0.0 (Hunter, 2007).

**Figure S21.**
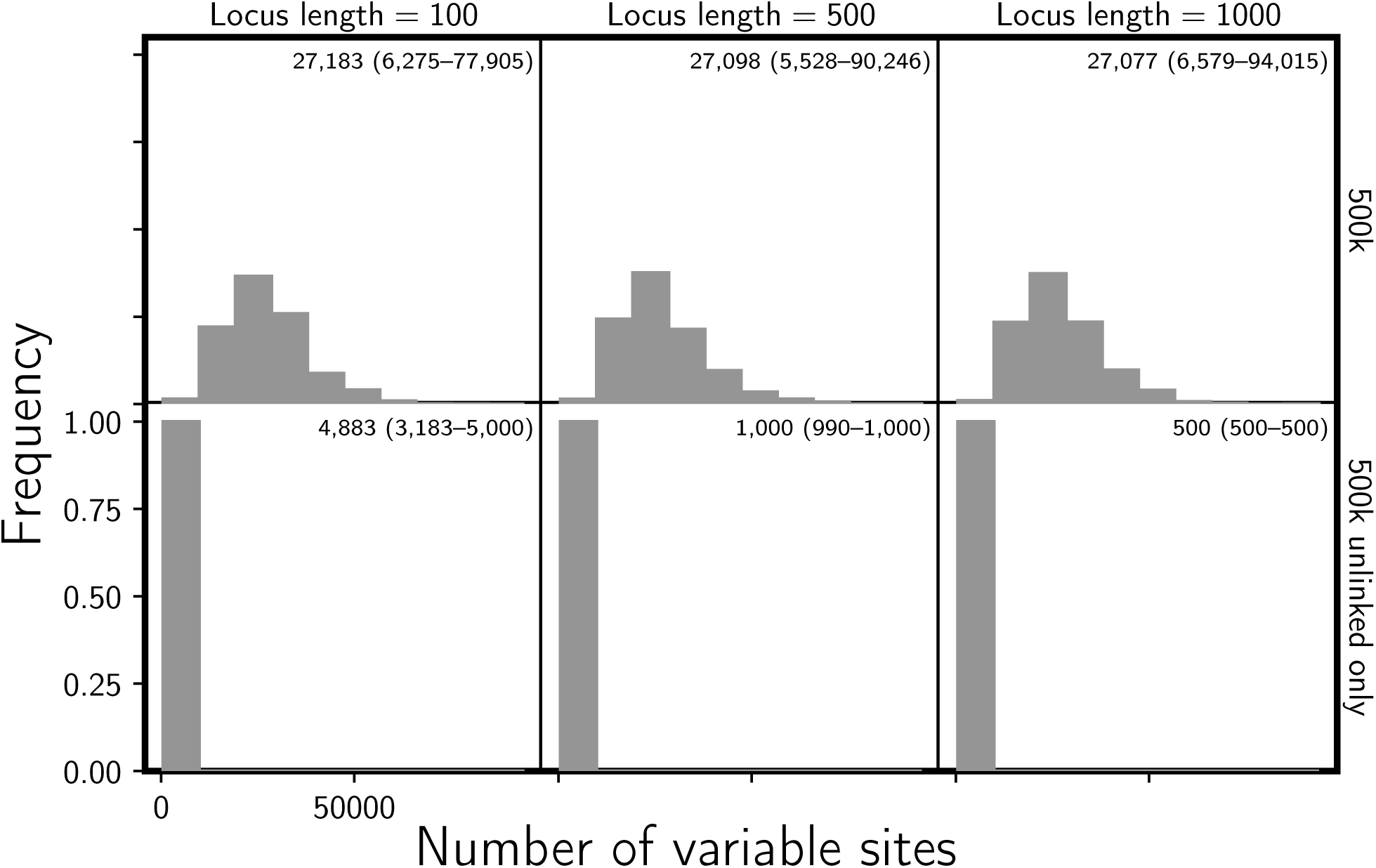
The number of variable characters in the simulated data sets with 500,000 characters per pair of populations that were linked in loci of length (left column) 100, (middle column) 500, and (right column) 1000 sites. The first row shows all the variable sites, whereas the second row shows when at most one variable site per locus is randomly choosen. The mean and range across the 500 data sets is indicated in the upper right corner of each plot. We generated the plot using matplotlib Version 2.0.0 (Hunter, 2007).

**Figure S22.**
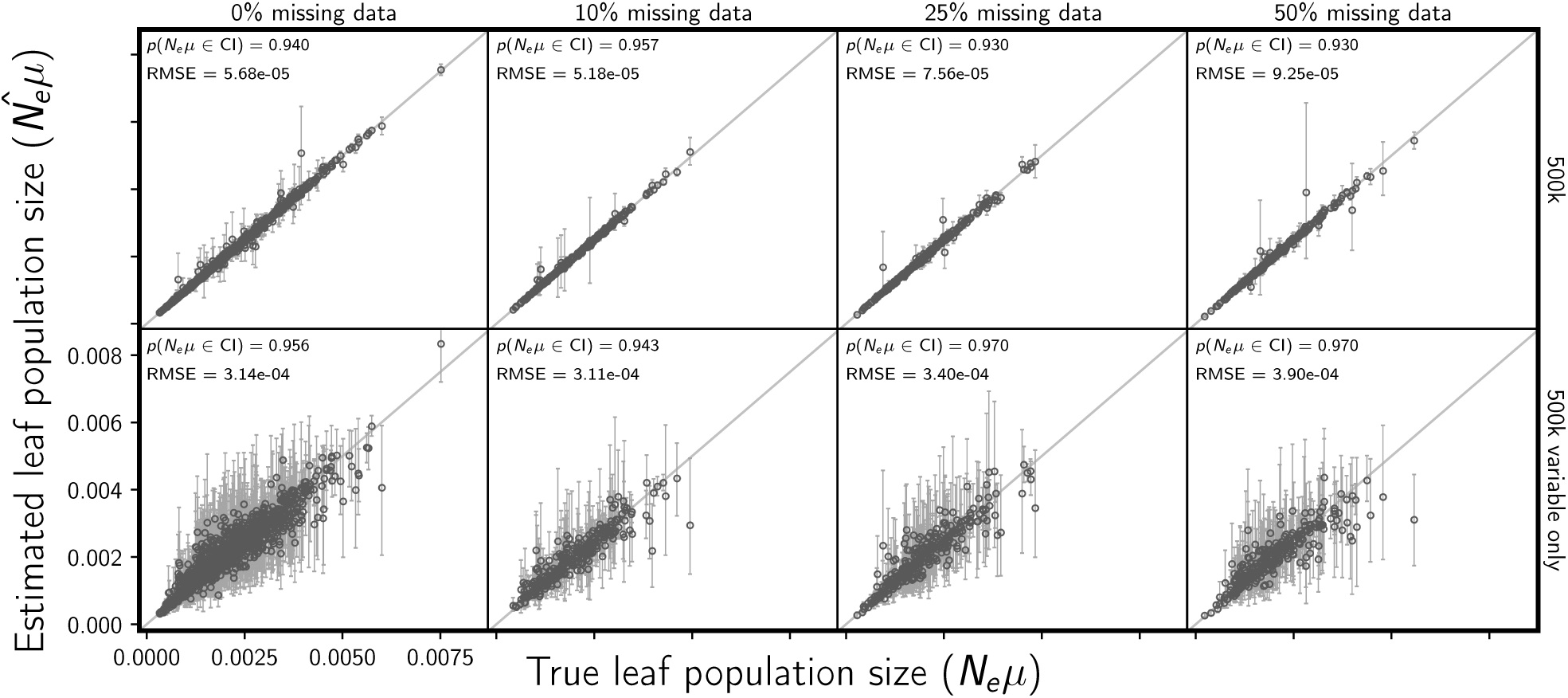
Assessing the effect of missing data on the the accuracy and precision of estimates of descendant (leaf) population sizes (scaled by the mutation rate). The columns, from left to right, show the results when each simulated 500,000-character matrix has approximately 0%, 10%, 25%, and 50% missing cells. For comparison, the first column shows the results of the 500 data sets from Figure S2; the remaining columns show the results of 100 data sets. The rows show the results when (top) all sites and (bottom) only variable sites are analyzed. For each plot, the root-mean-square error (RMSE) and the proportion of estimates for which the 95% credible interval contained the true value—*p*(*N*_*e*_*µ∈*CI)—is given. All simulated data sets had three pairs of populations. We generated the plot using matplotlib Version 2.0.0 (Hunter, 2007).

**Figure S23.**
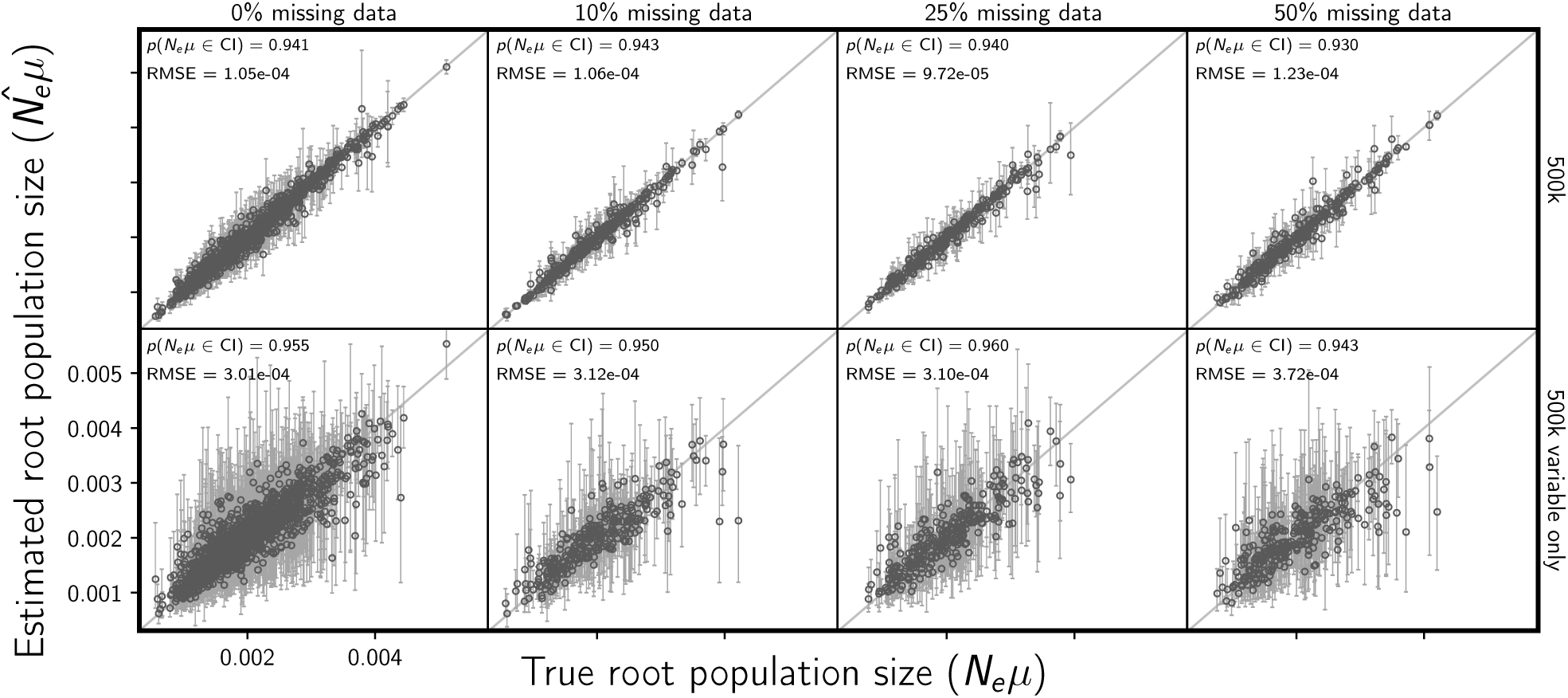
Assessing the effect of missing data on the the accuracy and precision of estimating ancestral (root) population size (scaled by the mutation rate). The columns, from left to right, show the results when each simulated 500,000-character matrix has approximately 0%, 10%, 25%, and 50% missing cells. For comparison, the first column shows the results of the 500 data sets from Figure S3; the remaining columns show the results of 100 data sets. The rows show the results when (top) all sites and (bottom) only variable sites are analyzed. For each plot, the root-mean-square error (RMSE) and the proportion of estimates for which the 95% credible interval contained the true value—*p*(*N*_*e*_*µ∈*CI)—is given. All simulated data sets had three pairs of populations. We generated the plot using matplotlib Version 2.0.0 (Hunter, 2007).

**Figure S24.**
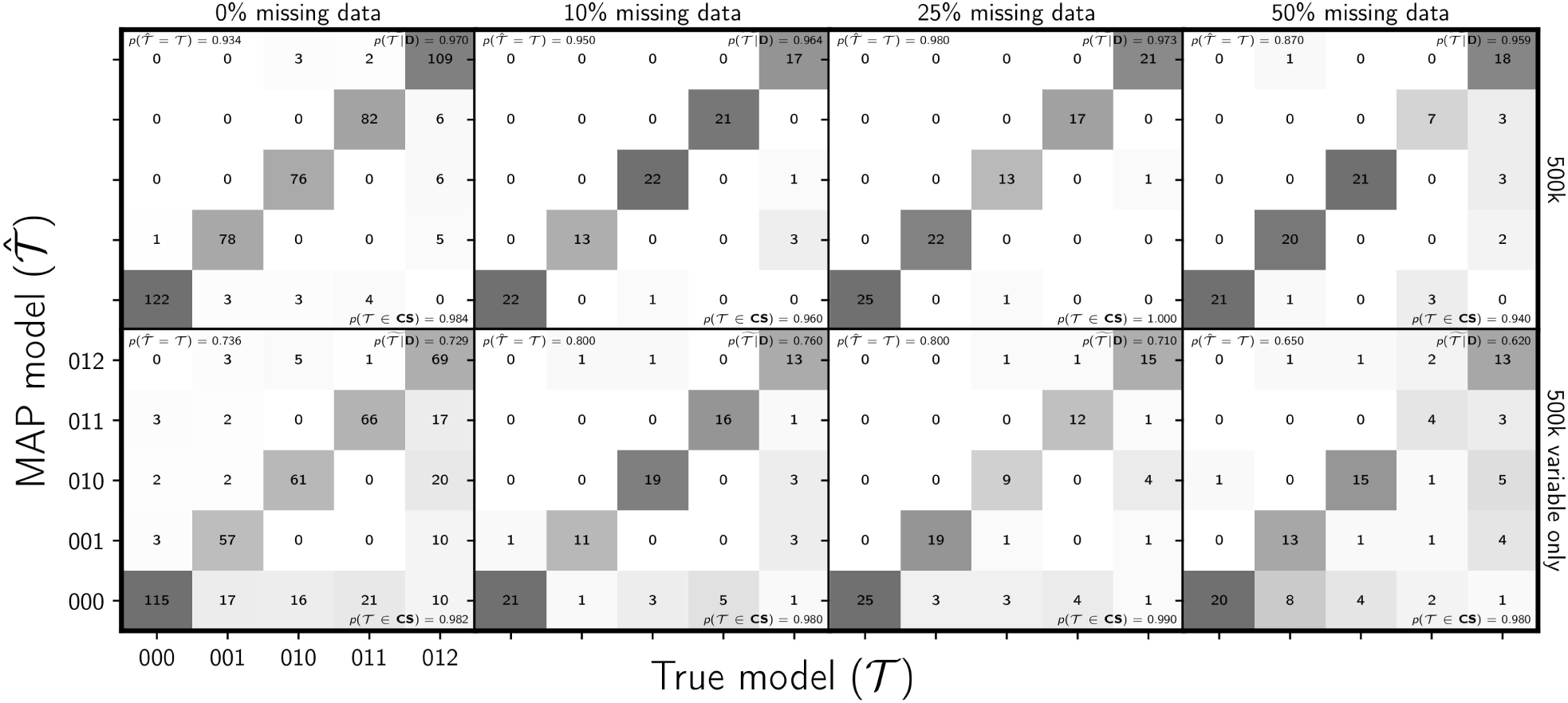
Assessing the effect of missing data on the ability of the new method to estimate the divergence model. The columns, from left to right, show the results when each simulated 500,000-character matrix has approximately 0%, 10%, 25%, and 50% missing cells. For comparison, the first column shows the results of the 500 data sets from Figure 4; the remaining columns show the results of 100 data sets. The rows show the results when (top) all sites and (bottom) only variable sites are analyzed. The number of data sets that fall within each possible cell of true versus estimated model is shown, and cells with more data sets are shaded darker. Each model is represented along the plot axes by three integers that indicate the divergence category of each pair of populations (e.g., 011 represents the model in which the second and third pair diverge at the same time, but separately from the first). The estimates are based on the model with the maximum *a posteriori* (MAP) probability. For each plot, the proportion of data sets for which the model with the largest posterior probability matched the true model—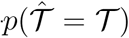—is shown in the upper left corner, the median posterior probability of the correct model across all data sets—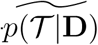—is shown in the upper right corner, and the proportion of data sets for which the true model was included in the 95% credible set—*p*(*𝒯∈*CS)—is shown in the lower right. All simulated data sets had three pairs of populations. We generated the plot using matplotlib Version 2.0.0 (Hunter, 2007).

**Figure S25.**
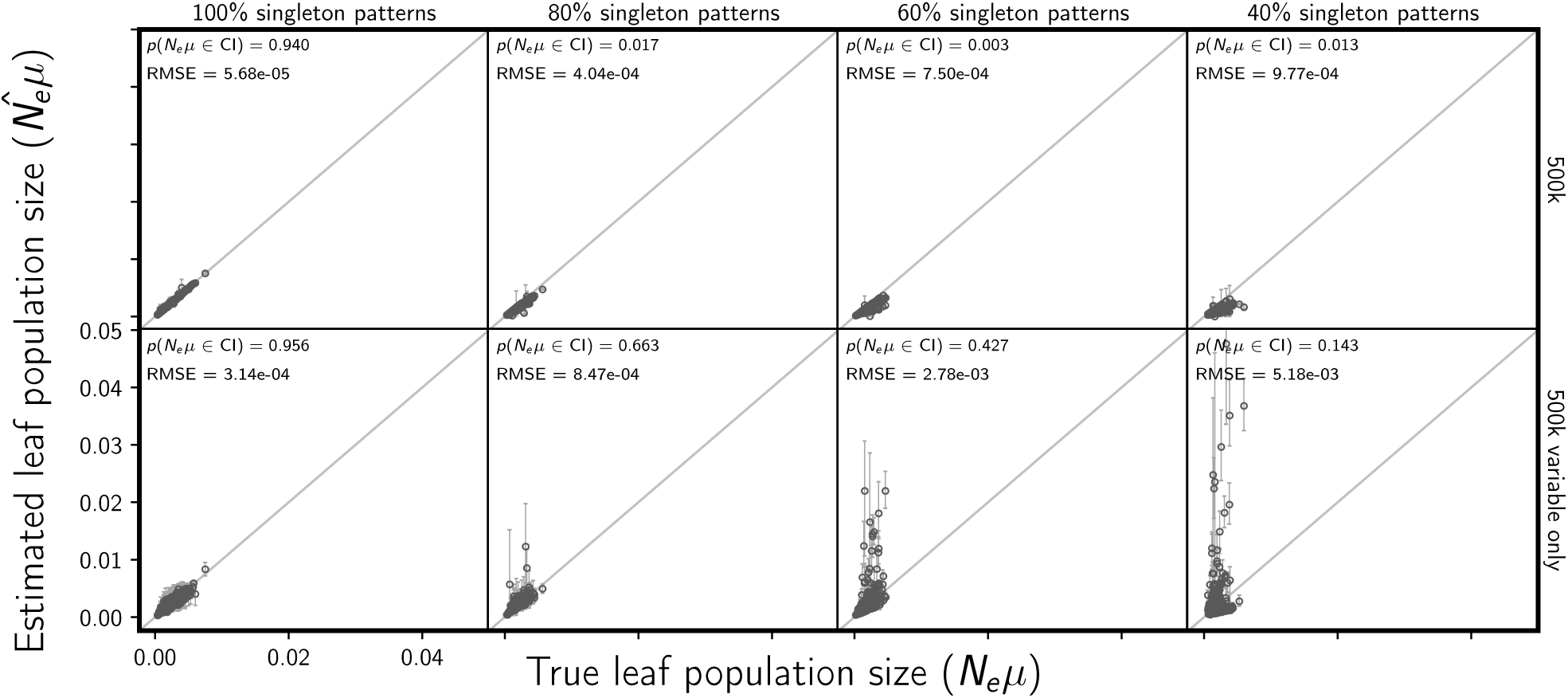
Assessing the effect of an acquisition bias against rare allele patterns on the accuracy and precision of estimating descendant (leaf) population sizes (scaled by the mutation rate). The columns, from left to right, show the results when each simulated 500,000-character data set has a probability of 100%, 80%, 60%, and 40% of sampling each simulated singleton pattern. E.g., each character matrix analyzed in the far right column is missing approximately 60% of characters where all but one gene copy has the same allele. For comparison, the first column shows the results of the 500 data sets from Figure S2; the remaining columns show the results of 100 data sets. The rows show the results when (top) all sites and (bottom) only variable sites are analyzed. For each plot, the root-mean-square error (RMSE) and the proportion of estimates for which the 95% credible interval contained the true value—*p*(*N*_*e*_*µ∈*CI)—is given. All simulated data sets had three pairs of populations. We generated the plot using matplotlib Version 2.0.0 (Hunter, 2007).

**Figure S26.**
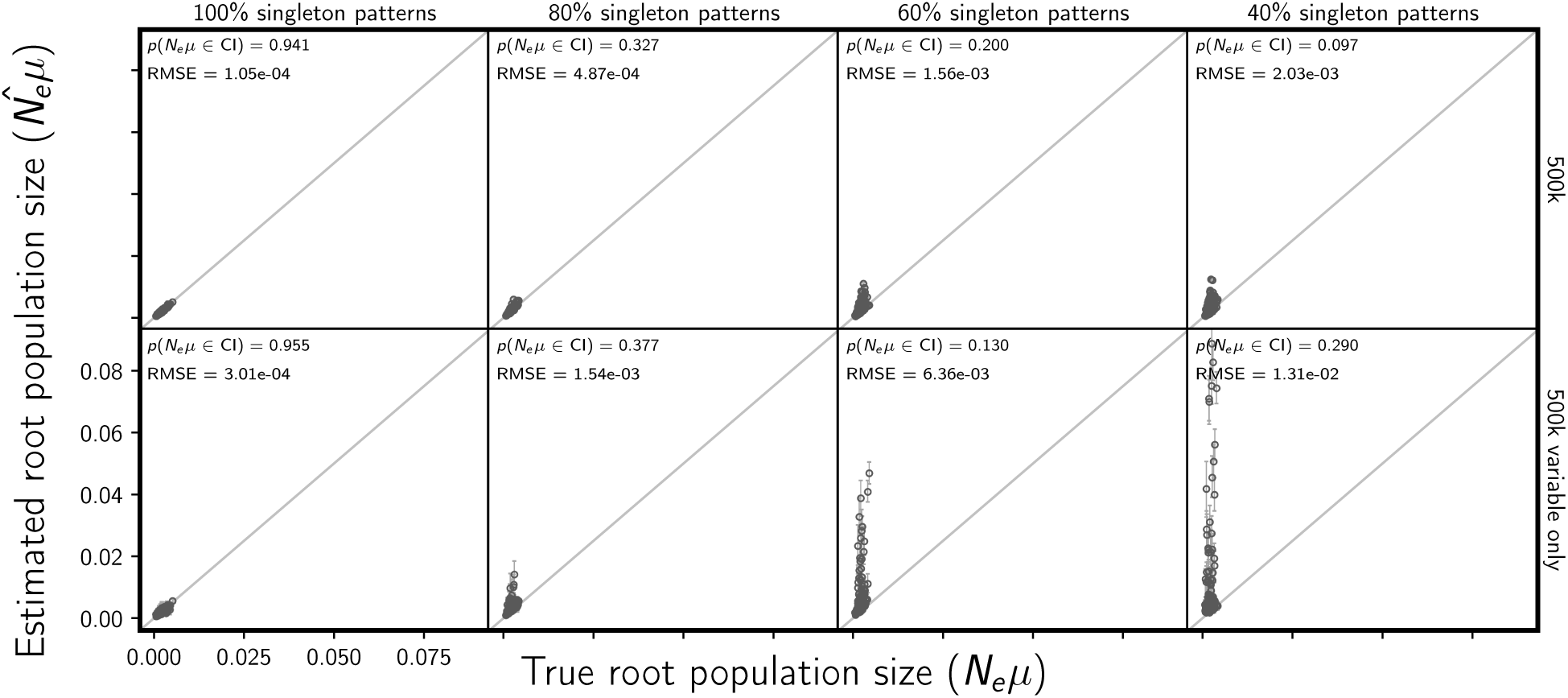
Assessing the effect of an acquisition bias against rare allele patterns on the accuracy and precision of estimating ancestral (root) population size (scaled by the mutation rate). The columns, from left to right, show the results when each simulated 500,000-character data set has a probability of 100%, 80%, 60%, and 40% of sampling each simulated singleton pattern. E.g., each character matrix analyzed in the far right column is missing approximately 60% of characters where all but one gene copy has the same allele. For comparison, the first column shows the results of the 500 data sets from Figure S3; the remaining columns show the results of 100 data sets. The rows show the results when (top) all sites and (bottom) only variable sites are analyzed. For each plot, the root-mean-square error (RMSE) and the proportion of estimates for which the 95% credible interval contained the true value—*p*(*N*_*e*_*µ∈*CI)—is given. All simulated data sets had three pairs of populations. We generated the plot using matplotlib Version 2.0.0 (Hunter, 2007).

**Figure S27.**
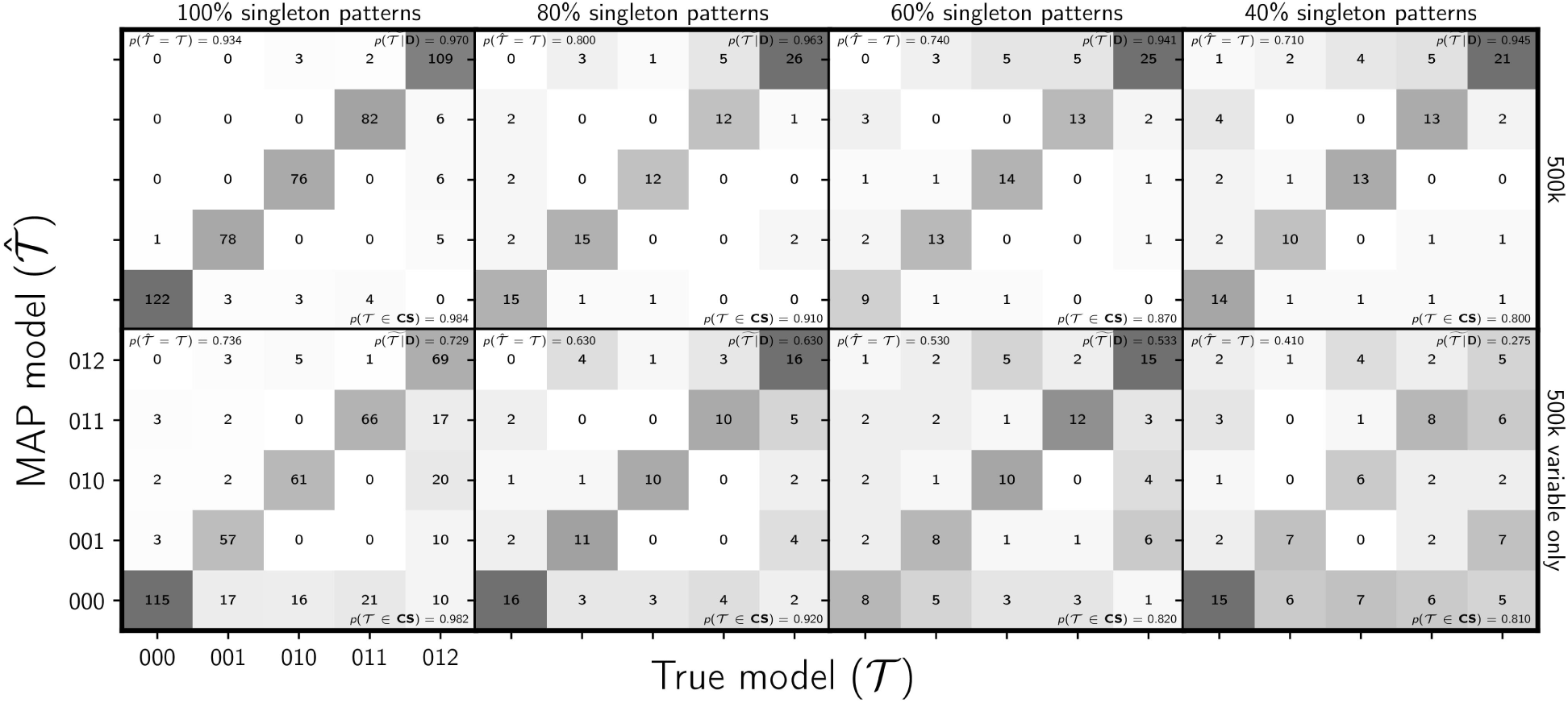
Assessing the effect of an acquisition bias against rare allele patterns on the ability of the new method to estimate the divergence model. The columns, from left to right, show the results when each simulated 500,000-character data set has a probability of 100%, 80%, 60%, and 40% of sampling each simulated singleton pattern. E.g., each character matrix analyzed in the far right column is missing approximately 60% of characters where all but one gene copy has the same allele. For comparison, the first column shows the results of the 500 data sets from Figure 4; the remaining columns show the results of 100 data sets. The rows show the results when (top) all sites and (bottom) only variable sites are analyzed. The number of data sets that fall within each possible cell of true versus estimated model is shown, and cells with more data sets are shaded darker. Each model is represented along the plot axes by three integers that indicate the divergence category of each pair of populations (e.g., 011 represents the model in which the second and third pair diverge at the same time, but separately from the first). The estimates are based on the model with the maximum *a posteriori* (MAP) probability. For each plot, the proportion of data sets for which the model with the largest posterior probability matched the true model—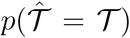—is shown in the upper left corner, the median posterior probability of the correct model across all data sets—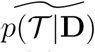—is shown in the upper right corner, and the proportion of data sets for which the true model was included in the 95% credible set— *p*(𝒯*∈*CS)—is shown in the lower right. All simulated data sets had three pairs of populations. We generated the plot using matplotlib Version 2.0.0 (Hunter, 2007).

**Figure S28.**
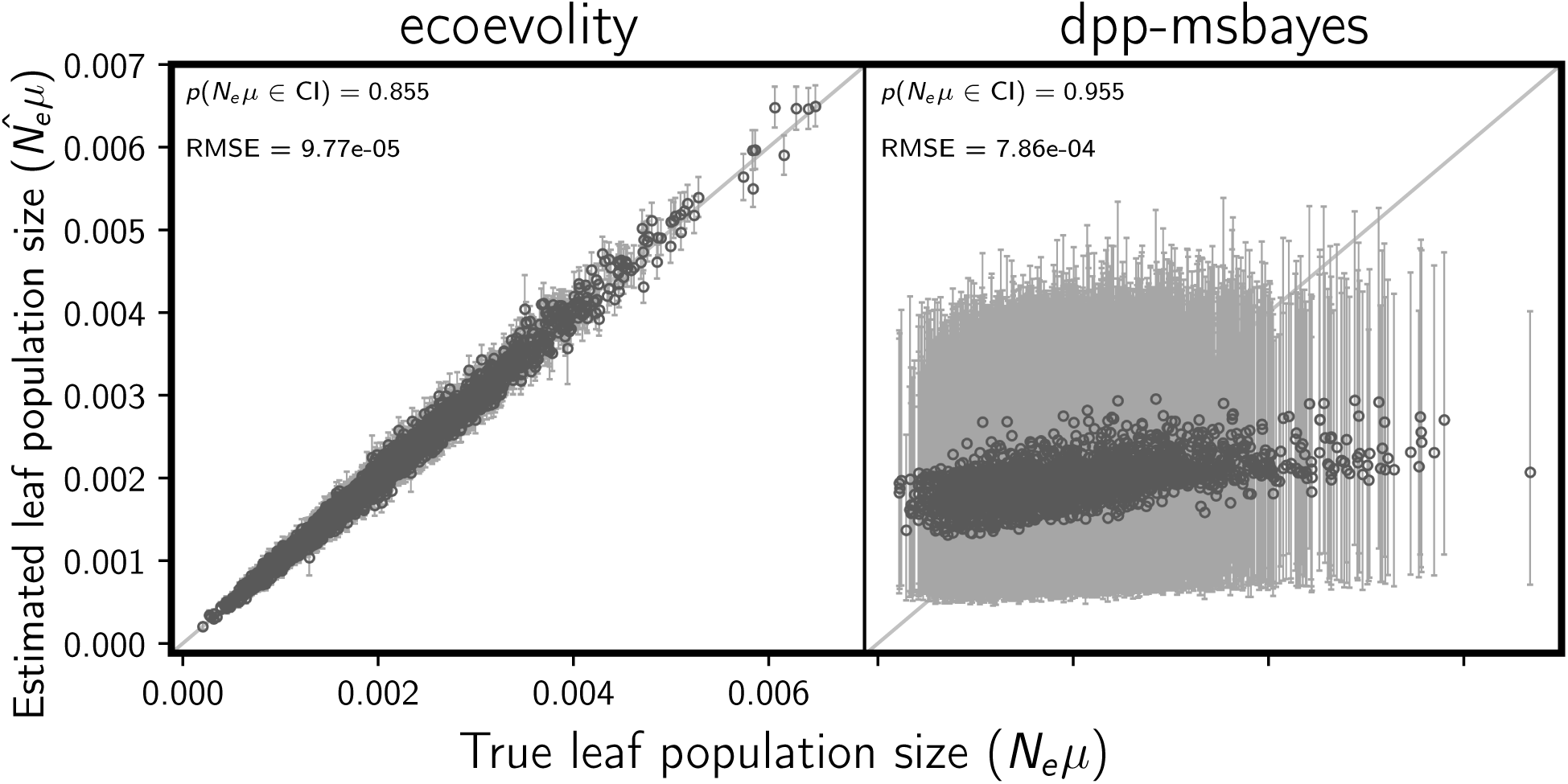
Comparing the accuracy and precision of estimates of the descendant (leaf) population sizes between (left) the new full-likelihood Bayesian method, `ecoevolity`, and (right) the approximate-likelihood Bayesian method, `dpp-msbayes`. Each plotted circle and associated error bars represent the posterior mean and 95% credible interval. Each plot consists of 3000 estimates—500 simulated data sets, each with three pairs of populations. The simulated character matrix for each population pair consisted of 200 loci, each with 200 linked sites (40,000 characters total). For each plot, the root-mean-square error (RMSE) and the proportion of estimates for which the 95% credible interval contained the true value—*p*(*N*_*e*_*µ∈*CI)—is given. We generated the plot using matplotlib Version 2.0.0 (Hunter, 2007).

**Figure S29.**
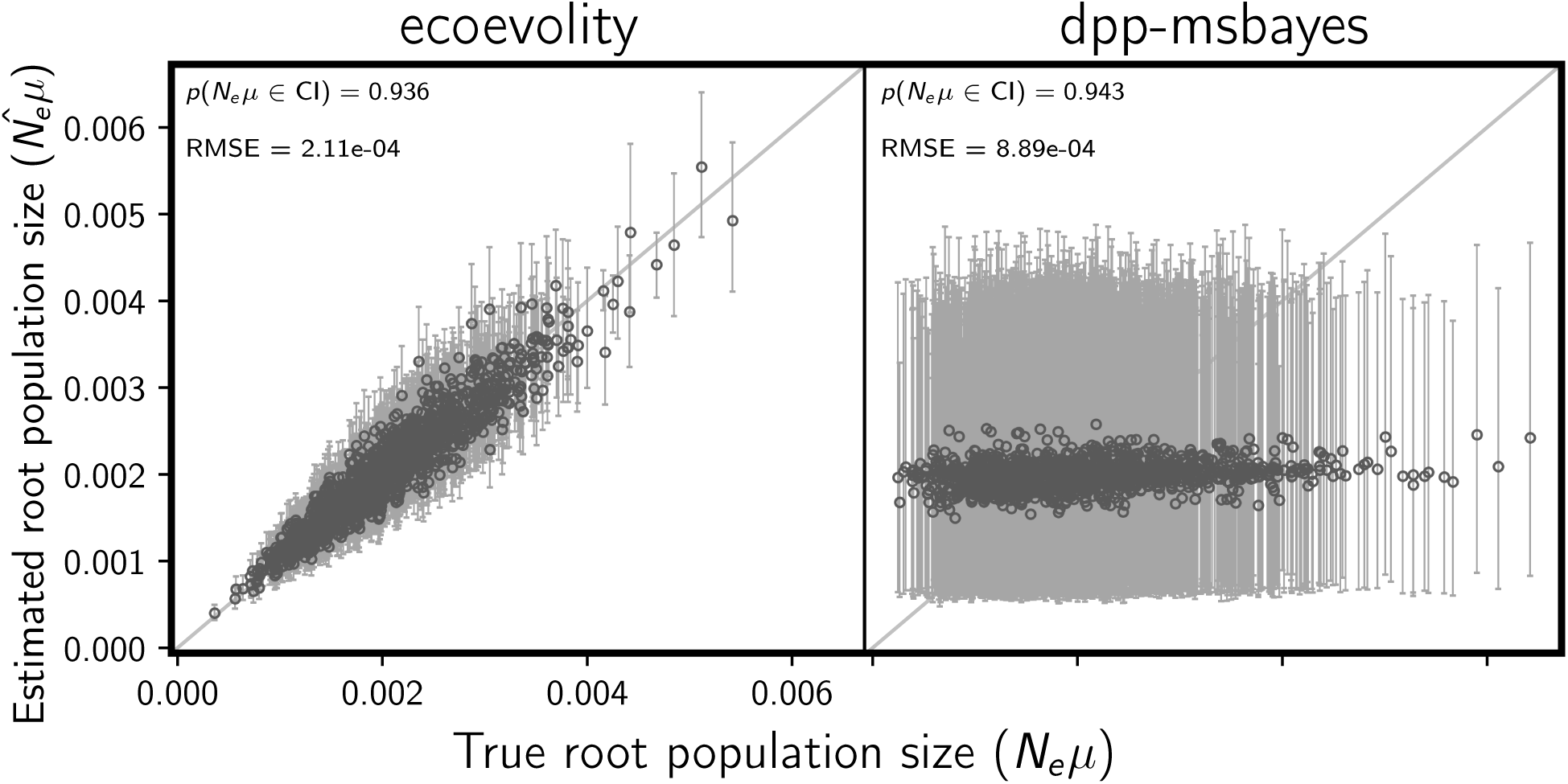
Comparing the accuracy and precision of estimates of the ancestral (root) population size between (left) the new full-likelihood Bayesian method, `ecoevolity`, and (right) the approximate-likelihood Bayesian method, `dpp-msbayes`. Each plotted circle and associated error bars represent the posterior mean and 95% credible interval. Each plot consists of 1500 estimates—500 simulated data sets, each with three pairs of populations. The simulated character matrix for each population pair consisted of 200 loci, each with 200 linked sites (40,000 characters total). For each plot, the root-mean-square error (RMSE) and the proportion of estimates for which the 95% credible interval contained the true value—*p*(*N*_*e*_*µ∈*CI)—is given. We generated the plot using matplotlib Version 2.0.0 (Hunter, 2007).

**Figure S30.**
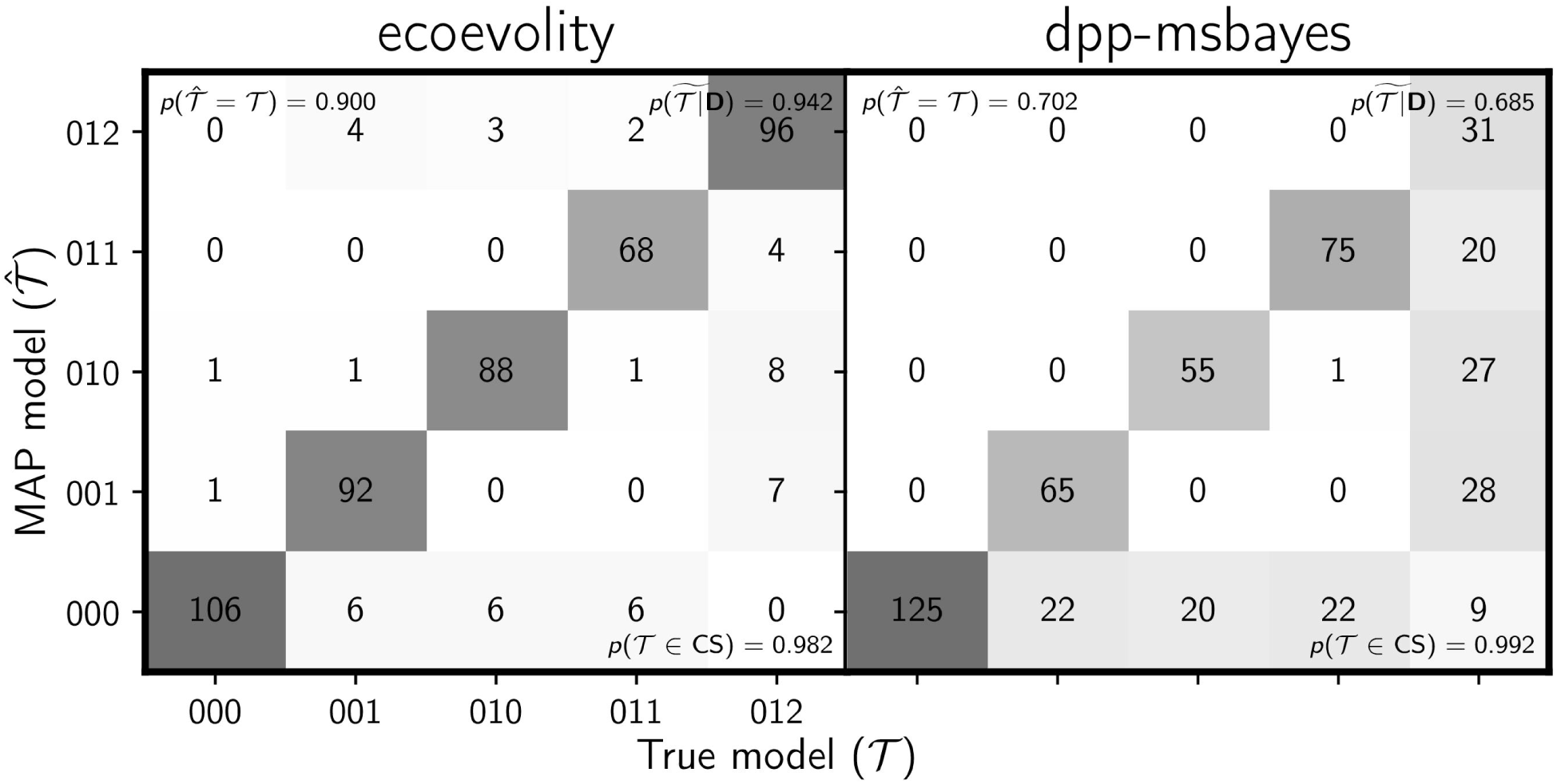
Comparing the ability to estimate the divergence model between (left) the new full-likelihood Bayesian method, `ecoevolity`, and (right) the approximate-likelihood Bayesian method, `dpp-msbayes`. Each plot shows the results of the analyses of 500 simulated data sets; the number of data sets that fall within each possible cell of true versus estimated model is shown, and cells with more data sets are shaded darker. Each simulated data set contained three pairs of populations, and the simulated character matrix for each pair consisted of 200 loci, each with 200 linked sites (40,000 characters total). Each model is represented along the plot axes by three integers that indicate the divergence category of each pair of populations (e.g., 011 represents the model in which the second and third pair diverge at the same time, but separately from the first). The estimates are based on the model with the maximum *a posteriori* (MAP) probability. For each plot, the proportion of data sets for which the model with the largest posterior probability matched the true model—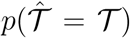—is shown in the upper left corner, the median posterior probability of the correct model across all data sets—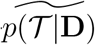—is shown in the upper right corner, and the proportion of data sets for which the true model was included in the 95% credible set— *p*(*𝒯∈*CS)—is shown in the lower right. We generated the plot using matplotlib Version 2.0.0 (Hunter, 2007).

**Figure S31.**
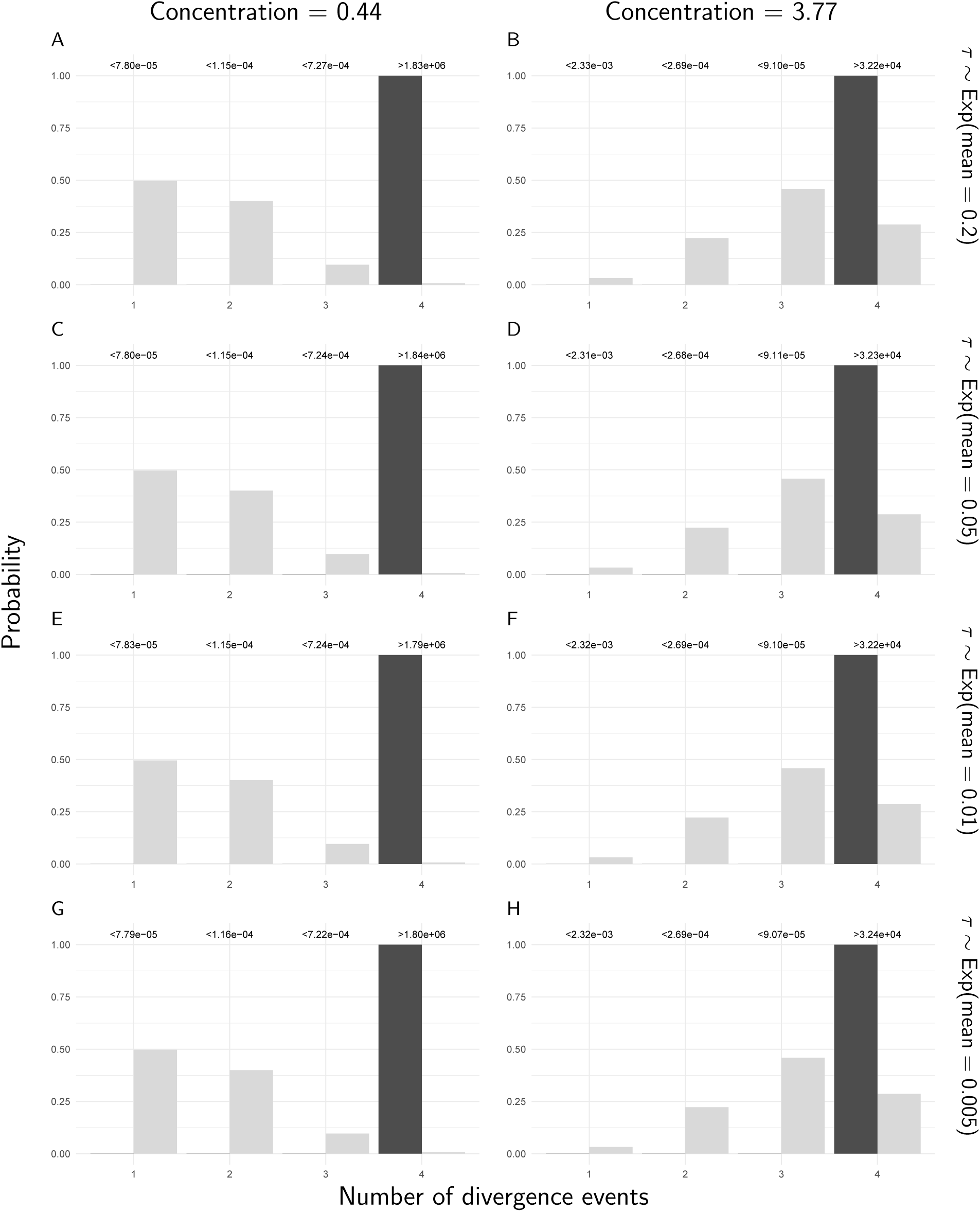
The prior (light bars) and approximated posterior (dark bars) probabilities of the number of divergence events across *Gekko* pairs of populations, under eight different combinations of prior on the divergence times (rows) and the concentration parameter of the Dirichlet process (columns). For these analyses, constant characters were included, but all characters with more than two alleles were removed. The Bayes factor for each number of divergence times is given above the corresponding bars. Each Bayes factor compares the corresponding number of events to all other possible numbers of divergence events. We generated the plots with ggplot2 Version 2.2.1 (Wickham, 2009).

**Figure S32.**
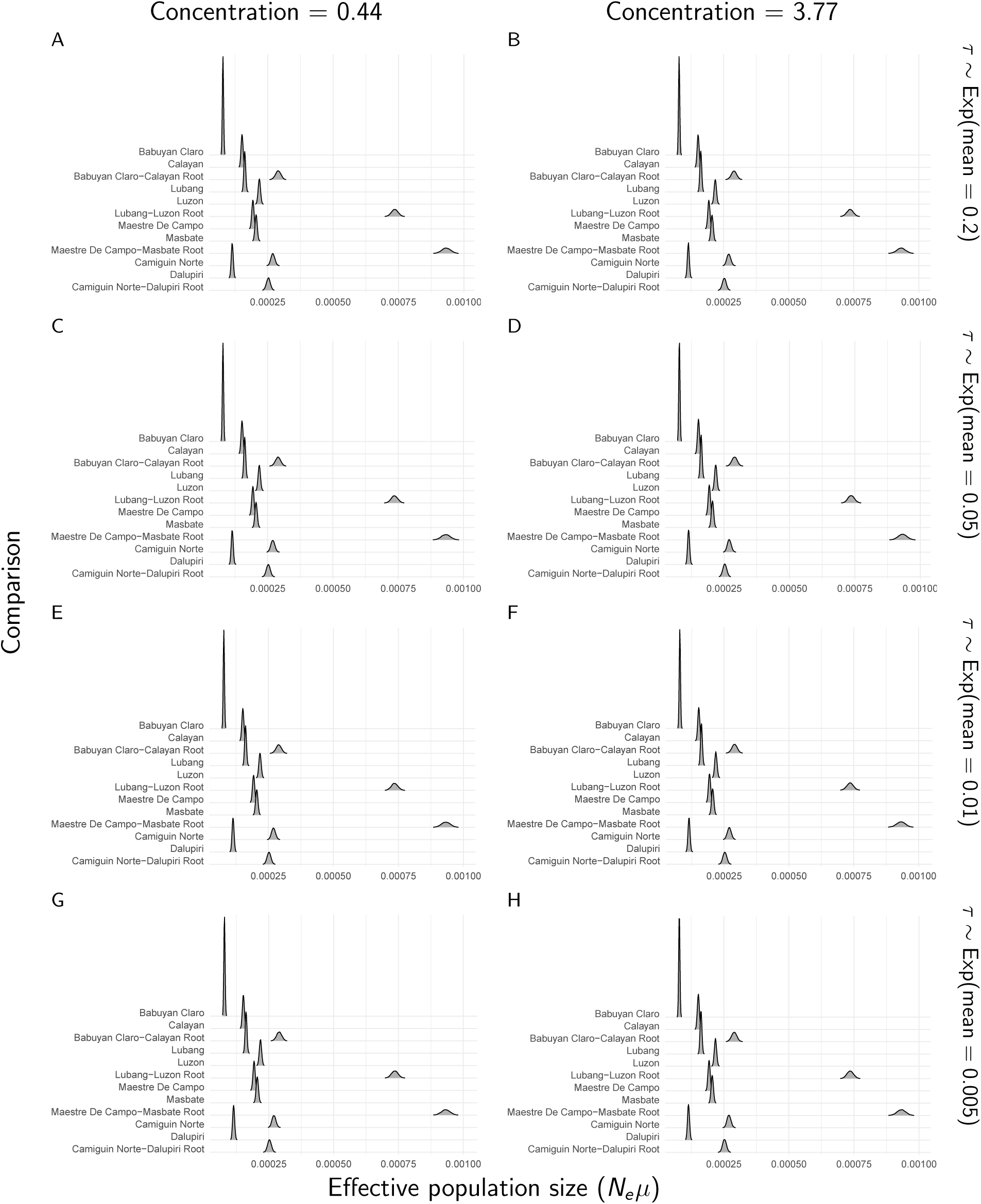
The approximate marginal posterior densities of population sizes for each *Gekko* pair of populations, under eight different combinations of prior on the divergence times (rows) and the concentration parameter of the Dirichlet process (columns). For these analyses, constant characters were included, but all characters with more than two alleles were recoded as biallelic. We generated the plots with ggridges Version 0.4.1 (Wilke, 2018) and ggplot2 Version 2.2.1 (Wickham, 2009).

**Figure S33.**
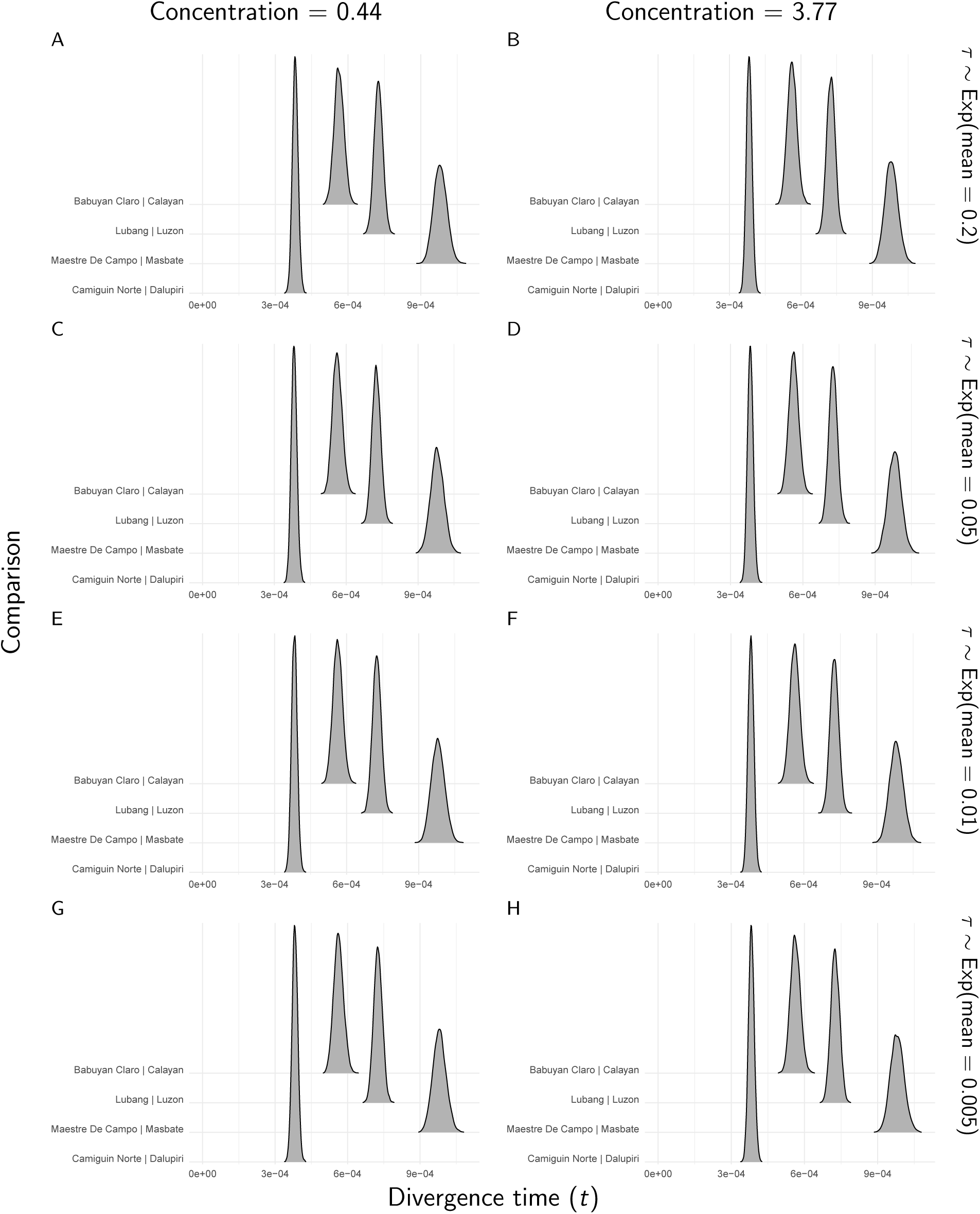
The approximate marginal posterior densities of divergence times for each *Gekko* pair of populations, under eight different combinations of prior on the divergence times (rows) and the concentration parameter of the Dirichlet process (columns). For these analyses, constant characters were included, but all characters with more than two alleles were removed. We generated the plots with ggridges Version 0.4.1 (Wilke, 2018) and ggplot2 Version 2.2.1 (Wickham, 2009).

**Figure S34.**
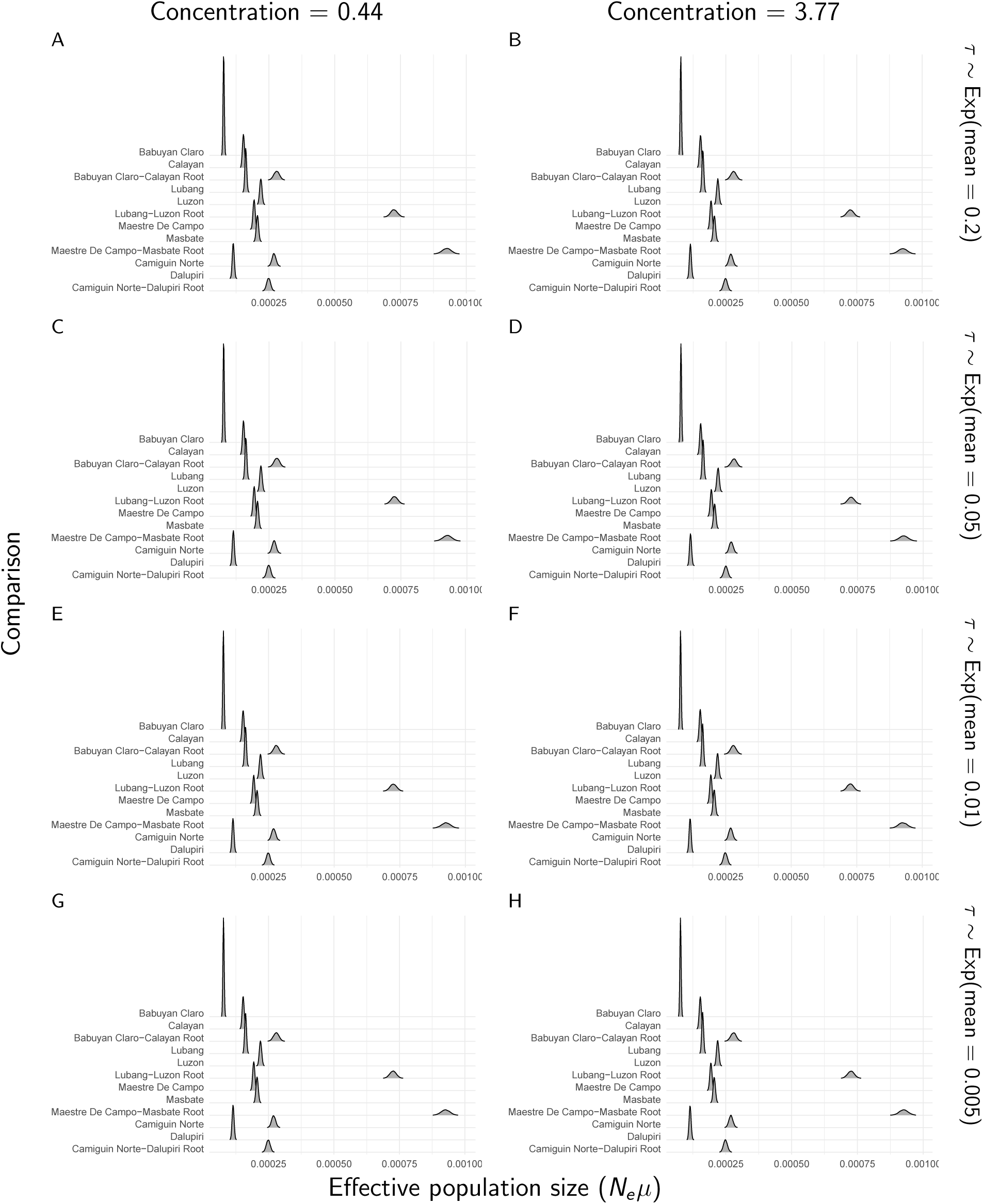
The approximate marginal posterior densities of population sizes for each *Gekko* pair of populations, under eight different combinations of prior on the divergence times (rows) and the concentration parameter of the Dirichlet process (columns). For these analyses, constant characters were included, but all characters with more than two alleles were removed. We generated the plots with ggridges Version 0.4.1 (Wilke, 2018) and ggplot2 Version 2.2.1 (Wickham, 2009).

**Figure S35.**
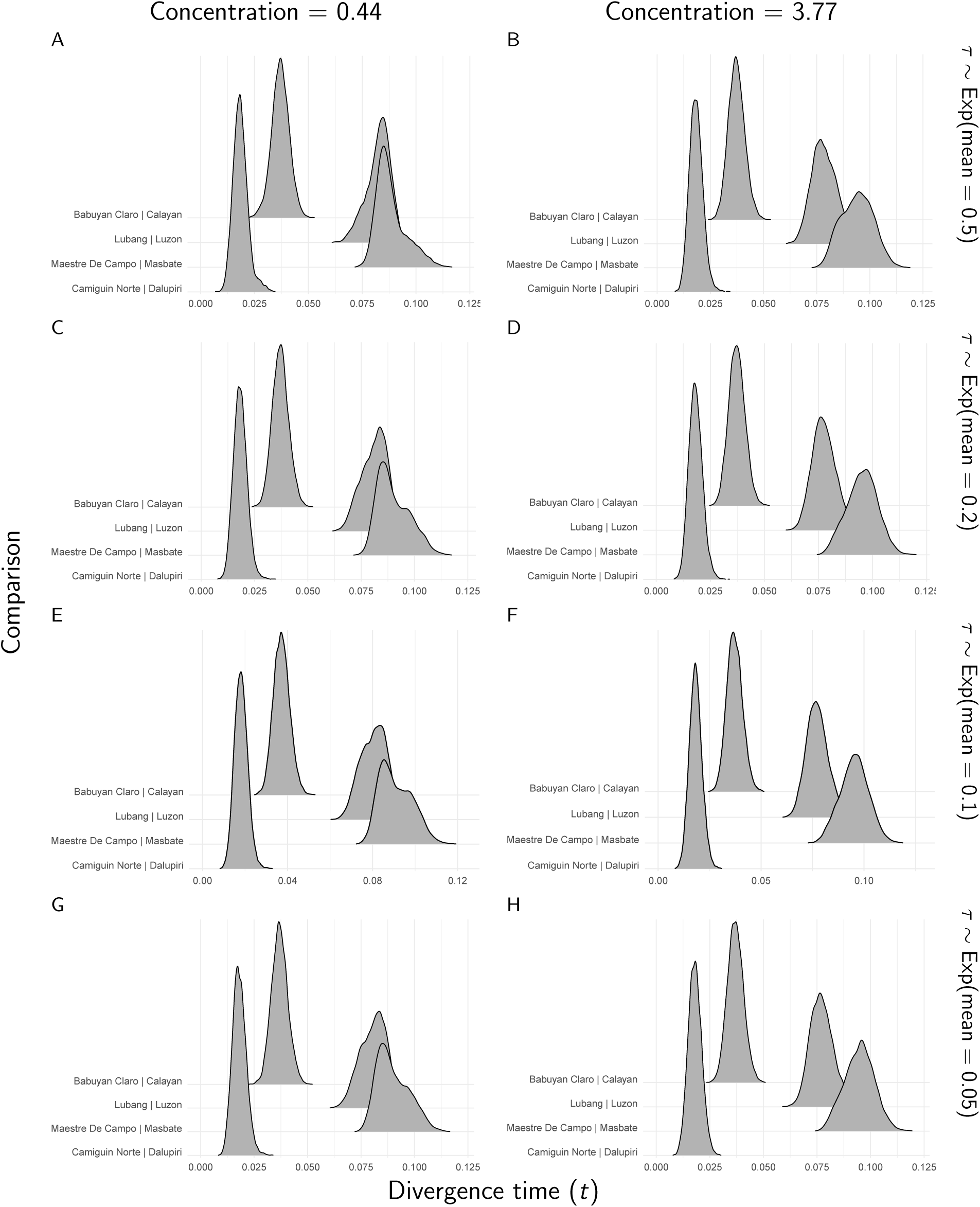
The approximate marginal posterior densities of divergence times for each *Gekko* pair of populations, under eight different combinations of prior on the divergence times (rows) and the concentration parameter of the Dirichlet process (columns). For these analyses, constant characters were excluded, and all characters with more than two alleles were recoded as biallelic. We generated the plots with ggridges Version 0.4.1 (Wilke, 2018) and ggplot2 Version 2.2.1 (Wickham, 2009).

**Figure S36.**
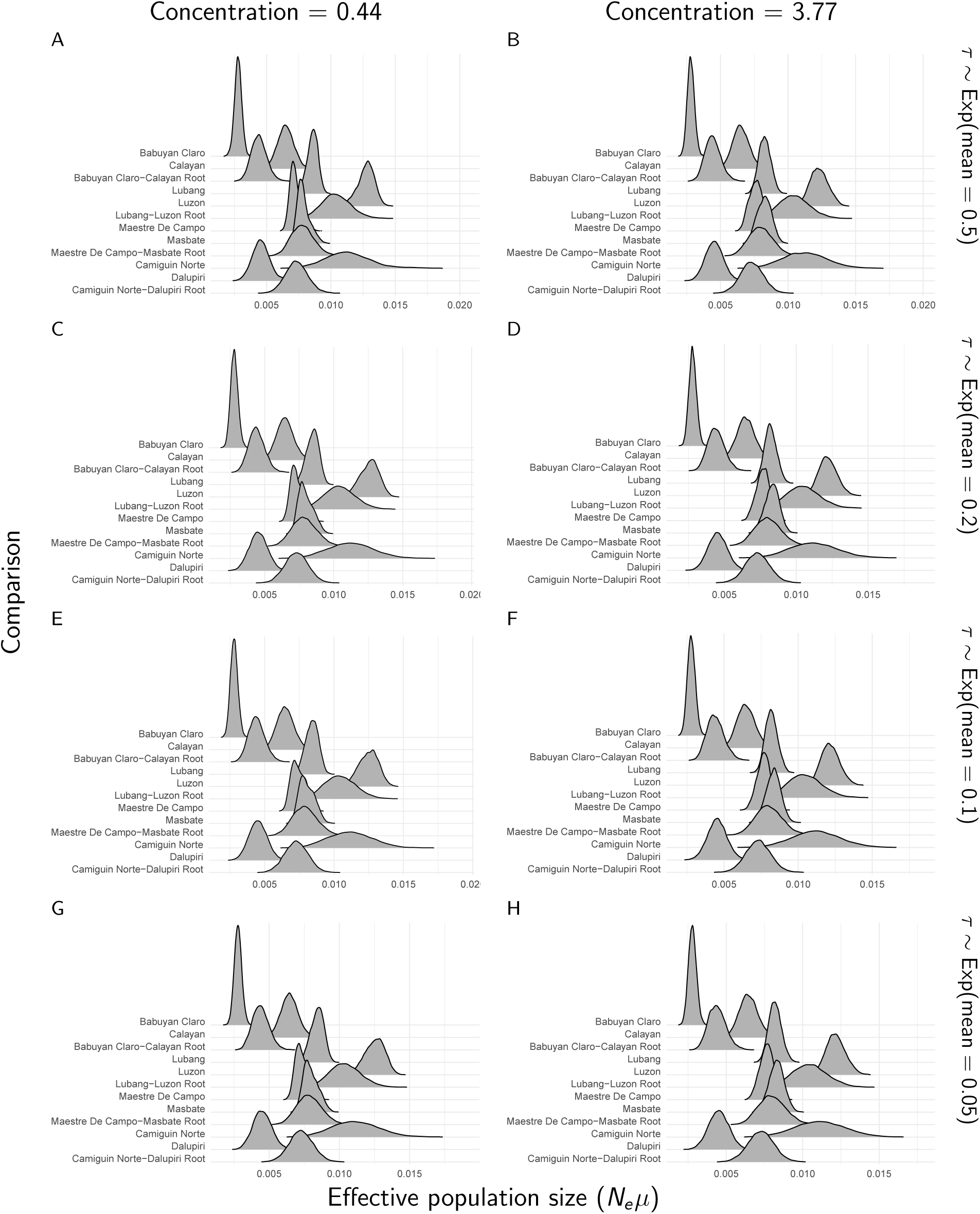
The approximate marginal posterior densities of population sizes for each *Gekko* pair of populations, under eight different combinations of prior on the divergence times (rows) and the concentration parameter of the Dirichlet process (columns). For these analyses, constant characters were excluded, and all characters with more than two alleles were recoded as biallelic. We generated the plots with ggridges Version 0.4.1 (Wilke, 2018) and ggplot2 Version 2.2.1 (Wickham, 2009).

**Figure S37.**
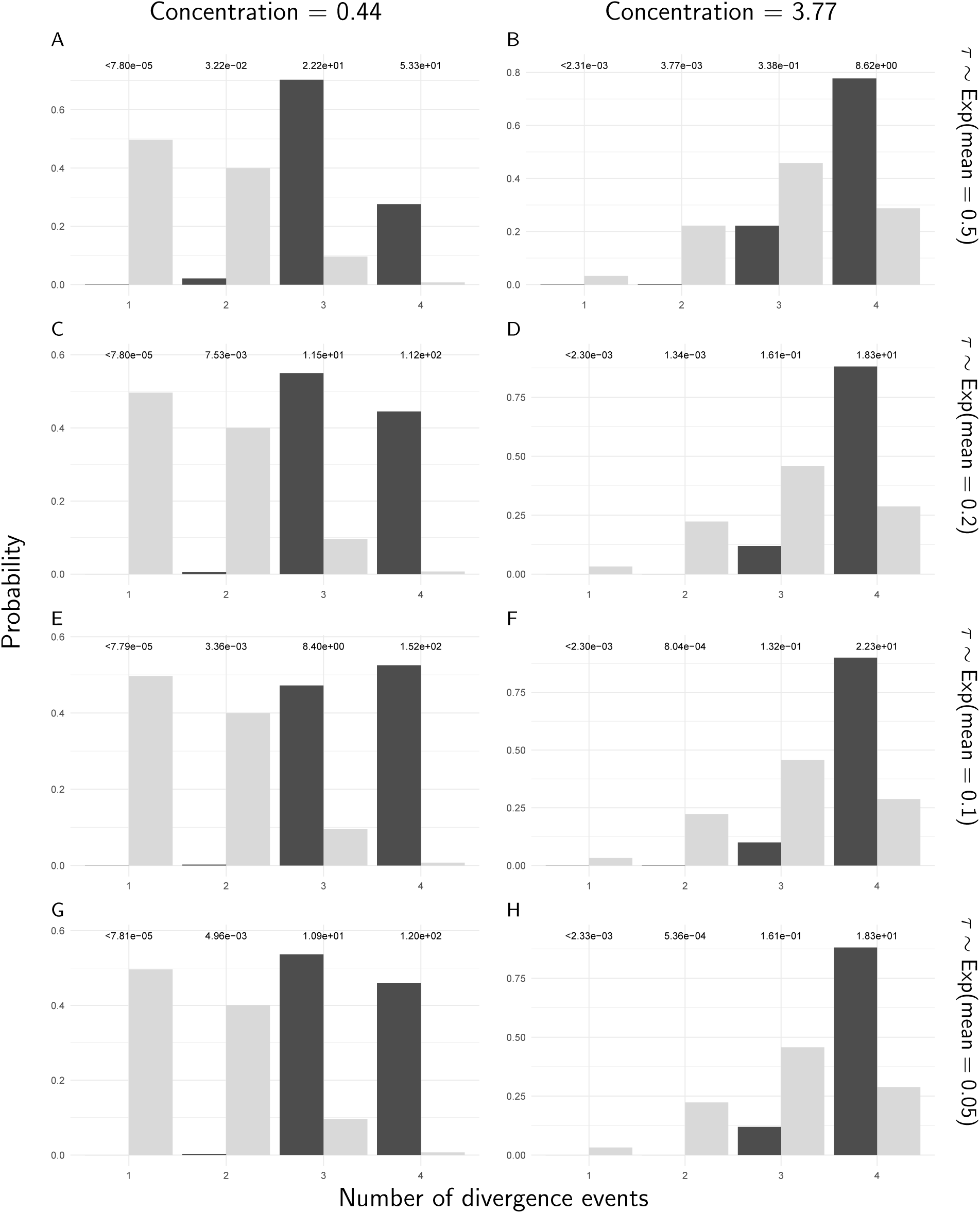
The prior (light bars) and approximated posterior (dark bars) probabilities of the number of divergence events across *Gekko* pairs of populations, under eight different combinations of prior on the divergence times (rows) and the concentration parameter of the Dirichlet process (columns). For these analyses, constant characters were excluded, and all characters with more than two alleles were recoded as biallelic. The Bayes factor for each number of divergence times is given above the corresponding bars. Each Bayes factor compares the corresponding number of events to all other possible numbers of divergence events. We generated the plots with ggplot2 Version 2.2.1 (Wickham, 2009).

**Figure S38.**
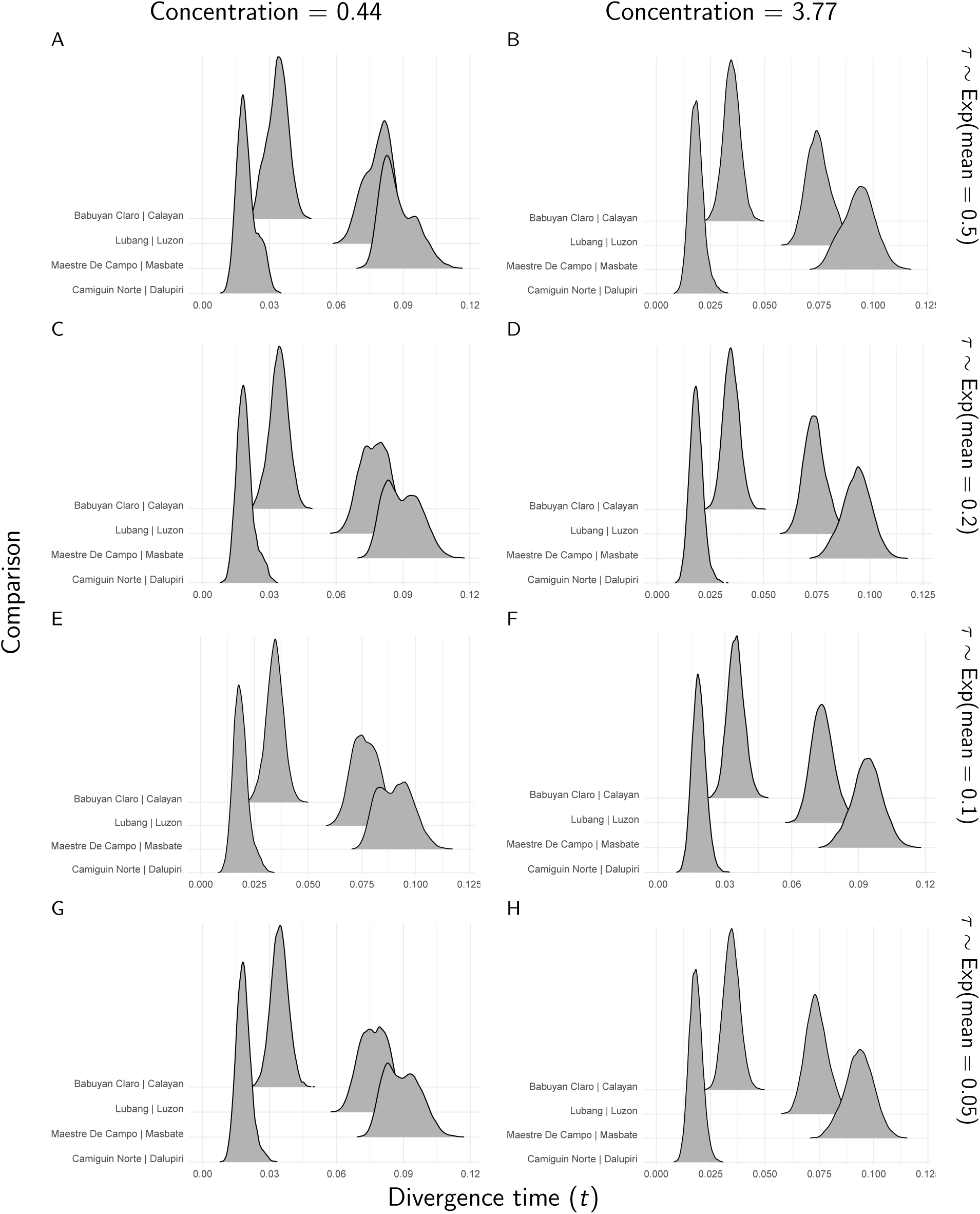
The approximate marginal posterior densities of divergence times for each *Gekko* pair of populations, under eight different combinations of prior on the divergence times (rows) and the concentration parameter of the Dirichlet process (columns). For these analyses, constant Effective population size (*N*_*e*_ *µ*)

**Figure S39.**
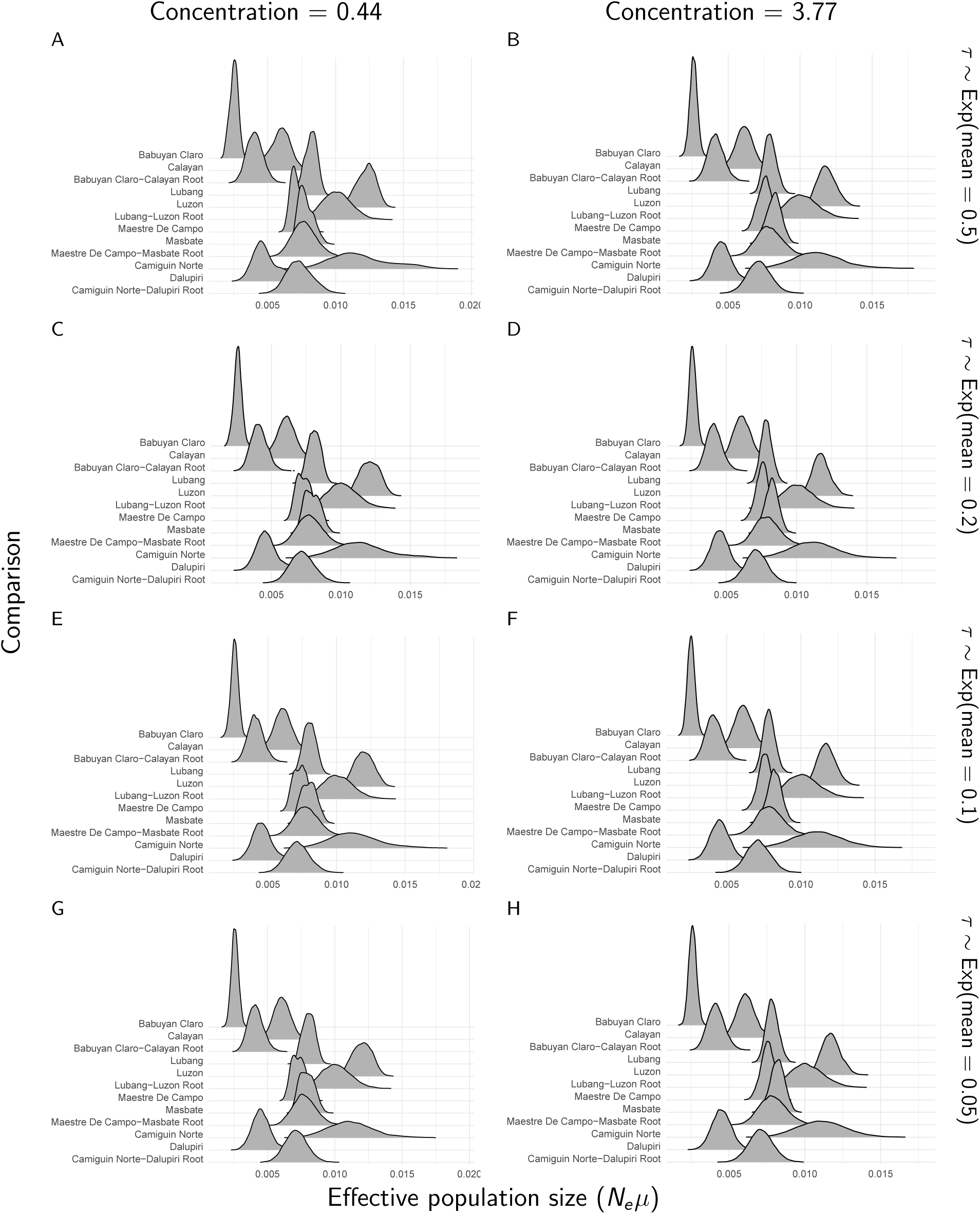
The approximate marginal posterior densities of population sizes for each *Gekko* pair of populations, under eight different combinations of prior on the divergence times (rows) and the concentration parameter of the Dirichlet process (columns). For these analyses, constant characters and all characters with more than two alleles were removed. We generated the plots with ggridges Version 0.4.1 (Wilke, 2018) and ggplot2 Version 2.2.1 (Wickham, 2009).

**Figure S40.**
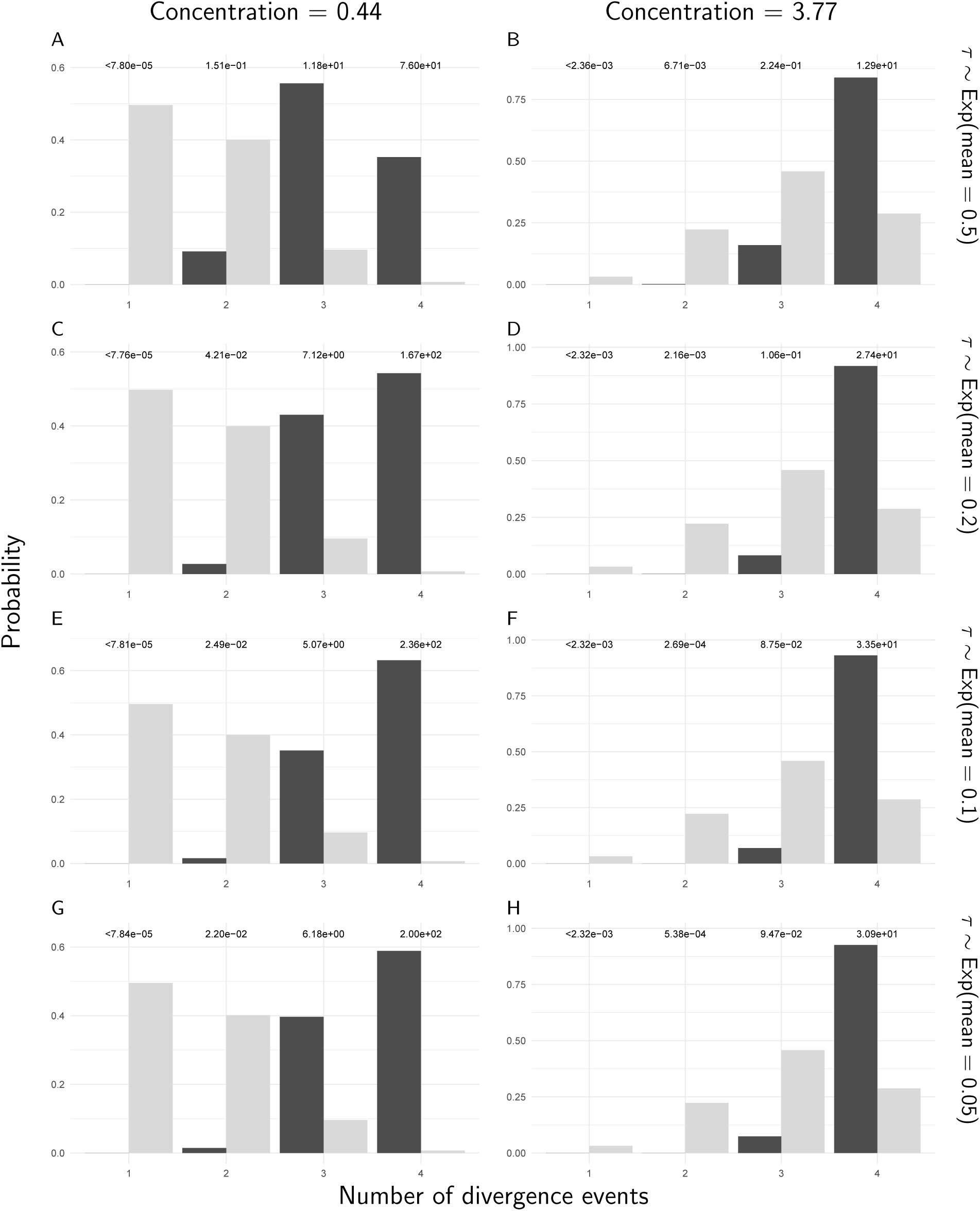
The prior (light bars) and approximated posterior (dark bars) probabilities of the number of divergence events across *Gekko* pairs of populations, under eight different combinations of prior on the divergence times (rows) and the concentration parameter of the Dirichlet process (columns). For these analyses, constant characters and all characters with more than two alleles were removed. The Bayes factor for each number of divergence times is given above the corresponding bars. Each Bayes factor compares the corresponding number of events to all other possible numbers of divergence events. We generated the plots with ggplot2 Version 2.2.1 (Wickham, 2009).

